# A fungal secondary metabolism gene cluster enables mutualist-pathogen transition in root endophyte *Colletotrichum tofieldiae*

**DOI:** 10.1101/2022.07.07.499222

**Authors:** Kei Hiruma, Seishiro Aoki, Yuniar Devi Utami, Masanori Okamoto, Nanami Kawamura, Masami Nakamura, Yoshihiro Ohmori, Ryohei Sugita, Keitaro Tanoi, Toyozo Sato, Wataru Iwasaki, Yusuke Saijo

## Abstract

Plant-associated fungi show diverse lifestyles from pathogenic to mutualistic to the host; however, the principles and mechanisms through which they shift the lifestyles require elucidation. The root fungus *Colletotrichum tofieldiae* (*Ct*) promotes *Arabidopsis thaliana* growth under phosphate limiting conditions. We reveal a *Ct* strain, designated *Ct3*, that severely inhibits plant growth. *Ct3* pathogenesis occurs through activation of host abscisic acid (ABA) pathways via a fungal secondary metabolism gene cluster related to sesquiterpene ABA and botrydial (BOT) biosynthesis. ABA-BOT cluster activation during root infection suppresses host nutrient uptake-related genes and changes the mineral contents, suggesting its role in manipulating host nutrition states. Conversely, disruption or environmental suppression of the cluster renders *Ct3* beneficial for plant growth, in a manner dependent on host phosphate starvation response regulators. Our findings indicate that a fungal metabolism cluster provides a means by which infectious fungi modulate lifestyles along the parasitic–mutualistic continuum in fluctuating environments.

## INTRODUCTION

Plants associate intimately with diverse microbes, including pathogens, nonpathogenic commensals, and beneficial (mutualistic) microbes that promote plant growth. Plant–microbe interactions are typically context-dependent, as these microbes dynamically change their infection modes according to the environment and host conditions (Ewald, 1987; Hiruma *et al*., 2018; Cheng *et al*., 2019; Bailey-Serres *et al*., 2019; Drew *et al*., 2021). Notably, one microbe can switch among pathogenic, nonpathogenic, and beneficial infection modes even in the same host, without changing the genomic sequences (Nongbri *et al*., 2012; Lahrmann *et al*., 2015; Mandyam and Jumpponen, 2014; Hiruma *et al*., 2016; Hacquard *et al*., 2016; Morcillo *et al*., 2020; Drew *et al*., 2021). These findings imply that different *in-planta* lifestyles of plant-associated microbes are continuous within the same plant species and even coexist within the plant individual, and microbes have the ability to refine infection strategies according to the given host environments (Fesel and Zuccaro, 2016; Kia *et al*., 2017). However, the mechanisms by which infectious microbes transit between their contrasting lifestyles remain elusive.

Plants have evolved an elaborate system called phosphate starvation response (PSR) to cope with low inorganic phosphate (Pi) conditions (Chiou and Lin, 2011; Puga *et al*., 2017). Plant PSR has been extensively studied in *Arabidopsis thaliana* and rice models. Under low Pi, plants induce extensive transcriptome reprogramming, mainly through the R2R3-MYB family transcription factors *PHOSPHATE RESPONSE1* (*PHR1*) and related *PHR1-LIKE* (*PHL1*) (Rubio *et al*., 2001; Bustos *et al*., 2010). Plant adaptation to Pi deficit involves activation of genes that promote phosphate absorption, allocation, and usage. Additionally, plants accommodate beneficial fungi that help nutrient acquisition, such as mycorrhizal fungi in the majority of land plants and root-associated endophytic fungus *Colletotrichum tofieldiae* (*Ct*) in nonmycorrhizal *Brassicaceae* plants. Their hyphae grow beyond the Pi depletion zone in the rhizosphere to acquire and transfer phosphorus to the host, providing an extension of the plant root system (Harrison, 2005; Hiruma *et al*., 2016; Almario *et al*., 2017; Choi *et al*., 2018; Frerigmann *et al*., 2021). *PHR1* and *PHL1* positively regulate phosphate transporter genes during *Ct* colonization and are required for *Ct*-mediated plant growth promotion under low Pi (Hiruma *et al*., 2016). *PHR1/PHL1* restricts fungal overgrowth and potential virulence during beneficial interactions with *Ct*, whereas negatively regulating plant immunity against bacteria (Castrillo *et al*., 2017; Tang *et al*., 2022). However, our knowledge on the mechanisms by which *PHR1/PHL1* play varied roles in different plant–microbe associations or host PSR influences microbial lifestyles is limited.

Microbes have evolved an enormous repository of secondary metabolites, including plant hormone mimics for auxin and gibberellic and abscisic acids (ABA). Whereas more than half of the genes are organized in operons in bacteria, functionally related genes are typically distributed across the genome in eukaryotic fungi (Nutzmann *et al*., 2018). However, fungal secondary metabolite biosynthetic genes, as well as their regulatory genes, are often clustered in genetic loci (Hoffmeister and Keller, 2007; Wisecaver and Rokas, 2015; Keller, 2019). Infectious fungal species show high intraspecies variations, compared to non-infectious ones, in the gene contents and genomic structures of secondary metabolism clusters, implying co-evolution with hosts as a driver for the diversification of fungal secondary metabolites (Lind *et al*., 2017). However, at least in the conventional laboratory settings, most of these secondary metabolism genes are not expressed even during host interactions (Brakhage and Schroeckh, 2011; Hacquard *et al*., 2016). Consistently, disruptions of individual genes in secondary metabolism clusters typically do not alter fungal phenotypes *in-planta* (Darma *et al*., 2019). These limitations have hampered the precise determination of the roles for secondary metabolism clusters in plant-infecting fungi.

In the present study, we describe a pathogenic strain, *Ct3*, causing severe plant growth inhibition, in contrast to beneficial strains prevailing in *Ct*. Comparative genomics and functional analyses between these *Ct* strains indicate that, during root colonization, *Ct3* displays transcriptional activation of biosynthesis gene clusters for ABA and the associated sesquiterpene metabolite botrydial (designated ABA-BOT), thereby promoting root infection and pathogenesis. Conversely, *Ct3* becomes beneficial for the plant when this cluster is suppressed. These findings highlight a key role of fungal secondary metabolites in the infection-mode transition of plant-associated fungi.

## RESULTS

### A *Ct* strain severely inhibits plant growth in a nutrient-dependent manner

A *Ct* strain, *Ct61*, isolated from a wild *A. thaliana* population in Spain, promotes plant growth under low Pi conditions by transferring phosphorus to the host (Hiruma *et al*., 2016). In addition to *Ct61*, different *Ct* strains have been isolated from various plant species and geographical locations (Cannon *et al*., 2012; Sato *et al*., 2015; Hacquard *et al*., 2016, Table S1). These *Ct* strains were diverged in culture growth in nutrient-rich media (Figure 1A).

**Figure 1.**
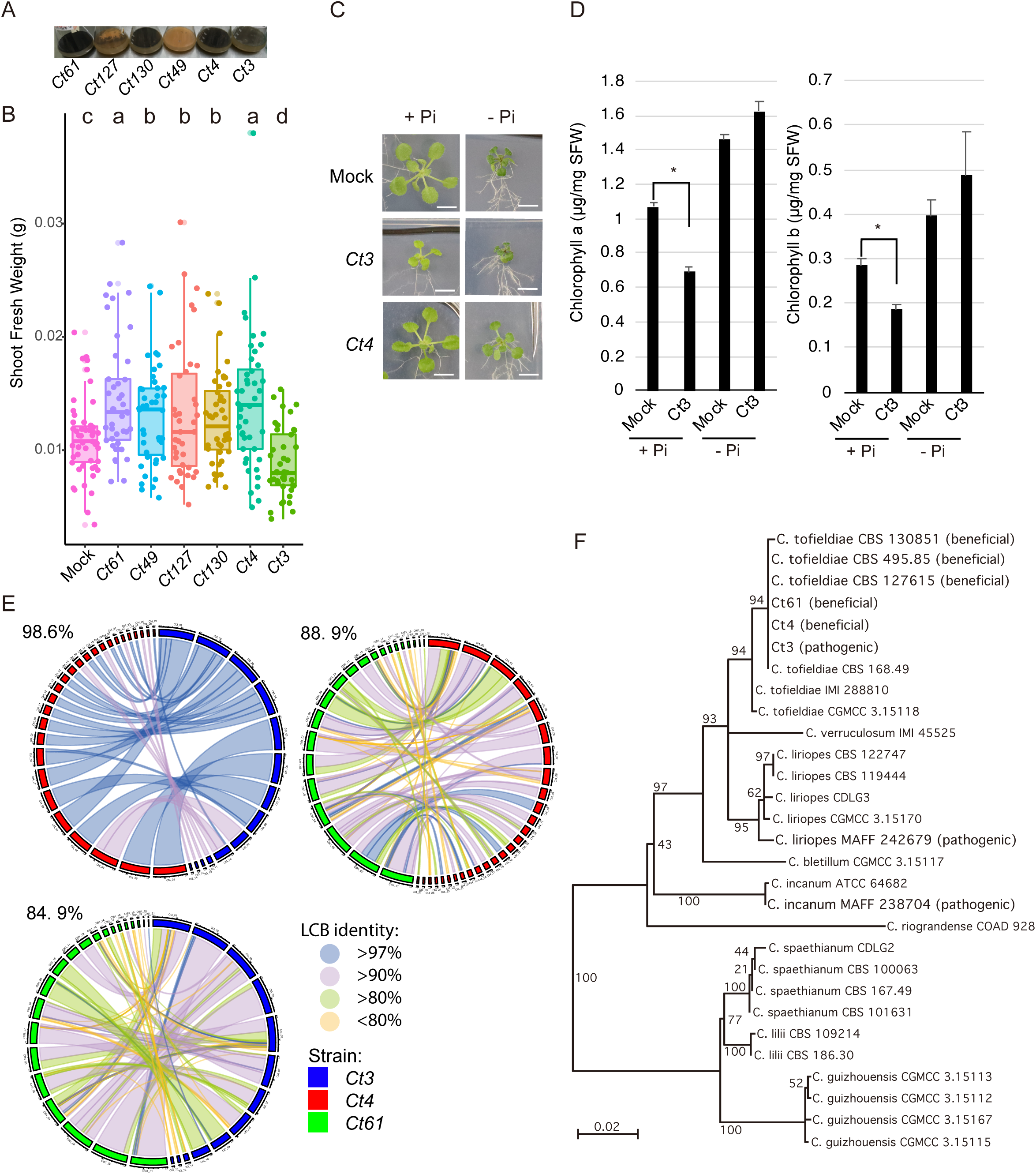
A *Ct* strain severely inhibits plant growth based on nutrient conditions. **a.** Growth of *Ct* strains was identified from the various geographical locations in Mathur’s nutrient media after 10 days of incubation. *Ct61* (*Ct61*) (Spain), *Ct127* (Portugal), *Ct130* (Germany), *Ct49* (Switzerland), *Ct4* (Japan), and *Ct3* (Japan). **b.** Quantitative measurement of *A. thaliana* fresh shoot weighted upon fungal inoculation under low Pi conditions (50 μM KH_2_PO_4_). Plants were incubated for 24 days from germination either with mock-treated, *Ct61*, C*t49*, *Ct127*, *Ct130*, *Ct4*, or *Ct3*, after which fresh weight was measured. Boxplot represents combined results from three independent experiments. Each dot represents individual plant samples. Different letters indicate significantly different statistical groups (ANOVA, Tukey-HSD test). **c.** Pictures of *A. thaliana* plants grown in normal (625 μM KH_2_PO_4_) or low Pi conditions with and without *Ct3* or *Ct4*. The picture was taken after 24-day post-infection (24 dpi) on the same scale (bars = 0.5 cm). **d.** Quantitative data of plant shoot chlorophyll a and b with and without *Ct3* with normal and low Pi. Asterisks indicate significantly different means between mock and *Ct3* treated plants (p < 0.01, two-tailed t-test). **e.** Syntenies of *Ct3*, *Ct4*, and *Ct61* genomes. The length of each contig was described as 0.8 Mb/scale. % in the figure represents the median % of nucleotide similarity against the whole locally collinear block (LCB). **f.** Phylogeny of *Colletotrichum* species inferred from analyzing sequences of three fungal gene markers (GAPDH, ACT, and TUB2). Bootstrap probability is shown on the branches. The lifestyles of some fungi in *A. thaliana* root under low Pi were investigated and are described in the tree (beneficial or pathogenic).

This prompted us to investigate the possible intraspecies variations of *Ct* in plant infection effects and strategies. We compared six *Ct* strains (Table S1) in their inoculation effects on *A. thaliana* Col-0 under gnotobiotic low Pi conditions (50 μM KH_2_PO_4_). Five of the tested *Ct* strains, including *Ct61*, significantly promoted plant growth, indicated by primary root length and shoot fresh weight (SFW), despite slight variations in the shoot growth promotion (Figures 1B and S1A-B). The results suggest that plant growth-promoting (PGP) function under low Pi, at least discernible in *A. thaliana*, is extensively shared by *Ct* strains from various host and geographical niches. Notably, in contrast to these PGP strains, *Ct3* severely inhibited shoot and root growth under low Pi (Figures 1B and S1B). *Ct3* inhibited plant growth in additional 15 *A. thaliana* accessions (Figure S1C), and also in *Brassica rapa* var. perviridis (Komatsuna) (Figure S1E, Left), whose growth was promoted by *Ct4* (a *Ct* strain isolated together with *Ct3,* Figure S1E, Right), Table S1), on unsterilized low nutrient soil. Plant growth inhibition by *Ct3* was weakly correlated with fungal root colonization in the tested accessions (Figures S1C-D, Pearson’s product–moment correlation, SFW fold change (*Ct3*/Mock) vs. fungal biomass, r = -0.5853342, p-value = 0.01722). These results suggest persistence of the pathogenic *Ct3* lifestyle across *Brassicaceae*, at least under the tested conditions.

Since *Ct61* PGP function is specific to low Pi (Hiruma *et al*., 2016), we tested whether phosphate availability influences *Ct3* infection phenotypes. Under normal Pi conditions (625 µM), *Ct3* caused plant growth inhibition and leaf chlorosis (Figures 1C–D and S1F), whereas *Ct4* did not inhibit plant growth (Figure 1C). Under low Pi, *Ct3* compromised plant growth, but did not cause leaf chlorosis or reduce leaf chlorophyll contents (Figures 1C–D). In contrast to *Ct3*, *Ct4* promoted plant growth (Figure 1C). The results suggest that *Ct3* pathogenesis is alleviated under phosphate deficiency, under which *Ct4* promotes plant growth.

To grasp the genomic basis for the intraspecies variations between pathogenic *Ct3* and beneficial *Ct4*, we determined their whole-genome sequences using a long-read generating PacBio sequencing platform. We also resequenced the genome of beneficial *Ct61,* following previous analyses with Illumina and 454 short-read generating platforms (Hacquard *et al*., 2016). PacBio long reads provided high-quality genome assemblies for all strains, ranging from 53 to 55 Mb, with similar gene numbers for candidate effector proteins, carbohydrate-active enzymes, and secondary metabolism clusters (Table S2). Whole-genome alignment performed by Mauve (Darling *et al*., 2004) between *Ct3* and *Ct4* showed a high degree (Median: >98%) of nucleotide identity between the two genomes, despite their contrast infection strategies. In contrast, a comparison between the beneficial strains *Ct61* and *Ct4* showed numerous genomic rearrangements and a lesser degree of nucleotide identity (88.9%), implying that *Ct61* and *Ct4* are quite diverged at the genome level, despite the similarities in host benefits (Figure 1F). These results suggest that nucleotide identity scores largely reflect the geographical distances of their origins. In contrast, molecular phylogenic analysis using three molecular markers currently best for identification of spaethianum complex (Vieira *et al*., 2020) suggests that beneficial *Ct* strains have evolved from the pathogenic relative, such as *Colletotrichum incanum* (Figure 1F). It seems that beneficial lifestyles have evolved from pathogenic ancestors in *Ct* species.

### Pathogenic *Ct3* activates host phosphate starvation responses and ABA signaling pathways during early root infection

We next investigated plant responses following root inoculation with beneficial and pathogenic *Ct* strains via RNA-seq analysis at two time points, 10 dpi (days post-inoculation) and 24 dpi (Figure 2A). At 10 dpi, *Ct3* caused plant growth inhibition under both normal and low Pi conditions, whereas PGP effects of beneficial *Ct* strains were not yet apparent (Figure S2A). At 24 dpi, the degree of plant growth inhibition by *Ct3* was lowered under Pi deficiency compared with Pi sufficiency (Figure S1F), whereas PGP by beneficial *Ct61* and *Ct4* became discernible only under low Pi (Figures 1B and S2B). In contrast to these *Ct* strains, KHC strain (previously classified as *C. tofieldiae*), closely related to *C. higginsianum* (Figure 1F), colonized the roots without significantly altering plant growth under either nutrient condition (Figures 2A and S2B), displaying a commensal lifestyle.

**Figure 2.**
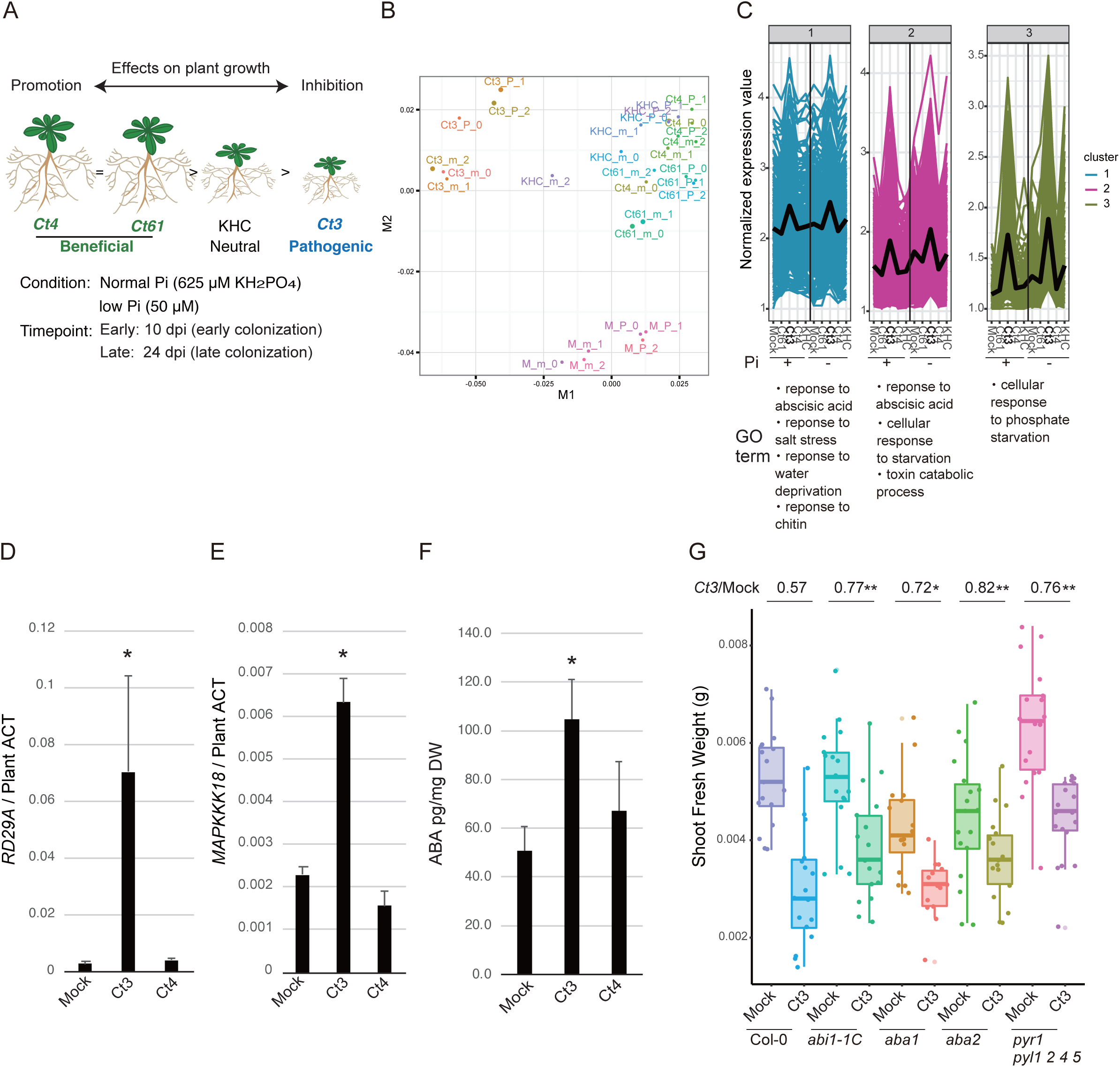
During early root colonization, pathogenic *Ct3* activates host ABA and phosphate starvation response pathways. **a.** Experimental setup for dual RNA-seq analysis of *A. thaliana* and *Ct* transcriptomes. **b.** Multidimensional scaling (MDS) analysis chart generated from plant transcriptome profile during association with mock (M), *Ct61*, *Ct4*, *Ct3*, or KHC in normal (P) or low Pi (m). Three biological replicates are shown in the chart. **c.** Expression patterns of *A. thaliana* genes differentially expressed during pathogenic *Ct3* compared with other beneficial *Ct* strains treatment ((|log2FC|>1, FDR < 0.05) using cluster analysis divided by k-mean (K = 3, PAM clustering with Jensen-Shannon distance). Genes listed in each cluster were subjected to Go analysis (The full list is available in Table S4). **d.**, **e.** Expression of ABA-responsive genes (*RD29A* and *MAPKKK18*) during treatment of mock, *Ct3*, and *Ct4* at 10 dpi under low Pi. Asterisks indicate significantly different means between Mock- and *Ct3-*treated plants (p < 0.05, two-tailed t-test). **f.** Quantitative measurement of *A. thaliana* root ABA upon fungal inoculation under low Pi. Asterisks indicate significantly different means between Mock-and *Ct3*-treated plants (SD, p < 0.05, two tailed t-test). **g.** Quantitative measurement of *A. thaliana* fresh shoot weight upon fungal inoculation. Shoot fresh weights of Col-0, *abi1-1C*, *aba1*, *aba2*, *pyr1-1 pyl1 pyl2 pyl4 pyl5* plants either with mock or *Ct3* for 24 dpi under low Pi conditions were measured. Asterisks indicate significantly different fresh weight ratio (*Ct3*/Mock) between Col-0 and each mutant (* p < 0.1, ** p < 0.05, two-tailed t-test).

We first focused on the divergence in plant responses between these *Ct* strains at 10 dpi. Multidimensional scaling analysis indicated a clear separation of plant transcriptome in response to *Ct3* compared to beneficial *Ct* and commensal KHC strains, suggesting distinct plant responses to *Ct3* at this stage (Figure 2B). The results revealed 758 *A. thaliana* genes that were differentially upregulated in *Ct3*-infected roots compared with beneficial *Ct* strains, under normal or low Pi conditions (log_2_FC > 1, FDR < 0.05, Table S3). K-means clustering classified these differentially expressed genes (DEGs) into three different clusters (Figure 2C and Table S4). In Clusters 1 and 2, Gene Ontology (GO) categories related to ABA signaling (“response to abscisic acid, FDR = 0.0025 [Cluster 1] and 0.0073 [Cluster 2],” “response to salt stress, FDR = 0.0023 [Cluster 1] and 0.0025 [Cluster 2] ”, “response to water deprivation, FDR = 0.0031 [Cluster1])” were overrepresented (Figure 2C and Table S4), suggesting that *Ct3* specifically induces host ABA responses. ABA promotes plant adaptation to water deficit (Umezawa *et al*., 2010), and also influences immunity or susceptibility-related pathways (de Torres-Zabala *et al*., 2007; Mine *et al*., 2017; Peng *et al*., 2019; Huai *et al*., 2019; Roussin-Léveillée *et al*., 2022; Hu *et al*., 2022). qRT-PCR analyses validated increased expression of ABA-responsive plant genes, *AtRD29A* (Yamaguchi-Shinozaki and Shinozaki, 1993) and *AtMAPKKK18* (Okamoto *et al*., 2013) during pathogenic *Ct3* infection (Figures 2D-E). ABA accumulation also increased in roots following *Ct3,* but not *Ct4,* colonization (Figure 2F). The results indicate that *Ct3*, but not *Ct4*, strongly induces ABA responses in roots. We then tested whether and if so how host ABA contributes to *Ct3*-mediated plant growth inhibition, with plant mutants disrupted in ABA signaling, *abi1-1C* (Umezawa *et al*., 2009); ABA biosynthesis, *aba1* (Morris *et al*., 2006) and *aba2-12* (Gonzalez-Guzman *et al*., 2002); and ABA perception, *pyr1-1 pyl1 pyl2 pyl4 pyl5* (Mega *et al*., 2019). *Ct3*-mediated plant growth inhibition was alleviated in these ABA-defective mutants under low Pi (Figure 2G). In contrast, plant growth promotion via beneficial *Ct4* under low Pi was not affected in *abi1-1C* (Figure S2C). These analyses suggest that pathogenic but not beneficial lifestyles of *Ct3* are dependent on the core ABA biosynthetic and response pathways of the host.

During beneficial interactions under low Pi, *Ct61* induced a subset of PSR-related genes in plants, including phosphate transporter *PHT* genes, at 24 dpi (Hiruma *et al*., 2016). Since PSR-related transcriptional reprogramming largely relies on *PHR1* and *PHL1* (Bustos *et al*., 2009), we next assessed the possible impact of beneficial *Ct* strains on *PHR1/PHL1*-regulated PSR genes under low Pi at 24 dpi. Cross-referencing our data with previously described, 193 *PHR1/PHL1* regulons (Bustos *et al*., 2009), indicated increased expression for approximately half of them [88 of 193 genes (*Ct61*), or 99 of 193 genes (*Ct4*), FDR < 0.05] during beneficial *Ct* interactions compared with the mock controls, specifically under low Pi (Figure S2D). In contrast, PSR gene activation was not increased during pathogenic or commensal interaction with *Ct3* or KHC, respectively, at 24 dpi (Figure S2D). At 10 dpi however, *Ct3* strongly upregulated a subset of PSR-related genes under low Pi compared with the mock controls (Figure S2E, 146 of 193 genes, FDR < 0.05), indicated by the induction of Clusters 2 and 3 genes related to PSRs, in which GOs overrepresented “Cellular response to starvation (0.00142, Cluster 2)” and “cellular response to phosphate starvation (0.0347, Cluster 3)” (Figure 2C and Table S4). Upregulation of PSR genes was associated with an increase in shoot P concentrations following *Ct3* inoculation under low Pi (Figure S2F), coincident with alleviated pathogenesis. Notably, *Ct3* inoculation upregulated a subset of *PHR1/PHL1* regulons, which were otherwise not induced, under Pi sufficient conditions (Figures 2C (Cluster 3) and S2E, 69 of 193 genes, FDR < 0.05). The results suggested that *Ct3* infection results in *PHR1/PHL1* regulon activation, at least during an early infection phase, despite the eventual negative effects on plant growth.

We then examined how *PHR1/PHL1*-dependent PSR influences pathogenic *Ct3* lifestyles, by testing for *Ct3* inoculation phenotypes in *phr1 phl1* plants. Compared to WT, *phr1 phl1* displayed severe growth inhibition following *Ct3* inoculation under low Pi (Figure S2G). The results indicate a critical role for *PHR1/PHL1* in the alleviation of plant growth inhibition under low Pi. Together with the transcriptome analyses above (Figure S2), we infer from the results that activation of *PHR1/PHL1* regulons serves to restrict *Ct3* pathogenesis.

### Genetic disruption of fungal ABA and BOT biosynthesis genes leads to switching from pathogenic to beneficial lifestyles in *Ct3*

To assess the significance for transcriptional activation of host ABA responses, we examined fungal transcriptome profiles during pathogenic *Ct3* infection (Table S5). As *Ct3*-mediated ABA response activation and plant growth inhibition were pronounced at 10 dpi (Figures 2C and S1F), we assembled *Ct3* genes that were differentially upregulated at 10 dpi compared to 24 dpi. This resulted in 304 fungal genes upregulated in an early phase under normal or low Pi conditions (Figure 3A, log_2_FC > 1, FDR < 0.05). The number of these genes was greater under normal Pi than low Pi (Figure 3A), consistent with enhanced *Ct3* impacts on plant growth (Figure 1).

**Figure 3.**
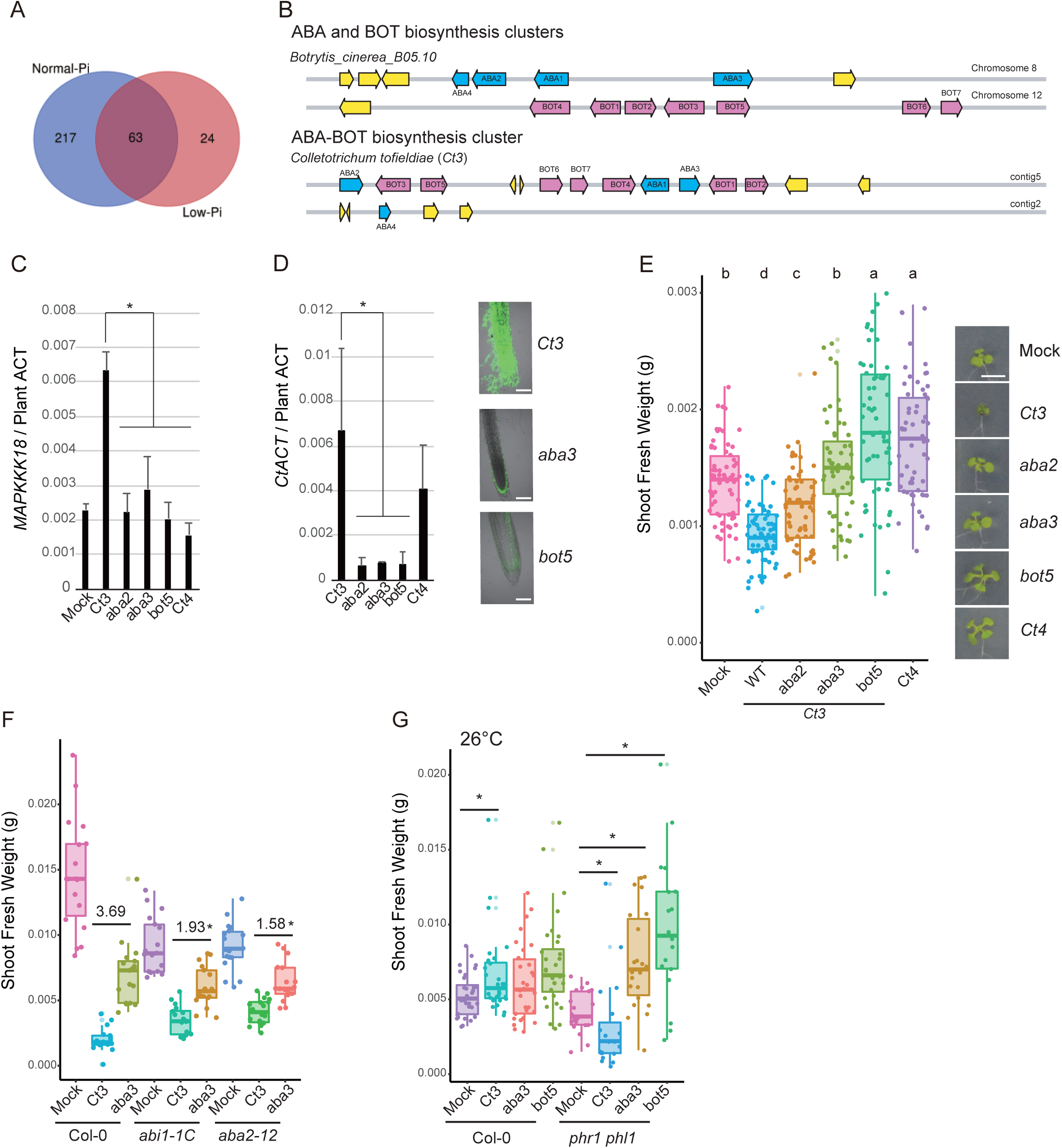
Disruption of the fungal ABA and BOT biosynthesis genes switches *Ct* lifestyle from pathogenic to beneficial. **a.** Venn diagram showing *Ct3* genes differentially upregulated in early colonization (10 dpi) compared with late colonization (24 dpi) (|log2FC|>1, FDR < 0.05, Table S5). Sixty-three *Ct3* genes were significantly upregulated during early root colonization by *Ct3* in both normal and low Pi. ABA and BOT biosynthesis genes were included in the venn diagram. **b.** Overview of *Ct3* ABA-BOT biosynthesis gene cluster. ABA and BOT biosynthesis gene clusters from *B. cinerea* were described for comparison. **c.** Expression of *MAPKKK18* during treatment of mock, *Ct3*, *Ct3aba2*, *Ct3aba3*, *Ct3bot5*, and *Ct4* at 10 dpi under low Pi. Asterisks indicate significantly different means between *Ct3* and other fungal-treated plants (n = 3, p < 0.05, two-tailed t-test). **d.** Fungal biomass measured by *Colletotrichum ACTIN* relative abundance in *A. thaliana* roots at 10 dpi under low Pi. Asterisks indicate significantly different means between *Ct3* and other fungal (*aba2*, *aba3*, *bot5*) treated plants (n = 3, p < 0.05, two-tailed t-test). The photos represent root colonization by *Ct3*, *aba3*, and *bot5*. *Ct* hyphae at 16 dpi and were stained by WGA-Lectin. **e.** Quantitative measurement of *A. thaliana* fresh shoot weighted upon fungal inoculation under low Pi. Plants were incubated for 14 days from the germinations with mock-treated, *Ct3* (WT), *aba2*, *aba3, bot5*, and *Ct4* with fresh weight measured. Different letters indicate significantly different statistical groups (ANOVA, Tukey-HSD test). The photos show the representative treated-plants (Bars = 1 cm). **f.** Quantitative measurement of *A. thaliana* fresh shoot weighted upon fungal inoculation under normal Pi. *aba3*/*Ct3* ratio is described in each genotype. Asterisks indicate significantly different between Col-0 and mutant plants (either *abi1-1C* or *aba2*, p < 0.05, two-tailed t-test). **g.** Quantitative measurement of *A. thaliana* fresh shoot weighted upon fungal inoculation at 26 °C under low Pi. Asterisks indicate significantly different mean (p < 0.05, two-tailed t-test).

We noticed that ABA biosynthesis genes (*ABA1*, *ABA2*, and *ABA3*) were included in the 304 early-inducible genes. They show high sequence similarity to ABA biosynthesis genes (*bcABA1*–*bcABA3*) in the gray mold pathogen *Botrytis cinerea*. *bcABA1* and *bcABA2* encode cytochrome P450, and *bcABA3* encodes sesquiterpene synthase catalyzing the initial step from farnesyl diphosphate, in a fungal ABA biosynthesis pathway that is entirely different from that of plants (Siewers *et al*., 2006; Endo *et al*., 2014; Takino *et al*., 2018). The three *Ct3* orthologues to fungal ABA biosynthesis genes, all clustered in one genomic region, were upregulated during an early, but not late, phase of root colonization (Table S6). By contrast, their expression was not detected in *Ct4* or *Ct61* during beneficial interactions (Table S6), despite high conservation of their cluster across these *Ct* genomes, including nearly identical coding and regulatory DNA sequences for all three genes (particularly in *Ct4*; Figures 3B and S3N). Notably, *ABA3* expression was also detected in *C. incanum* during root infection (Figure S3A). The results suggested that the mobilization of the fungal ABA biosynthesis gene cluster is specific to pathogenic lifestyles.

This prompted us to test whether fungal ABA biosynthesis genes are required for *Ct3* pathogenesis. We generated several lines of *Ct3* knockout mutants for *ABA2* and *ABA3* using homologous recombination. *aba2* and *aba3* mutant fungi showed WT-like growth and spore formation on nutrient-rich and low Pi media (Figures S3B–C) and WT-like hyphal growth on glass slides (Figure S3D), indicating that these genes are dispensable for culture growth in *Ct3*. We examined a possible role for these fungal genes in the induction of host ABA responses in root inoculation. Compared to WT fungi, ABA-dependent *AtMAPKKK18* induction was reduced in the roots inoculated with *aba2* and *aba3* fungi, suggesting that the fungal ABA biosynthesis genes mediate host ABA responses (Figure 3C). Furthermore, root fungal biomass was dramatically reduced for *aba2* and *aba3* fungi compared with WT fungi (Figures 3D and S3E), indicating that these fungal genes contribute to root infection of *Ct3*.

Plant growth inhibition was also reduced when inoculated with *aba2* and *aba3* fungi compared with WT fungi (Figures 3E and S3F–G). *Ct3* virulence was however restored when ABA was exogenously applied (Figure S3H). Fungal *ABA3*-mediated plant growth inhibition was reduced in host mutants defective in ABA signaling or biosynthesis (Figure 3F). The results suggest that *Ct3* ABA biosynthesis involves and requires the plant ABA core pathway in fungal pathogenesis.

Interestingly, in *Ct3*, these ABA biosynthesis genes are clustered together with those highly related to the biosynthesis of botrydial (BOT), an ABA-related sesquiterpene metabolite promoting virulence in *B. cinerea* (Siewers *et al*., 2005). Five BOT biosynthesis genes (named *BOT1*–*BOT5*) were co-activated with the three ABA biosynthesis genes during root colonization in *Ct3* but not in other beneficial *Ct* strains (Table S6). *BOT5* was also co-activated with *ABA3* in *C. incanum* during root infection (Figure S3A). Therefore, co-activation of ABA and BOT biosynthesis genes was specifically associated with pathogenic lifestyles in the root-infecting *Colletotrichum* species.

We tested whether BOT biosynthesis genes are also required for pathogenesis in *Ct3*. Similar to *aba2* and *aba3* (Figures S3B-C), disruption of three *BOT* genes (*bot1*, *bot3*, and *bot5*) did not alter fungal growth or spore formation in normal nutrient media (Figures S3B-C). However, *bot* and *aba* fungi did not induce *AtMAPKKK18* in the host roots (Figure 3C), suggesting a role for *BOT* genes in mobilizing plant ABA responses. Furthermore, root infection was reduced in *bot* compared with WT fungi (Figures 3D and S3I). Remarkably, *bot5* fungi did not inhibit, but rather promoted plant growth under low Pi, reminiscent of *Ct4* (Figures 3E and S3J). The results suggest that the loss of BOT biosynthesis disables *Ct3* virulence, and even renders the fungus beneficial for the host.

Increasing temperatures (from 22°C to 26°C) favored shoot growth in *A. thaliana*, and even *Ct3* promoted plant growth at 26°C under low Pi (Figure 3G). This was accompanied by a decrease in fungal root colonization at 26°C compared to 22°C (Figure S3K). Plant growth promotion and fungal colonization at high temperatures were unaffected when inoculated with *aba3* or *bot5* fungi, consistent with a great decrease in *Ct3 ABA3* and *BOT5* expression during root colonization at 26°C (Figures 3G and S3L–M). Limitation of *Ct3* root colonization at high temperatures was thus associated with low expression of the fungal ABA-BOT cluster genes. However, in *phr1phl1* plants, *Ct3* inhibited plant growth even at 26°C in a manner dependent on the ABA-BOT cluster (Figure 3G), suggesting that the suppression of fungal ABA-BOT cluster and pathogenesis at high temperatures requires the host *PHR1/PHL1*. Notably, *aba3* and *bot5* fungi both promoted plant growth even in *phr1 phl1* plants at 26°C (Figure 3G), suggesting that *PHR1/PHL1* are not required for PGP *per se* at high temperatures when fungal ABA-BOT cluster is disrupted. Consistently, in the absence of fungal inoculation, WT and *phr1 phl1* plants were indistinguishable in shoot growth at 26°C even under low Pi, at least in our settings (Figure 3G). These results indicate a critical role for ABA-BOT cluster and its negative regulation by host *PHR1/PHL1,* in the pathogen-mutualist transition of *Ct3*.

### ABA and BOT biosynthesis gene clusters are distributed across plant-associated fungi, possibly along with fungal virulence evolution

Our comparative genomic analysis indicated that ABA and BOT biosynthesis genes are conserved as a single cluster in beneficial and pathogenic *Ct* genomes but are separate in the *B. cinerea* genome (Figure 4A). Furthermore, these genes, in particular the BOT cluster genes, show very high degrees of sequence conservation between *Colletotrichum* spp. (class Sordariomycetes) and *B. cinerea* (class Leotiomycetes). Such sequence conservation was unexpected given the considerable phylogenetic distance between the two fungal lineages, which diverged approximately 261.6 million years ago (Hacquard *et al*., 2016).

**Figure 4.**
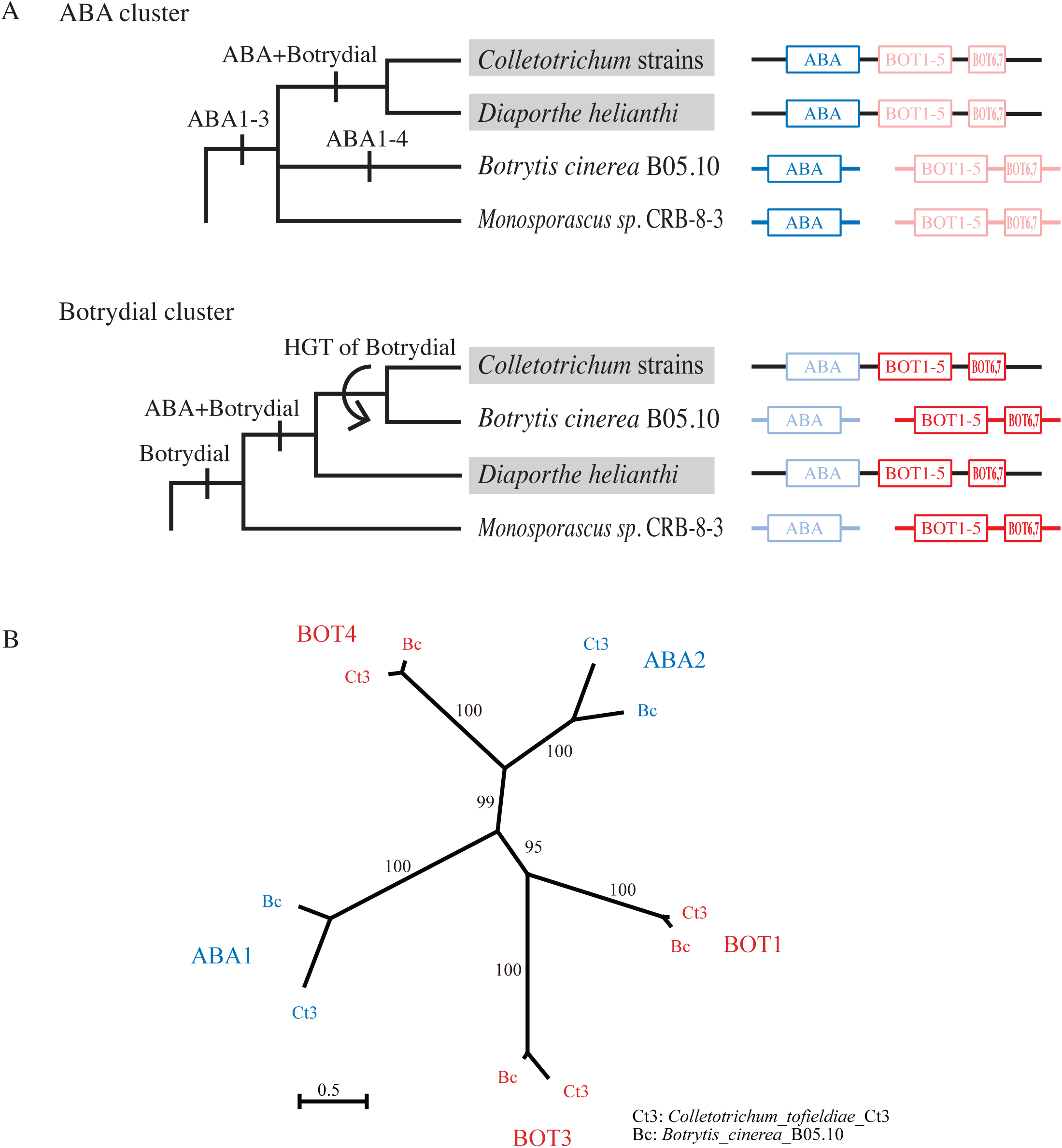
Evolution of ABA and BOT biosynthesis genes. **a.** Phylogenetic relationship and genetic structure of ABA and BOT gene clusters. All four genera with ABA and BOT gene clusters in their genome were used for OTU. Organisms harboring adjacent ABA and BOT clusters (ABA + BOT (ABA-BOT) clusters) in their genome are labeled with gray backgrounds. Phylogeny of each gene in the clusters is shown in Figures S4A–K. **b.** Maximum Likelihood (ML) tree of ABA and BOT genes in Cytochrome P450 family using IQ-TREE version 1.6.11. *Ct3*: *C. tofieldiae Ct3*; Bc: *B. cinerea* B05.10. The phylogenetic relationship of the ABA and BOT genes and other 409 P450 genes in *C. tofieldiae*, *B. cinerea*, and *H. sapiens* is indicated in Figure S4N. Bootstrap probability is shown on the branches. Scale bar represents substitutions per site.

To assess the evolutionary origin(s) of ABA and BOT biosynthesis genes, we generated molecular phylogenetic trees for these genes, from the five *Colletotrichum* strains (*Ct3, Ct4, C. liriopes*, *C. spaethianum* [Utami and Hiruma, 2022], all four belonging to the *spaethianum* clade, and KHC strain, belonging to *destructivum* clade), and from the GenBank non-redundant protein sequences (NR) database (February/March, 2020; Figure 4, Tables S7-8). Phylogenetic analyses showed that ABA and BOT cluster genes were distributed across distantly related plant-associated fungi, in a manner different from their phylogenic relationships (Figures S4A–L), (2) that ABA and BOT cluster genes each formed monophyletic groups (Figures S4A-L), (3) that genes in ABA or BOT cluster each have similar evolutionary histories (Figures S4A–C, S4E–M), but (4) that the ABA and BOT clusters have different evolutionary histories, indicated by their divergence in molecular phylogenetic trees (Figures 4A and S4A–L). Those substantial differences between species and gene trees suggest horizontal gene transfers (HGTs) distributing the ABA and BOT biosynthesis gene clusters among these plant-associated fungi. Our comparative genomic analysis revealed that *Diaporthe helianthi*, a plant pathogen, also possesses a single ABA-BOT biosynthesis gene cluster of high synteny when compared with *Ct* (Figure S4M). This suggests a common evolutionary origin for these ABA-BOT clusters, which predated HGT of the BOT cluster to *Botrytis* spp (Figures 4A and S4L–M). Given that the ABA/BOT clusters have likely arisen multiple times in different phytopathogenic fungi, and that ABA and BOT biosynthesis genes are both required for *Ct3* pathogenesis (Figure 3), it is conceivable that the acquisition of ABA and BOT biosynthesis gene cluster(s) contributes to pathogenesis evolution in plant-associated fungi.

Gene composition in ABA and BOT biosynthesis clusters was slightly different across species. The ABA biosynthesis cluster consists of *ABA1*–*ABA3* in e.g., *Ct*, *C. incanum*, *C. liriopes,* and *D. helianthi* or *ABA1*-*ABA4* in e.g., *B. cinerea* (Figures 4, S4A–D, and S4L–M). *ABA4* was separately located in *Ct* genomes (Figure S4M), and unlike *ABA1*–*ABA3,* its expression was not detected in *Ct3* root colonization (Table S6), suggesting that *ABA4* is dispensable for *Ct3* root colonization or pathogenesis. The BOT gene cluster typically consists of *BOT1*–*BOT7*, where *BOT6* and *BOT7* correspond to Zn_clus (Zn_2_Cys_6_ transcription factor) and Adh_short (retinol dehydrogenase 8) genes, respectively, named in *B. cinerea* (Porquier *et al*., 2016). *BOT6* and *BOT7* show similar phylogenetic distributions with the other five BOT genes within *Colletotrichum* and *Botrytis* strains, indicating functional coupling of these genes in these fungal lineages (Figures S4E–M). We confirmed that *BOT6* and *BOT7* were strongly expressed during *Ct3* pathogenesis (Table S6).

### Host nitrogen and iron uptake genes are targeted by fungal ABA-BOT cluster

*ABA1, ABA2, BOT1, BOT3*, and *BOT4*, all annotated as P450 genes, constitute a large monophyletic group (Figures 4B and S4N). Co-expression of these genes and their requirements for activating the plant core ABA pathway during *Ct3* infection implied the existence of a common host target(s) for the metabolites generated from the two fungal biosynthesis pathways. To gain insight into the host pathways affected by the fungal ABA-BOT cluster, we assembled *A. thaliana* genes specifically induced or repressed by pathogenic *Ct3*, but not beneficial *Ct4*, in a manner dependent on the *Ct3* ABA-BOT cluster. We examined root transcriptome at 10 dpi with WT, *aba2*, *aba3,* and *bot5* of *Ct3*, and WT *Ct4* under low Pi, where the *ABA/BOT* expression status greatly influences *Ct3* lifestyles (Table S9). Plant responses to *Ct3 aba* and *bot* fungi were similar to *Ct4* responses (Figure S5A), consistent with their PGP effects under low Pi (Figure 3). Plant DEGs following inoculation with WT *Ct3*, compared with *aba2*, *aba3*, and *bot5* mutants of *Ct3* and WT *Ct4*, included 288 *Ct3-*induced genes and 375 *Ct3-*repressed genes (Figures 5A–B and Tables S10–11). *Ct3-*induced genes were overrepresented with the genes responsive to oxidative stress, chitin, ABA, and cellular response to phosphate starvation. DNA motif analysis revealed DNA sequences bound by NAC transcription factors related to drought tolerance (Sakuraba *et al*., 2015) being enriched within 1000 bp upstream of their transcription initiation sites (Figure S5B). Notably, the RNA-seq analyses also indicated that PSR gene activation at 10 dpi with *Ct3* was dependent on the fungal ABA-BOT cluster. We further showed that expression of *AT4*, a PHR1-regulon during PSR (Bustos *et al*., 2010), was significantly lowered in *abi1-1C* and *aba2* plants when colonized with *Ct3* (Figure S5C), indicating a role for the host ABA pathway in *Ct3* induction of host PSR genes. Interestingly, a large subset of GOs overrepresented in the 375 *Ct3-*repressed plant genes were related to hosting nutrition, such as inorganic anion transport, cation homeostasis, and inorganic ion homeostasis (Table S11 and Figure 5D). Consistently, *Ct3* root colonization changed host shoot nutrition status depending on ABA-BOT (Figure S5D). The results suggest that *Ct3* root infection suppresses host nutrient uptake and impacts host mineral homeostasis.

**Figure 5.**
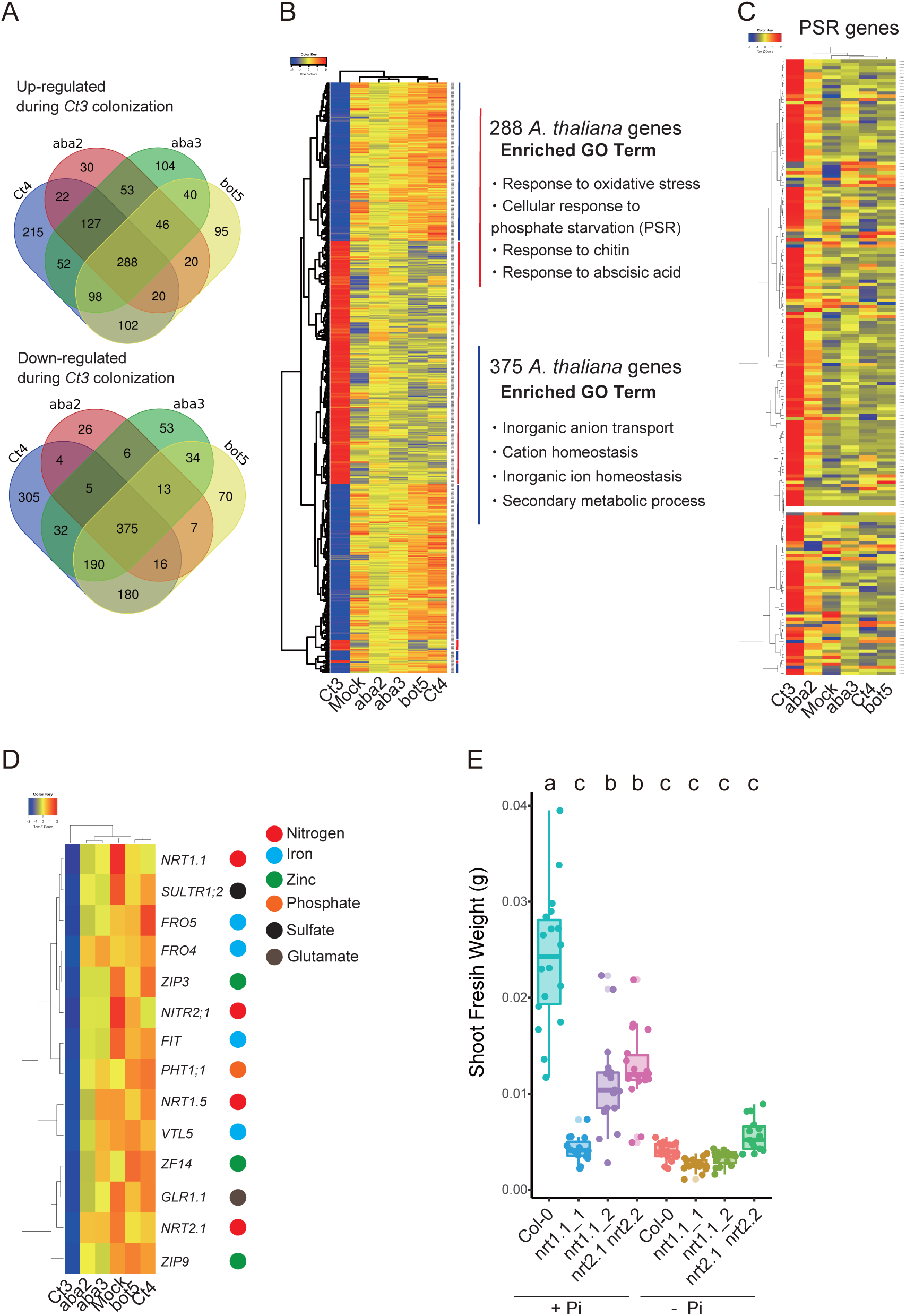
*Ct3* root colonization suppresses host nitrogen uptake genes while activating host PSR genes based on the ABA-BOT cluster. **a.** Venn diagram representing significantly up (Upper)- or down (Down)-regulated *A. thaliana* genes in *Ct3* treatment compared with each treatment (*Ct4*, *aba2, aba3*, and *bot5*) (|log2FC|>1, FDR < 0.05) at 10 dpi under low Pi. **b.** Transcript profiling of 375 commonly upregulated and 288 commonly down-regulated *A. thaliana* genes in *Ct3* treatment compared with *Ct4*-, *aba2*-, *aba3*-, and *bot5*-colonized roots. Overrepresented (red to yellow) and underrepresented (yellow to blue) modules are depicted as log10 (fpkm + 1). The major enriched GO of 288 or 375 genes IDs are described. Red and blue lines next to gene IDs represent upregulated or downregulated *A. thaliana* genes, respectively. **c.** Hierarchical clustering of *A. thaliana* PSR 193 genes. Overrepresented (red to yellow) and underrepresented (yellow to blue) modules are depicted as log10 (fpkm + 1). **d.** Hierarchical clustering of *A. thaliana* genes related to nutrient uptake whose expression was significantly suppressed by *Ct3* that depends on the ABA-BOT cluster (FDR < 0.05). Overrepresented (red to yellow) and underrepresented (yellow to blue) modules are depicted as log10 (fpkm + 1). **e.** Quantitative measurement of *A. thaliana* fresh shoot weight under normal (+) or low (-) Pi. The fresh weight of each plant was measured after 24 days of incubation. Different letters indicate significantly different statistical groups (ANOVA, Tukey-HSD test, p < 0.05).

To test the biological significance of this suppression, we examined whether the genes repressed by *Ct3* ABA and BOT genes and by host ABA pathways under low Pi, contribute to plant growth (Figures 5E and S5G-H). *Ct3-*repressible genes included *NRT1.1* and *NRT2.1*, two major transporters for nitrate uptake (Okamoto *et al*., 2003), and *FIT*, FER-like iron deficiency-induced transcription factor inducing Fe uptake genes (Colangelo and Guerinot, 2004). Disruption of *NRT1.1*, *NRT2.1*, or *FIT* resulted in strong growth deficits or chlorosis under normal Pi, in the absence of *Ct3*, whereas further decreases in plant growth by their disruption was not discernible under low Pi (Figures 5E and S5G-H). Although this has hampered assessing *Ct3* infection phenotypes in these mutant plants, the results suggest that these nutrition-related genes are rate-limiting in plant growth under Pi sufficiency, and that they define a host target for the fungal ABA-BOT cluster during *Ct3*-mediated plant growth inhibition.

## DISCUSSION

In this study we obtain evidence in a novel *Ct* strain (*Ct3*) that a dynamic transition from pathogenic to mutualistic lifestyles is achieved by altering the expression of a single fungal secondary metabolism cluster (ABA-BOT). It has been well documented that the presence or absence of one genetic component(s) in the genome, *e.g.* a virulent locus, island, plasmid or chromosome, is associated with the distinction between pathogenic and nonpathogenic (or potentially beneficial) lifestyles in different bacteria and fungi (Freeman and Rodriguez, 1993; Ma *et al*., 2010; Savory *et al*., 2017; Melnyk *et al*., 2019). In contrast, our studies reveal that the beneficial and pathogenic *Ct* strains share a nearly identical ABA-BOT cluster in their genomes, and that its expression status plays a critical role in the diversification of fungal lifestyles.

Our studies further indicate that altering *Ct3* ABA-BOT cluster expression enables its lifestyle transition on the same host. The divergence between adapted (pathogenic) and non-adapted (nonpathogenic) fungal pathogens was often attributed to transcriptional induction of virulence-related genes in the former (Baetsen-Young *et al*., 2019; Kusch *et al*., 2021; Zuo *et al*., 2021). Our findings extend this view that the loss of ABA-BOT cluster not only results in the suppression of *Ct3* pathogenesis but the expression of PGP function under low Pi, reminiscent of other beneficial *Ct* strains. This is also the case at high temperatures, where not only *Ct3* virulence suppression but PGP is also achieved, likely through the host *PHR1/PHL1*-dependent suppression of ABA-BOT cluster expression. The results highlight an important role played by a secondary metabolism gene cluster in the plastic lifestyle transition of plant-infecting fungi, and the convergence of environmental and host modulations on its transcriptional regulation.

How does *Ct3 ABA-BOT* cluster inhibit plant growth? Under our conditions, *Ct3* and *Ct4* are largely indistinguishable in root colonization levels (Figure 3), making it unlikely that *Ct3* pathogenesis is caused by fungal overgrowth in the roots. It has been described that bacterial and fungal pathogens mobilize the host ABA pathway to suppress plant immunity, in particular salicylic acid (SA)-based defenses (Zabala *et al*., 2009; Pieterse *et al*., 2009). However, our transcriptomic analyses on *Ct3* wild type and *aba/bot* fungi did not detect *ABA/BOT*-dependent alterations in plant induction of defense-related genes. Consistently, beneficial *Ct61* promotes plant growth in the plants simultaneously disrupted with defense-related hormones, SA, jasmonate, ethylene and defense regulator *PAD4* as well as in the WT plants, without fungal overgrowth (Hiruma *et al*., 2016). These data suggest that *Ct3*-mediated virulence is not expressed through ABA-SA antagonism. Rather, our transcriptome and genetic data suggest that *Ct3* utilizes ABA-BOT cluster to suppress the host genes required for the acquisition of different key nutrients, such as nitrogen, iron and zinc, which are rate-limiting in plant growth particularly when phosphate is sufficient. These results suggest that *Ct*3 virulence via ABA-BOT occurs at least in part through perturbation of host nutrition.

Host plants require *PHR1/PHL1* for suppression of fungal overgrowth in beneficial interactions with *Ct61*, a prerequisite for PGP (Hiruma *et al*., 2016). This work further shows a critical role of *PHR1/PHL1* in restricting *Ct3* pathogenesis. Consistently, PSR-related *PHR1/PHL1* regulons are induced during both pathogenic and beneficial interactions, albeit at different timings. In *Ct3*, ABA-BOT genes serve to accelerate early induction of host PSR-related genes (at 10 dpi) even under Pi sufficient conditions, likely through the host ABA core pathways. This may reflect a positive role for ABA in PSR under low Pi (Aleksza *et al*., 2017; Zhang *et al*., 2022). In contrast, beneficial *Ct* strains induce host PSR-related genes specifically under low Pi, at a later phase (24 dpi), coincident with the appearance of PGP effects. Given the absence of *ABA/BOT* gene expression in beneficial *Ct* strains, their PSR activation seems independent of fungal ABA or BOT biosynthesis. Indeed, like *Ct3*, beneficial *Ct* strains efficiently colonize *A. thaliana* roots under low Pi, without expressing *ABA/BOT* genes (Figure 3). Therefore, separate fungal mechanisms, in terms of ABA dependence, seem to confer host PSR activation between pathogenic *Ct3* and beneficial *Ct* interactions. Conversely, *PHR1/PHL1* restricts fungal growth under low Pi, through the suppression of separate fungal infection strategies between *Ct3* and beneficial *Ct* strains, diverged in ABA-BOT dependence. Our evidence attributes *PHR1/PHL1*-mediated alleviation of *Ct3* pathogenesis to the suppression of fungal ABA-BOT expression, under low Pi and at high temperatures. Plant transcriptome data not detecting differential defense gene regulation, imply that *PHR1/PHL1*-mediated control of fungal growth is also distinct from the previously described suppression of pattern-triggered immunity (Castrillo *et al*., 2017; Tang *et al*., 2022).

Interestingly, the loss of *PHR1/PHL1* allows *Ct3 aba3* and *bot5* fungi to promote plant growth in *phr1 phl1* at elevated temperatures, which is not seen in the WT plants (Figure 3G). This indicates that *PHR1/PHL1* are not required for PGP *per se*, which *Ct3* confer at high temperatures when ABA-BOT cluster is disrupted. In addition to stimulating host ABA signaling, ABA-BOT may suppress PGP function, *i.e.,* promoting host Pi acquisition, conserved in *Ct* species including *Ct3* (suggested by Figure 3G). This also seems to apply to pathogenic *Ci*, in which inherent ABA-BOT expression is associated with a massive decrease in phosphorus transfer to the host compared to beneficial *Ct61* (Hiruma *et al*., 2016). Notably however, in *Ct3,* this PGP function is likely suppressed by *PHR1/PHL1*, suggested by the absence of PGP in WT plants by *aba3/bot5* fungi (Figure 3G). It is conceivable that *PHR1/PHL1* regulons activated at 10 dpi with *Ct3* contributes to the suppression of PGP, given the previously described negative role for *PHR1* in nitrate transporter expression, including *NRT1.1*, required for plant growth (Medici *et al*., 2015; Kiba *et al*., 2018; Maeda *et al*., 2018; Wang *et al*., 2020). The mechanisms underlying multifaceted functions of *PHR1/PHL1* in the complex host-fungus interactions require further studies.

Consistent with our molecular phylogenetic analysis that the beneficial strains are derived from the pathogenic ancestors (e.g., *C.i* or *C. liriopes*), the present evidence indicates that the acquisition and expression of ABA-BOT gene cluster (or ABA and BOT gene clusters) contributes to fungal pathogenesis, likely through the mobilization of host ABA biosynthesis and signaling during infection. It seems that ABA and BOT gene clusters also contribute to fungal pathogenesis in *Botrytis* as shown for *Ct*, since host ABA signaling and fungal BOT genes are required for the infection and pathogenesis of the necrotrophic fungus (Audenaert *et al*, 2002; Liu *et al,* 2015; Siewers *et al*., 2005). Although the definite proof remains to be obtained, ABA-BOT gene clusters are likely to confer virulence in phylogenetically diverse fungi beyond *Ct* species. The additional requirements for host ABA biosynthesis in fungal pathogenesis imply that these fungal genes alone are not sufficient for the effective ABA biosynthesis and virulence. Future studies will be required for elucidating the precise mechanisms by which fungal and plant ABA pathways work in concert to promote fungal virulence.

Our findings indicate that a single fungal metabolism cluster enables the expression and evolution of virulence in otherwise endophytic (asymptomatic) PGP fungi. This predicts that manipulation of a very limited number of virulence factors in fungal pathogens is sufficient to prevent their virulence and even possible to gain host benefits from their colonization. A better understanding of how endophytic fungi regulates virulence expression is important for elucidating their plastic infection mechanisms and promoting their effective use in agricultural settings.

## Acknowledgments

We thank Mie Matsubara, Akemi Uchiyama, Sachiko Minezaki, Shunsuke Imai, Hiromi Haba, and Takafumi Shimizu for technical assistance. In addition, we thank Paul Schulze-Lefert, Soledad Sacristán, Ryo Tabata, Chika Tateda, and Yuri Tajima for fruitful discussion. We thank Yasuyuki Kubo, Maarten Koornneef, Javier Paz-Ares, Shigetaka Yasuda, and Haruhiko Inoue for published materials. This work was supported in part by the Japan Society for the Promotion of Sciences (JSPS) KAKENHI Grant (16H06279, 18K14466, 18H04822, 19H05688, 20H02986, 21H05150), the Japan Science and Technology Agency (JST) grant (JPMJPR16Q7, JPMJCR19S2, JPMJSC1702, JPMJFR200A). Computations were partially performed on the NIG supercomputer at ROIS National Institute of Genetics.

## Author Contributions

KH initiated the project. KH and YS directed the research. KH, MO, NK, MN, YO, RS, KT, and TS conducted the experiments. YU conducted comparative genomic analyses shown in Figure 1 and Table S2. SA conducted analyses of molecular phylogenetic trees in Figure 1, Figure 4 and the related supplementary data, supervised by WI. KH conducted RNA-seq analyses. TS provided unique material. KH, SA, MO, YU, NK, MN, YO, RS, KT, and WI analyzed the data. KH, SA, WI, and YS wrote the manuscript with feedback from all authors.

## Declaration of Interests

No

## STAR Methods

### Key resources table

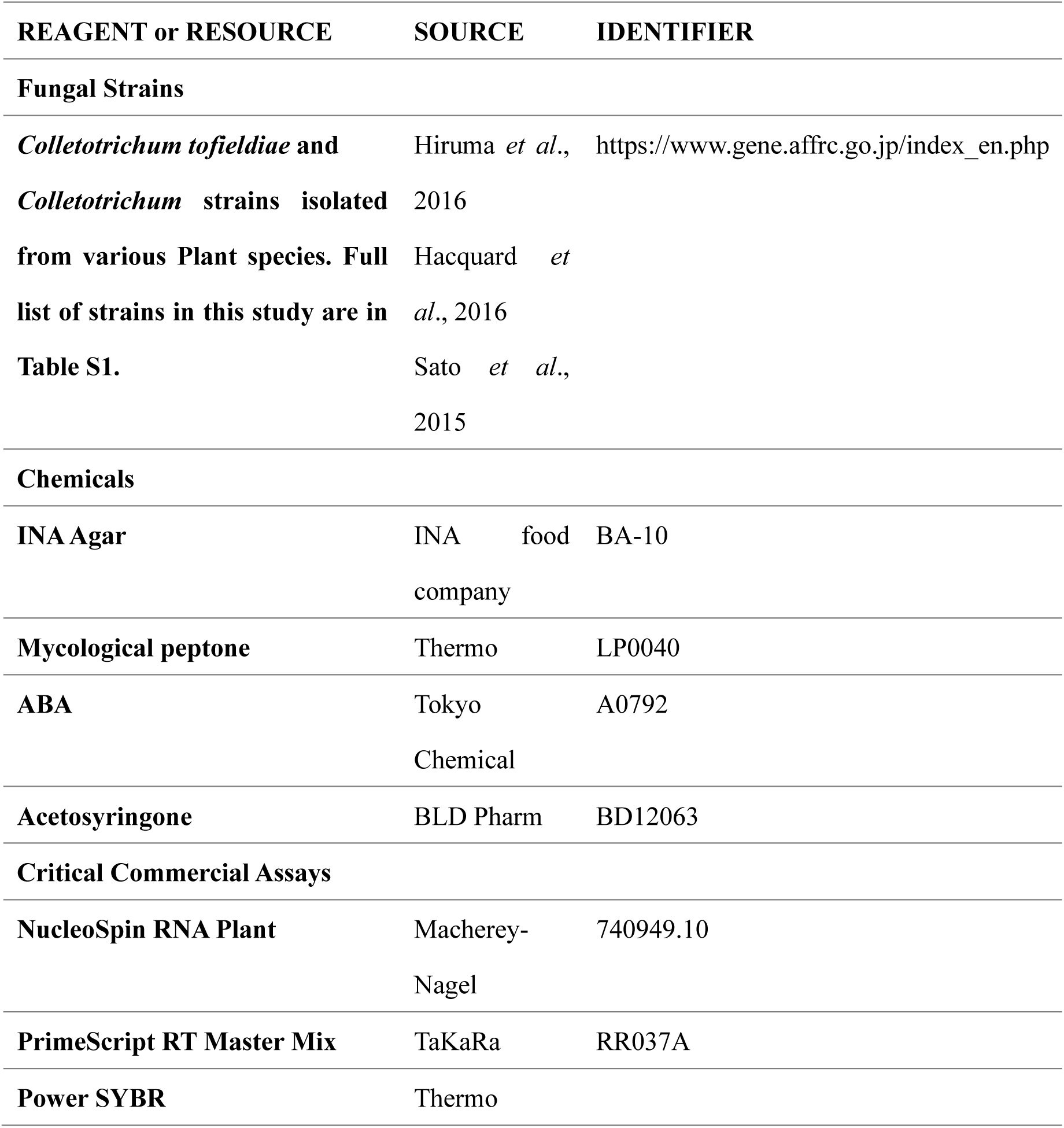
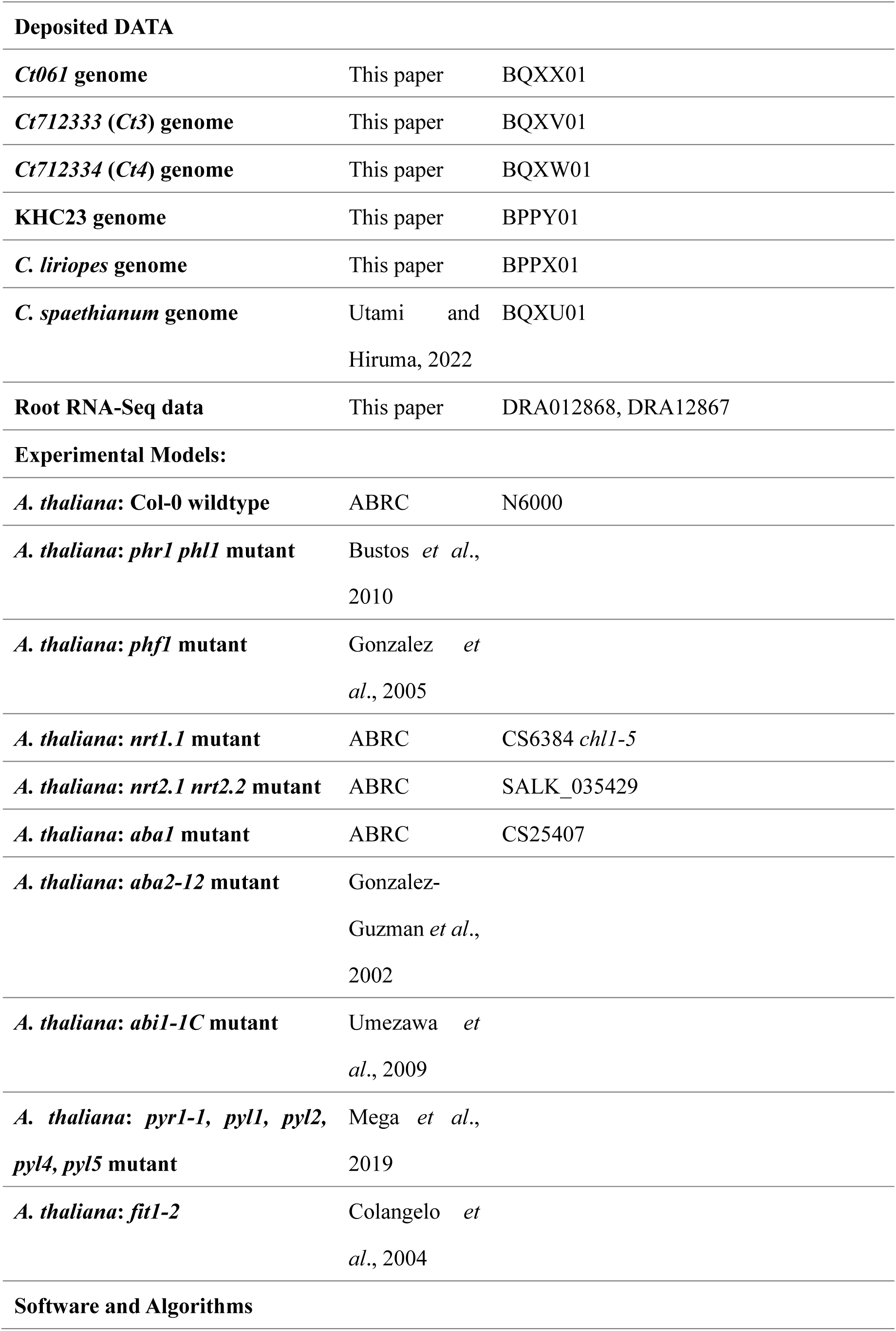
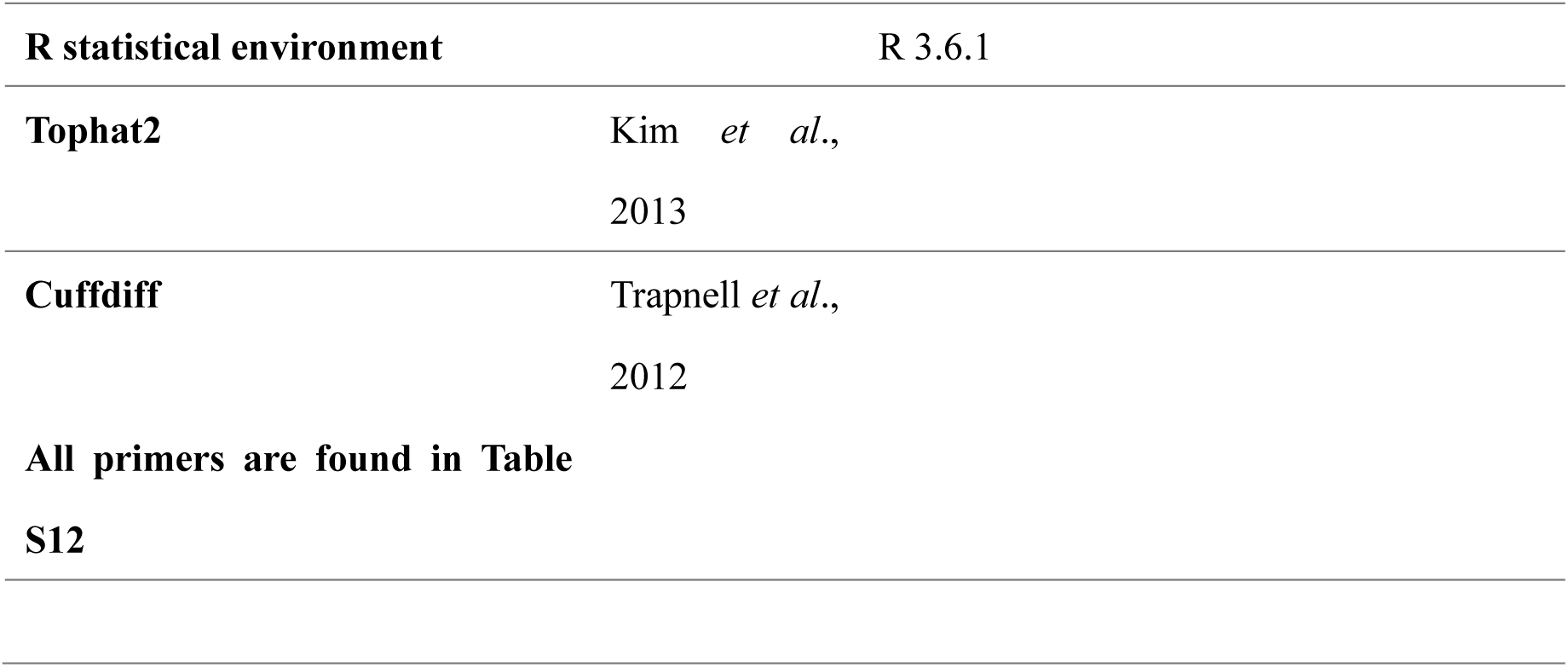

| REAGENT or RESOURCE | SOURCE | IDENTIFIER |
| --- | --- | --- |
| **Fungal Strains** |  |  |
| *Colletotrichum tofieldiae* and *Colletotrichum* strains isolated from various Plant species. Full list of strains in this study are in Table S1. | Hiruma *et al.*, 2016<br>Hacquard *et al.*, 2016<br>Sato *et al.*, 2015 | https://www.gene.affrc.go.jp/index_en.php |
| **Chemicals** |  |  |
| INA Agar | INA food company | BA-10 |
| Mycological peptone | Thermo | LP0040 |
| ABA | Tokyo Chemical | A0792 |
| Acetosyringone | BLD Pharm | BD12063 |
| **Critical Commercial Assays** |  |  |
| NucleoSpin RNA Plant | Macherey-Nagel | 740949.10 |
| PrimeScript RT Master Mix | TaKaRa | RR037A |
| Power SYBR | Thermo |  |
| **Deposited DATA** |  |  |
| *Ct061* genome | This paper | BQXX01 |
| *Ct712333 (Ct3)* genome | This paper | BQXV01 |
| *Ct712334 (Ct4)* genome | This paper | BQXW01 |
| **KHC23 genome** | This paper | BPPY01 |
| *C. liriopes* genome | This paper | BPPX01 |
| *C. spaethianum* genome | Utami and Hiruma, 2022 | BQXU01 |
| **Root RNA-Seq data** | This paper | DRA012868, DRA12867 |
| **Experimental Models:** |  |  |
| *A. thaliana*: Col-0 wildtype | ABRC | N6000 |
| *A. thaliana*: *phr1 phl1* mutant | Bustos *et al.*, 2010 |  |
| *A. thaliana*: *phf1* mutant | Gonzalez *et al.*, 2005 |  |
| *A. thaliana*: *nrt1.1* mutant | ABRC | CS6384 *chl1-5* |
| *A. thaliana*: *nrt2.1 nrt2.2* mutant | ABRC | SALK_035429 |
| *A. thaliana*: *aba1* mutant | ABRC | CS25407 |
| *A. thaliana*: *aba2-12* mutant | Gonzalez-Guzman *et al.*, 2002 |  |
| *A. thaliana*: *abi1-1C* mutant | Umezawa *et al.*, 2009 |  |
| *A. thaliana*: *pyr1-1, pyl1, pyl2, pyl4, pyl5* mutant | Mega *et al.*, 2019 |  |
| *A. thaliana*: *fit1-2* | Colangelo *et al.*, 2004 |  |
| **Software and Algorithms** |  |  |

|  |  |
| --- | --- |
| **R statistical environment** | R 3.6.1 |
| **Tophat2** | Kim *et al.*,<br>2013 |
| **Cuffdiff** | Trapnell *et al.*,<br>2012 |
| **All primers are found in Table S12** |  |

## RESOURCE AVAILABILITY

### Lead Contact

Further information and requests for resources and reagents should be addressed to the Lead Contact, Kei Hiruma.

### Material Availability

All *A. thaliana* materials and WT fungal strains used in this study are publicly available. In addition, the *Ct3* mutants are available from the lead contact.

### Data Availability

Newly obtained fungal whole-genome data have been deposited in DDBJ. Additionally, raw sequencing data of RNA-seq transcriptomic data have been deposited in DDBJ (for Figure 2, Figure 3, and Figure 5).

## EXPERIMENTAL MODEL AND SUBJECT DETAILS

### Fungal strains

All fungal strains used in this study are described in Table S1.

### Plant material and growth conditions

*Arabidopsis thaliana* Col-0, the *abi1-1C* (Umezawa *et al*., 2009), *aba1* (Morris *et al*., 2006), *aba2-12* (Gonzalez-Guzman *et al*., 2002), *pyr1-1 pyl1 pyl2 pyl4 pyl5* (Mega *et al*., 2019), *phf1* (González *et al*., 2005), *phr1 phl1* (Bustos *et al*., 2010), *nrt1.1 (chl1-5,* Tsay, *et al*., 1993), *nrt2.1nrt2.2* (Li *et al*., 2007), and *fit1-2* (Colangelo and Guerinot, 2004) mutants (Col-0 background) and *nrt1.1_2* (Ler background) were used in this study. We also used 16 *A. thaliana* WT accessions as described in Figure S1. Seeds were surface sterilized with 70% ethanol for the 30s, followed by 6% sodium hypochlorite with 0.01% Triton X for 15 min. After being washed three times in sterilized water, seeds were placed in cold treatment at 4°C for 48 h before sowing.

### Fungal inoculation assay

Sterilized *A. thaliana* seeds were sowed on half MS agarose medium containing defined Pi concentrations. *Ct* spores (3 μL of 1 × 10^4^ spores/ml) were inoculated 3 cm below the sowed seeds. Plates were placed vertically in a plant growth chamber under a 10:12 hr light: dark cycle at ∼ 22°C (± 1°C) (80 μE/m^2^s) around 50% humidity unless otherwise described. For 26°C assays, plants were incubated in a plant growth chamber under a 10:12 hr light: dark cycle at ∼26°C (80 μE/m^2^s) during light, and ∼22°C during dark. Effects on plant growth by fungal inoculation were evaluated by measuring fresh shoot weight, root length, or chlorophyll a and b from ∼15 plants per experiment. *Ct* hyphae were mixed with soils for soil inoculation, and *B. rapa* seeds were incubated for 24 days on the soils before measuring SFW.

## METHOD DETAILS

### Plant growth conditions

Half-strength MS medium used in this study are based on Gruber *et al*. (2013), with minor modification. The MS medium contains MgSO_4_. 7H_2_O (750 μM), KH_2_PO_4_ (625 for plus Pi, 50 for low Pi), NH_4_NO_3_ (10300), KNO_3_ (9400), CaCl_2_.2H_2_O (1500), CoCl_2_.6H_2_O (0.055), CuCl_2_.2H_2_O (0.053), H_3_BO_3_ (50), KI (2.5), MnCl_2_.4H_2_O (50), Na_2_MoO_4_.2H_2_O (0.52), ZnCl_2_ (15), Na-Fe-EDTA (75), MES pH 5.5 (1000), and KCL (0 for normal Pi, 575 for low Pi) with 1% Agar. Agar granulated (Difco) was initially used as an agar containing limited phosphate in this study. However, as the new batches of the agars caused unstable growth in mock-treated plants under low Pi, INA Agar BA-10 (INA food) with further limited phosphate was used as an alternative. pH was adjusted to 5.1. 55 ml of the mixture was then poured into square Petri plates (15 × 15 cm, Greiner). For *Ct* inoculation in soils, *Ct* spores (50 mL of 1× 10^4^ spores/ml) or distilled water (mock treatment) were incubated on mixtures of autoclaved 180 g sawdust, 60 g rice bran, 60 g bran, and 180 ml water for 2 weeks, where mixtures with or without *Ct* hyphae were mixed with soils containing Kanuma soil, Exo sand, and black soil (15:42.5:42.5) with 3%–5% weight. Before measuring *SFW, B. rapa* seeds (Misaki, Sakata seeds) were then sowed in the soil and incubated for 24 days.

### Leaf chlorophyll measurements

The method of chlorophyll quantification was adapted from Porra *et al*., 1989. Samples were prepared from ∼100 mg of leaf tissue pooled from ∼4 plants per sample and weighed. Chlorophyll was extracted by adding 800 µL chilled cold 100% acetone, and samples were shaken for 10 min until plant tissue was transparent. After the samples were diluted four times by 80% acetone, the absorbance of tissue-free chlorophyll extract was measured at 646 nm, 663 nm, and 750 nm with an Eppendorf Biospectrometer following formula.

A: Chlorophyll a (ug/ml) = 12.25 × (OD_663.6_ − OD_750_) − 2.85 × (OD_646.6_ − OD_750_)

B: Chlorophyll b (ug/ml) = 20.31 × (OD_646.6_ − OD_750_) − 4.91 × (OD_663.6_ − OD_750_)

### Quantitative real-time PCR

cDNA was synthesized from 300–500 ng total RNA using the PrimeScript RT Master Mix (Takara) in a volume of 10 µL. We then amplified 3 µL of cDNA (10 ng/µL) in Power SYBR (Thermo) with 1.6 µM primers using the AriaMx real-time PCR system (Agilent) in a volume of 12 µL. Primers used in this study are listed in Table S12.

### Fungal transformation via *Agrobacterium*-mediated transformation

For targeted gene replacement, we used the method described by Hiruma *et al*., 2016. Briefly, to generate replacement mutants lacking *Ct3* ABA biosynthesis genes (*ABA2* and *ABA3*) or BOT biosynthesis genes (*BOT1*, *BOT3*, and *BOT5*), we constructed a plasmid in which the Hyg resistance gene cassette was inserted between DNA sequences flanking the target gene. Around 1.5-kb fragments of 5′ (5F) and 3′ sequences (3F) flanking the gene ORF were amplified from *Ct3* genomic DNA by PrimeSTAR HS DNA polymerase (TaKaRa). The purified PCR products 5F and 3F were mixed with *Sal*I-treated pBIG4MRHrev for In-Fusion reactions. This generated plasmid was then introduced into *Agrobacterium* C58C1 strains by electroporation using the default setting for the Eppendorf Eporator. The transformed *Agrobacterium* strains (OD_600_ = 0.4) were mixed with *Ct3* spores (1 × 10^7^ spores) in a 1:1 ratio on a paper filter attached to incubation media containing 200 μM acetosyringone (BLD Pharm). Selection of hygromycin (150 µM)- resistant strains were conducted by transferring the paper filter to PDA media with Hyg, cefotaxime, and spectinomycin (all at 50 μg/mL) and incubated for two days before the paper was eliminated from the media. The resistant strains from media were tested by PCR using primers targeting the outside sequence and a Hyg sequence to determine whether the Hyg resistance cassette successfully replaced the interesting genes. Several independent knockout strains were used for further analyses. The PCR primers used are listed in Table S12.

### Fluorescence microscopy

Inoculated *A. thaliana* roots were observed through confocal microscopy. The observations were performed using a confocal laser scanning microscope Olympus FV1000 with filter settings suitable for fluorescein (FITC), i.e., blue excitation light. WGA-FITC conjugate (Sigma) was used to specify fungal cell walls as described in Both *et al*. (2005). Briefly, infected roots were incubated overnight in a 1:3 mixture of chloroform and ethanol. Then, the roots were transferred to chloral hydrate (2 g/mL in water) and incubated for 2 hours. After PBS washed roots, the roots were incubated in wheat germ agglutinin (WGA) conjugated to a FITC solution (5 ug/ml; Sigma) for 1 h at room temperature.

### Genome sequencing and assembly

CTAB with RNase treatment extracted fungal DNA from fungal hyphal aggregates grown on liquid Mathur’s medium (glucose 2.8 g/l, MgSO_4_×_7_H_2_O 1.2 g/l, KH_2_PO_4_ 2.7 g/l, and mycological peptone LP0040 2.2 g/l) for 3 days. Macrogen conducted the following procedures for genome sequencing and assembly. The genomes of *Ct3*, *Ct4*, and *Ct61* were obtained using PacBio RSII single-molecule real-time sequencing (4∼5 cells) (SMRT cell 8 Pac V3, DNA Polymerase Binding Kit P6). *De novo* assemblies of *Ct61*, *Ct3,* KHC, and *C. liriopes* genomes were provided via FALCON (v. 0.2.1). Proteins and CDSs data of *C. tofieldiae* 0861 strain (Hacquard *et al*., 2016) were downloaded from NCBI. The data were used for SNAP [1] (v2.31.8) gene training. Gene prediction was conducted using MAKER (v. 2.31.8) (Cantarel *et al*., 2008) based on the SNAP gene training model, and BLAST+ conducted gene annotation (v. 2.6.0; E-value: 1e^−3^, Databases NT, NR) with UniProt Swiss-Prot (2015/0214). *De novo* assembly of *Ct4* genomes was conducted by FALCON-integrate (v. 1.8.18). Illumina HiSeq for 100 bp paired-end short reads against *Ct4* genomes was also conducted by Magrogen (TrueSeq DNA PCR-Free kit), resulting in 59 million reads filtered by the criteria that a pair of PE reads for each read should have a base quality greater than or equal to Q20 for more than 90% of bases. The filtered sequence reads were used for error collection by mapping Hiseq reads to the PacBio assembly using Pilon (v. 1.21). Gene prediction was conducted using MAKER (v. 2.31.8) based on the SNAP gene training model obtained via CDSs and proteins of *C. tofieldiae* 0861 strain (Hacquard *et al*., 2016), and gene annotation was conducted using BLAST+ (v. 2.6.0; E-value: 1e^−3^, Databases NT, NR) with UniProt Swiss-Prot (2018/06).

Completeness of obtained draft genomes was estimated by BUSCO v.5.1.2 (Simão *et al*., 2015) based on the glomerellales_odb10 lineage dataset. Detection and annotation of tRNA and rRNA were conducted by tRNA-scan v.2.0.7 (Lowe and Chan, 2016) and barrnap v.0.9 (Seeman, 2018: https://github.com/tsee). Prediction of secretome was conducted as described in Crestana *et al*. (2021) using SignalP v.5.0 (Armenteros *et al*., 2019), TMHMM v.2.0 (Krogh *et al*., 2001), and ProSite PS-Scan against motif PS00014 (de Castro *et al*., 2006). Candidate effector proteins were identified using EffectorP v.3.0 (Sperschneider and Dodds, 2022). Carbohydrate-active enzymes (CAZymes) were predicted with the hmmscan function from HMMER v.3.1b2 (Eddy, 2011) using dbCAN HMMdb release 9.0 (Zhang *et al*., 2018) as a database. We detected candidates of secondary metabolites by anti-SMASH v.6.0.1 (Blin *et al*., 2021).

### Syntenies of *Ct3*, *Ct4*, and *Ct61* genomes

Syntenies of *Ct3*, *Ct4*, and *Ct61* genomes were detected as locally collinear blocks (LCB) partly by performing whole-genome alignment in Mauve program v1.1.1 (Darling *et al*., 2004). Location of LCBs and synteny correlations were visualized as circos diagram by package circlize v.0.4.3 (Gu, 2014) in R.

### Construction of phylogenetic trees of ABA and BOT biosynthesis genes

For this analysis, we re-annotated ABA and BOT biosynthesis *Ct* (*Ct3*, *Ct4*, and *Ct61*) genes based on both genome assembly and RNA-seq expression data for each gene. Sequences similar to ABA and BOT genes of *Colletotrichum* strains, *B. cinerea*, and some other strains were collected with BLASTP search using Biopython (v.1.7.6; E-value: 1e-4 or 1e-5) against newly analyzed genomes in this study, Homo sapiens genome assembly GRCh38.p13, and GenBank NR database on February/March 2020. Those containing large indels or that were highly diverged were omitted from the data set of sequences (Table S7). Sequences were aligned with MAFFT using the E-INS-i strategy (v.7.273) (Katoh and Standley, 2013). Phylogenetic analysis was conducted with IQ-TREE (v.1.6.11; -m MFP -nt AUTO -b 100) (Nguyen *et al*. 2015). Microsynteny of ABA and BOT genes was analyzed with GenomeMatcher (v.3.00) (Ohtsubo *et al*. 2008).

### RNA extraction and RNA-sequencing analyses

Total RNA was extracted using the NucleoSpin RNA Plant kit (Macherey-Nagel). RNA samples (1µg each) were then sent to BGI for quality assurance, library preparation, and subsequent sequencing using their default process. The generated libraries were sequenced by HiSeq-illumina, resulting in approximately 40 million reads per sample, paired-end 150 bp (Figures 2 and 3) or BGIseq, resulting in about 40 million reads per sample, paired-end 100 bp (Figure 5). Tophat2 with default settings, except that average fragment size was specified (Kim *et al*., 2013), was used for sequence mapping based on the *A. thaliana* genome (TAIR10) or fungal genomes (*Ct3*, *Ct4,* or *Ct61*). The generated BAM files from the Tophat2 platform were used to analyze differentially expressed genes by cuffdiff with default settings (Trapnell *et al*., 2012). The following statistical analysis and data visualization were conducted in R Bioconductor packages, such as Cummerbund, gplots, and ggplot2 (Ginestet 2011; Trapnell *et al*., 2012). As judged by the Cummerbund platform, DEGs were identified using an FDR threshold of < 0.05. Heatmaps representing gene expression profiles were generated with the R package heatmap.2 in gplots. GO analysis for *A. thaliana* genes was conducted by AgriGO v. 2 (FDR < 0.05, Tian *et al*., 2017). Motif analysis was conducted by CentriMo (Bailey and Machanick, 2012) with default setting using 1,000 bp upstream of the target *A. thaliana* genes extracted via TAIR Bulk Data Retrieval.

### ABA measurement

Extraction, purification, and quantification of ABA were carried out as described in Saika *et al*. (2007) using around 5-mg dry weights of around 10-day-seedling roots per replicate (equivalent to around 260 roots).

### Mineral measurements

Plant samples were dried for 3 days in 70 °C oven. Dried samples were set as 10 to 15 mg in one biological sample. Each sample was digested with nitric acid and hydrogen peroxide as described in the previous report (Ohmori *et al*., 2016). The dried pellets after digestion were dissolved in 0.08 M HNO_3_. Elemental concentrations in samples were measured by inductively coupled plasma mass spectrometry (ICP-MS) Agilent 7800 (Agilent Technologies Co., Ltd, Japan) according to manufacturer’s instruction. To determine the concentration of P, ICP-MS NexION 350S (PerkinElmer, Waltham, MA, USA) was used.

### Clustering analysis

Clustering analysis was done by using mineral concentrations of the samples. NbClust package (version 3.0) of R was used for k-means clustering (Hartigan-Wong algorithm). Optimal number of cluster was determined by silhouette score.

### Fungal growth assay *in vitro*

Agar plugs of ten-day-old fungal cultures grown on Mathur’s media were transferred to new Mathur’s plates or 1/2 MS plates containing low Pi (50 µM KH_2_PO_4_). Colony formation was determined three days after inoculation by measuring colony radius from center to edge. Fungal spore counts were conducted as described previously (Hiruma and Saijo, 2016). For fungal growth on glass slides, fungal spores were placed on glass slides and incubated for three days.

### Quantification and statistical analysis

Details of statistical analysis used in this study can be found in each figure legend. Boxplots display individual data points and median values. Statistical analyses were performed using Student’s two-tailed t-test or Tukey’s HSD, with correction for multiple comparisons.

## Supplementary figure Legends

**Supplementary Figure 1.**
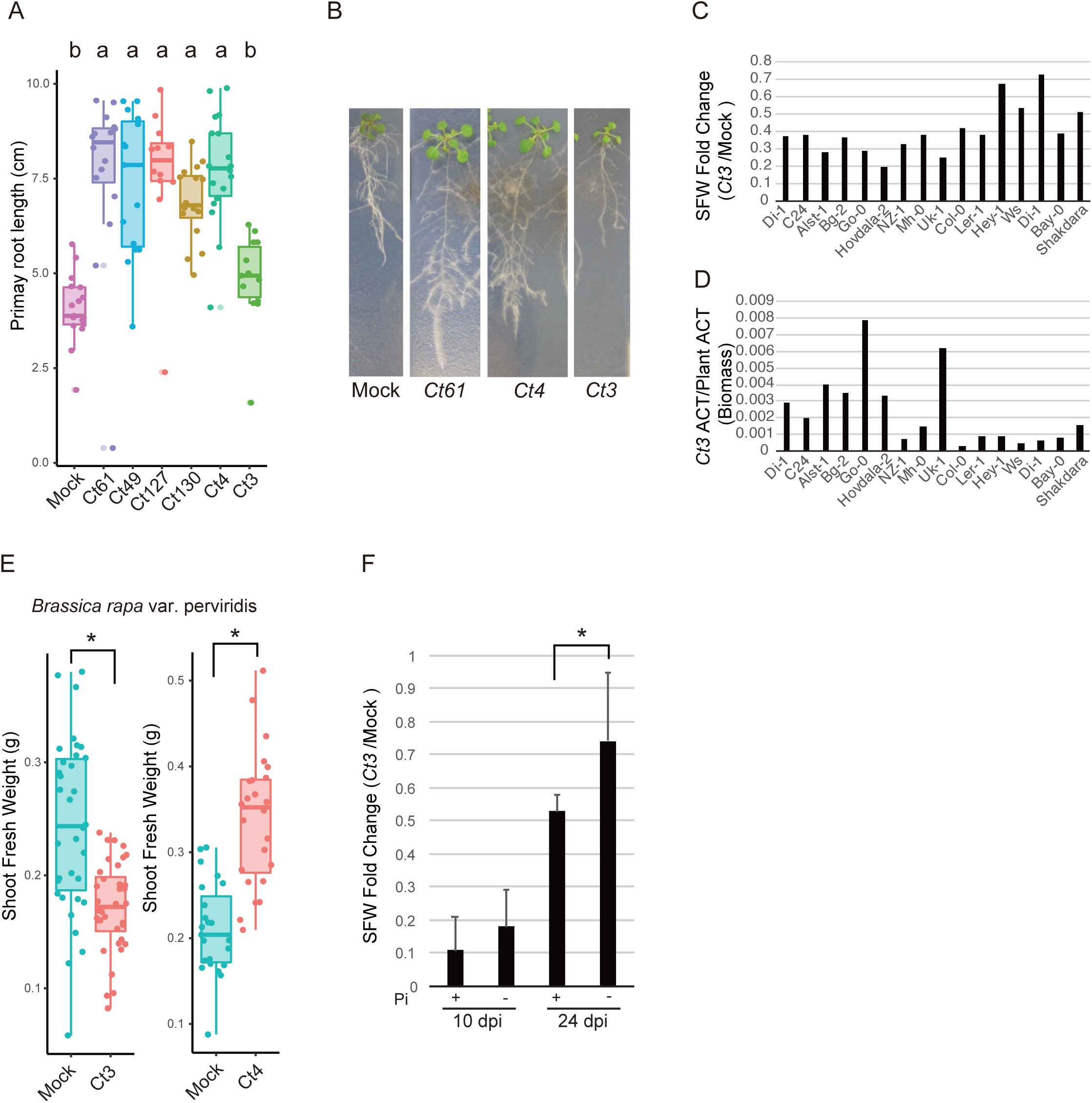
A *Ct* strain (*Ct3*) inhibits plant growth of *A. thaliana* and *B. rapa*. Related to Figure 1. **a.** Quantitative measurements of *A. thaliana* primary root length upon fungal inoculation. Plants were incubated for 24 days under low Pi (50 μM KH_2_PO_4_) from germination with mock treatment, *Ct61*, C*t49*, *Ct127*, *Ct130*, *Ct4* or *Ct3,* and root length was measured at 24 dpi. Each dot represents an individual plant sample. Different letters indicate significantly different statistical groups (ANOVA, Tukey-HSD test, p < 0.05). **b.** The representative *A. thaliana* plants inoculated by *Ct61*, *Ct4*, or *Ct3* under low Pi at 24 dpi. **c. d.** Inoculation of *Ct3* against 16 different *A. thaliana* accessions under low Pi. The negative impacts on plant growth by *Ct3* were estimated by SFW fold change (*Ct3*/Mock) at 24 dpi. At least four plants per accession and treatment were used to calculate SFW fold change. qRT-PCR analyses were used to measure fungal biomass as estimated by relative expression between *Ct* ACTIN and Plant ACTIN (*Ct* ACTIN/Plant ACTIN). **e.** Quantitative measurement of *B. rapa* SFW either upon *Ct4* or *Ct3* inoculation under low nutrient soils. Each dot represents an individual plant sample. Asterisks indicate significantly different statistical groups (p < 0.01, two-tailed t-test). **f.** Degree of *Ct3*-mediated plant inhibition under normal (+, 625 μM KH_2_PO_4_) and low (-) Pi conditions. Negative impacts by *Ct3* were estimated by SFW fold change (*Ct3*/mock). Asterisks indicate significantly different statistical groups (SD, p < 0.01, Two-tailed t-test). Bars represent SD.

**Supplementary Figure 2.**
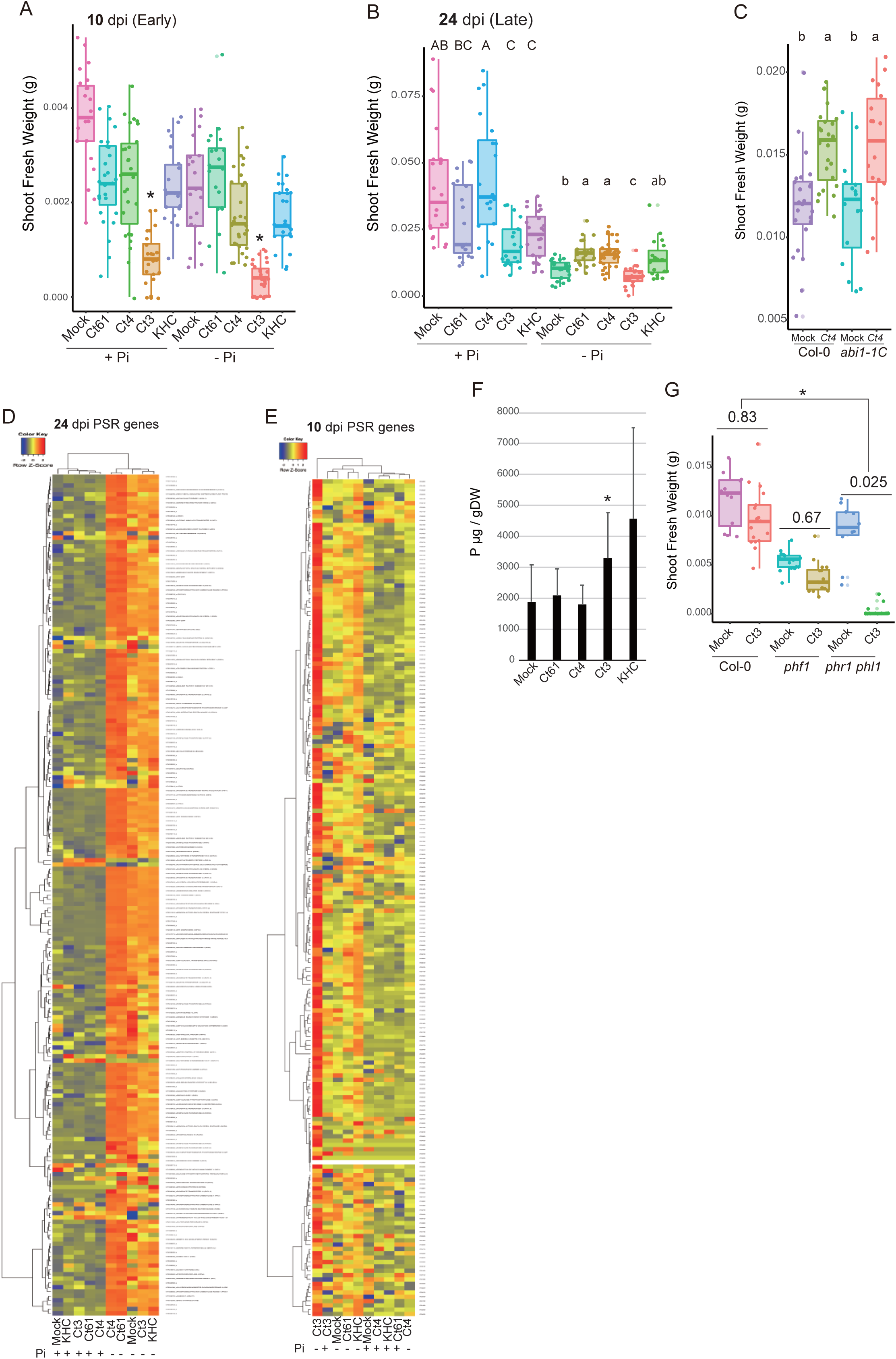
*Ct3* specifically induced *A. thaliana* ABA-responsive and phosphate starvation responsive genes during early root colonization. Related to Figure 2. **a.** Quantitative measurement of *A. thaliana* fresh shoot weighted upon fungal inoculation. Fresh weight of plants with mock-treated, *Ct61*, *Ct4*, *Ct3*, or KHC at 10 dpi was measured. The plant roots were used for the subsequent RNA-seq analysis. Asterisks indicate significantly different means between mock and *Ct3* treated plants under normal or low Pi (p < 0.05, two-tailed t-test). **b.** Quantitative measurement of *A. thaliana* SFW upon fungal root colonization at 24 dpi. Each dot represents an individual plant sample. The plant roots were used for the subsequent RNA-seq analysis. **c.** Quantitative measurement of *A. thaliana* SFW upon beneficial *Ct4* root colonization at 24 dpi. *Ct4* was inoculated onto either Col-0 or *abi1-1C* mutant plants and inoculated plants were incubated for 24 dpi under low Pi. Each dot represents an individual plant sample. Different letters indicate significantly different statistical groups (ANOVA, Tukey-HSD test, p < 0.05). **d. e.** Expression profiles for 193 *A. thaliana* PSR genes during root colonization by each fungus using RNA-seq data at (**d**) 24 dpi: 88 of 193 genes (*Ct61* vs. mock, FDR <0.05), 99 of 193 genes (*Ct4* vs. mock, FDR < 0.05), 30 of 193 genes (*Ct3* vs. mock, FDR < 0.05), and **(e)** 10 dpi. Overrepresented (light red to orange) and underrepresented (light blue to dark blue) transcripts are shown as log10 (fpkm + 1). **f.** Measurement of P concentration in *A. thaliana* shoots whose roots were inoculated and incubated with individual fungal strains for 10 day under low Pi. Bars represent SD (n = 9∼12). Asterisks indicate significant differences between mock and *Ct3* (p < 0.05, two-tailed t-test). **g.** Quantitative measurement of *A. thaliana* SFW upon *Ct3* root colonization at 28 dpi. *Ct3* was inoculated onto Col-0, *phf1*, and *phr1 phl1* plants and inoculated plants were incubated for 28 dpi under low Pi. Each dot represents an individual plant sample. Asterisks indicate significantly different SFW ratios (*Ct3*/mock) between Col-0 and *phr1 phl1* (p < 0.05, two-tailed t-test).

**Supplementary Figure 3.**
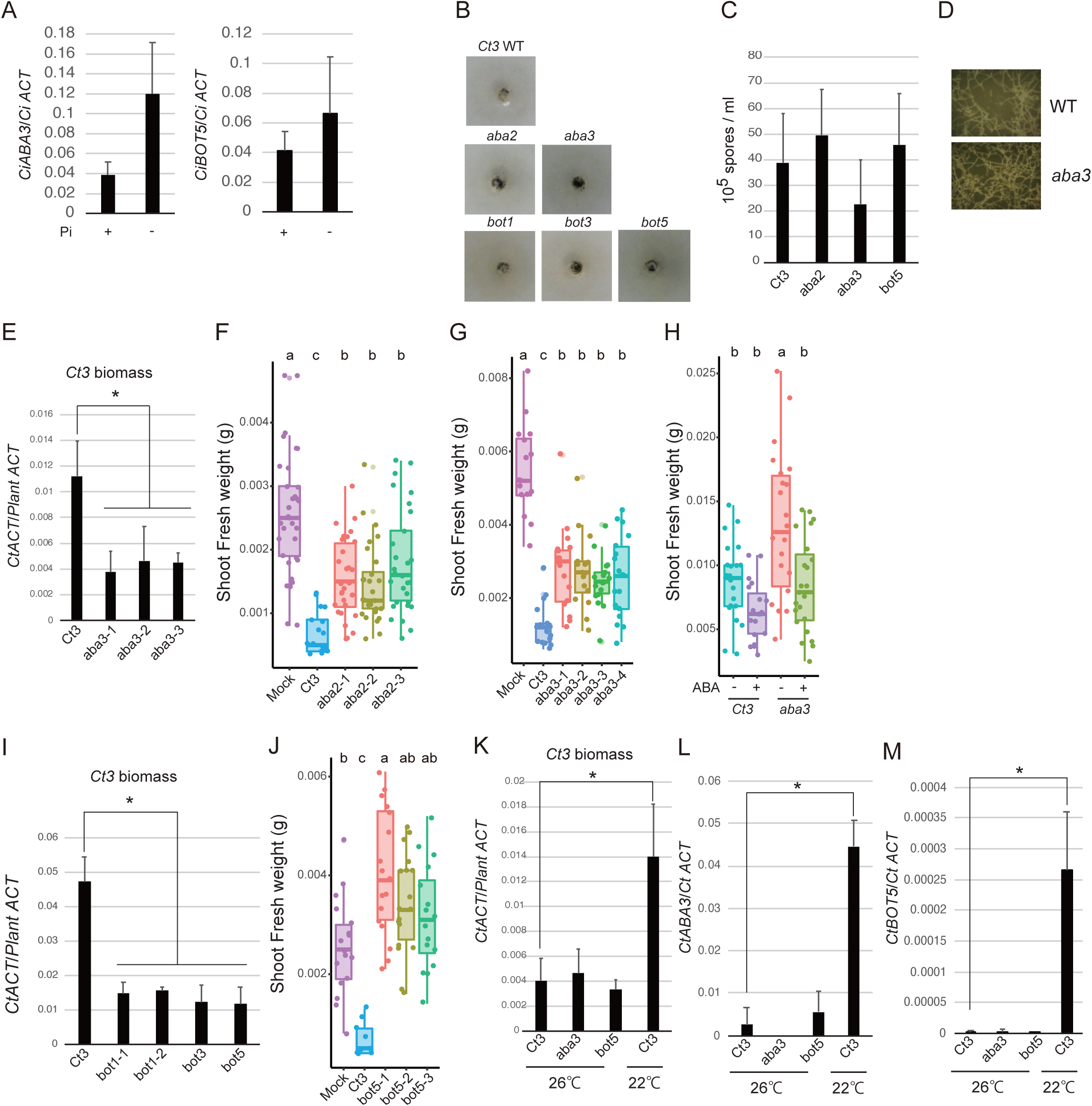
Fungal ABA and BOT biosynthesis genes are required for root colonization by *Ct3* and its virulence. Related to Figure 3. **a.** The expression of *CiABA3* or *CiBOT5* during root colonization by *C. incanum*. Plants were incubated with *C. incanum* for 23 dpi under normal (+) or low (-) Pi (SD, n =3). **b.** Fungal growth on low Pi media. Hyphae of *Ct3*WT, *aba2*, *aba3*, *bot1*, *bot3*, and *bot5* were transferred to low Pi media (without sucrose) and incubated for three days. **c.** Measurements of fungal spores generated from fungal strains grown in Mathur’s nutrient media. Spores generated from *Ct3*WT, *aba2*, *aba3*, and *bot5* were measured using a hemocytometer (SD, n = 3). **d.** Spore germination and hyphal growth on glass slides. The same spore amount of *Ct3*WT or *aba3* was placed on glass slides and incubated for two days after which pictures were captured on the same scale. **e.** Fungal biomass measurements in *A. thaliana* roots. *Ct3*WT, *aba3-1*, *aba3-2,* and *aba3-3* were inoculated onto *A. thaliana* roots, and inoculated plants were incubated for 24 dpi under low Pi, after which roots were collected. qRT-PCR analyses were used to measure fungal biomass as estimated by relative expression between *Ct* ACTIN and Plant ACTIN (*Ct* ACTIN/Plant ACTIN) (SD, n =3, p < 0.05, two-tailed t-test). **f, g.** Quantitative measurement of *A. thaliana* SFW upon *Ct3* root colonization at 24 dpi under low Pi. Each dot represents an individual plant sample. Different letters indicate significantly different statistical groups (ANOVA, Tukey-HSD test, p < 0.05). **h.** Quantitative measurement of *A. thaliana* SFW upon fungal root colonization at 24 dpi with and without ABA treatment. *Ct3* or *Ct3 aba3* was incubated with *A. thaliana* for 24 dpi on low Pi media with 10 μM ABA. Different letters indicate significantly different statistical groups (ANOVA, Tukey-HSD test, p < 0.05). **i.** Fungal biomass measurements in roots. *Ct3*WT, *bot1-1*, *bot1-2*, and *bot1-3* were inoculated onto *A. thaliana* roots, and inoculated plants were incubated for 24 dpi under low Pi. qRT-PCR was conducted to measure fungal biomass in *A. thaliana* roots, as estimated by *Ct* ACTIN/Plant ACTIN (SD, n =3, p < 0.05, two-tailed t-test). **j.** Quantitative measurement of *A. thaliana* SFW upon fungal root colonization. The SFW of *A. thaliana* with roots incubated either with *Ct3*WT, *bot5-1*, *bot5-2*, or *bot5-3* for 24 dpi under low Pi were measured. Different letters indicate significantly different statistical groups (ANOVA, Tukey-HSD test, p < 0.05). **k.** Fungal biomass measurements in roots under 22°C or 26°C. Inoculated plants were incubated under 22°C or 26°C at 24 days, and the fungal biomass was measured using qRT-PCR (SD, n =3∼4, *Ct ACTIN*/Plant *ACTIN*). **l, m.** The expression of *Ct3ABA3* or *Ct3BOT5* in roots under 22°C or 26°C. Inoculated plants were incubated under 22°C or 26°C at 24 days, and the fungal biomass was measured using qRT-PCR (*ABA3*/*Ct ACTIN* or *BOT5*/*Ct ACTIN*) (SD, n =3∼4, p < 0.05, two-tailed t-test).

**Supplementary Figure 4.**
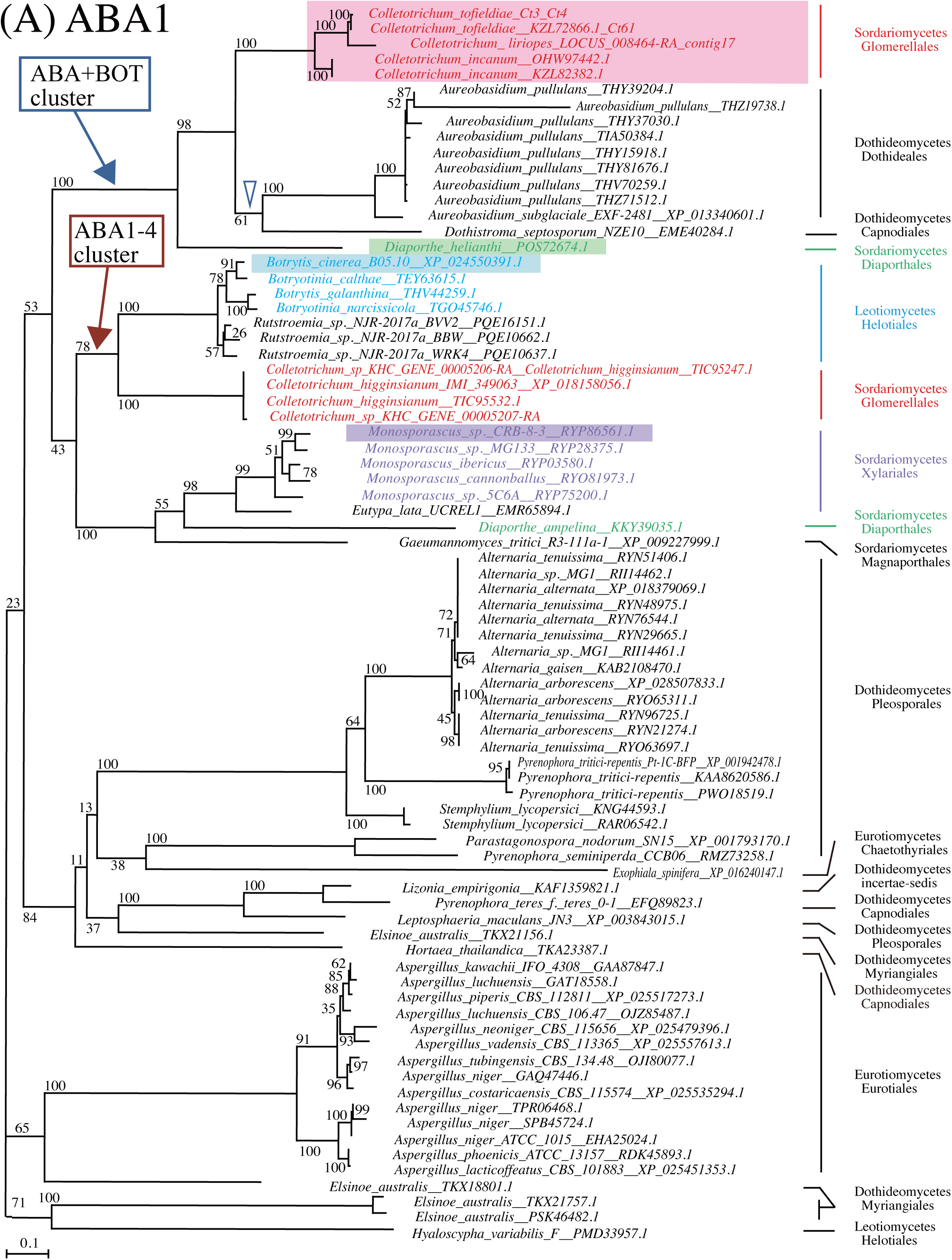
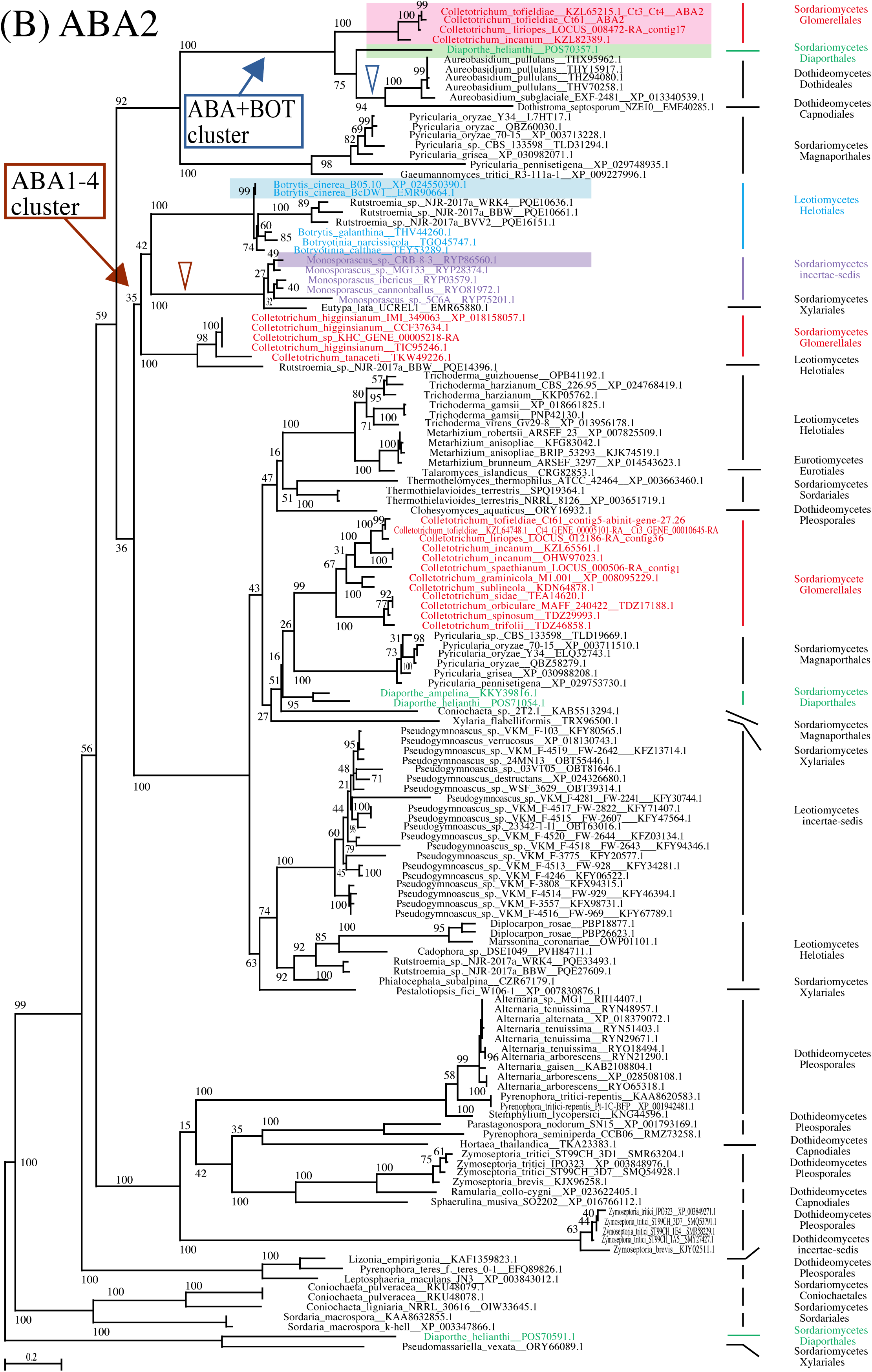
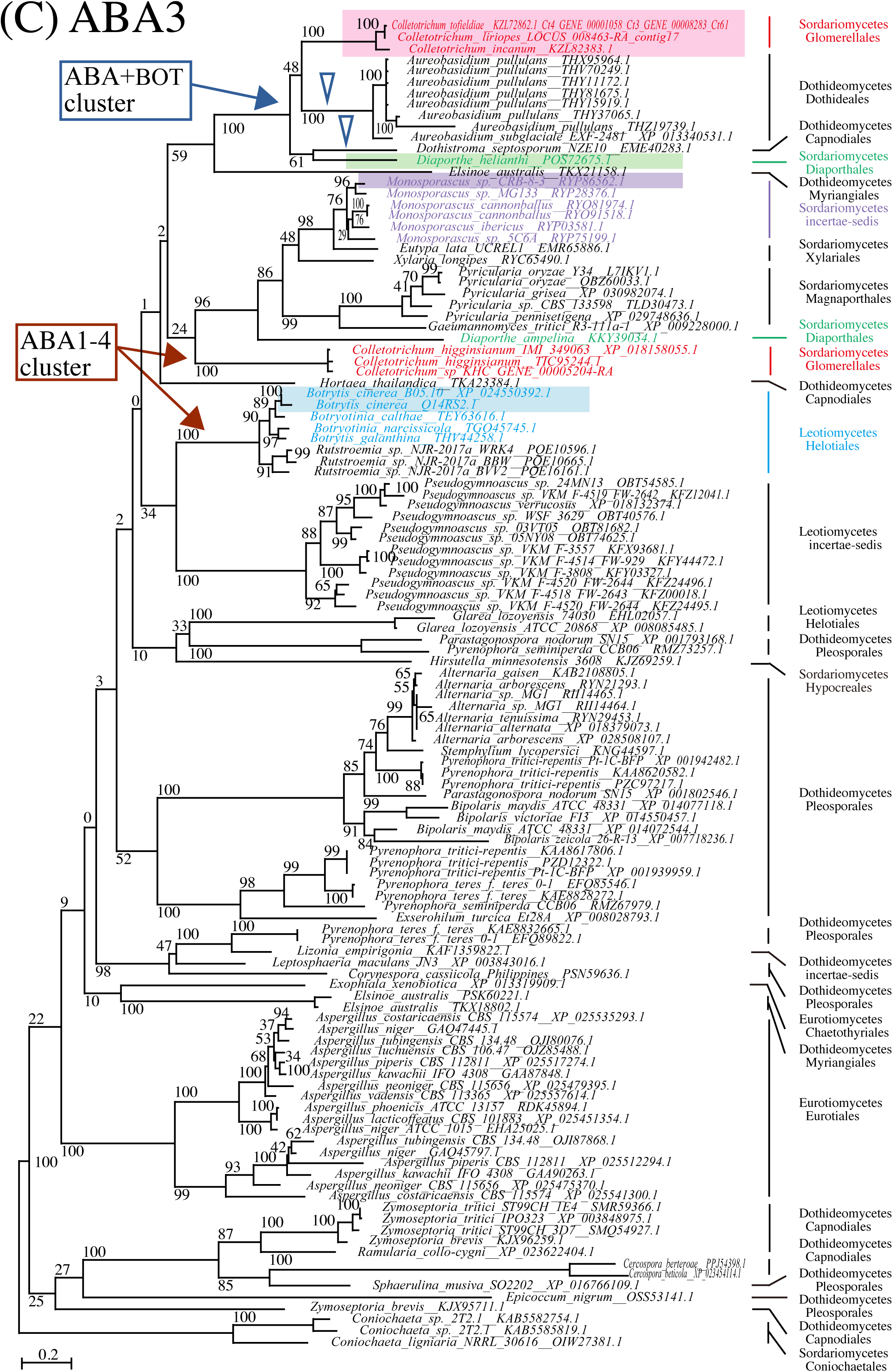
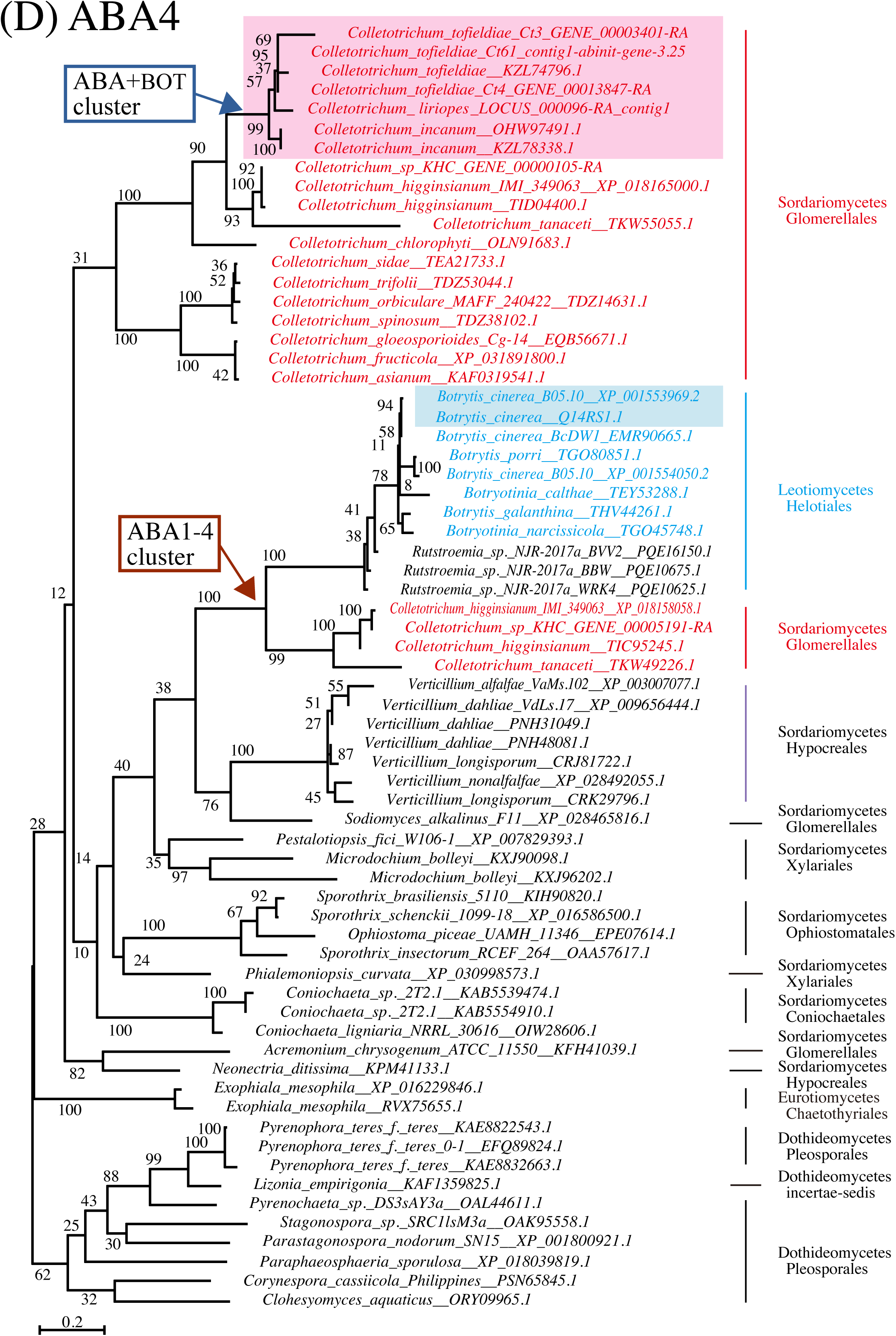
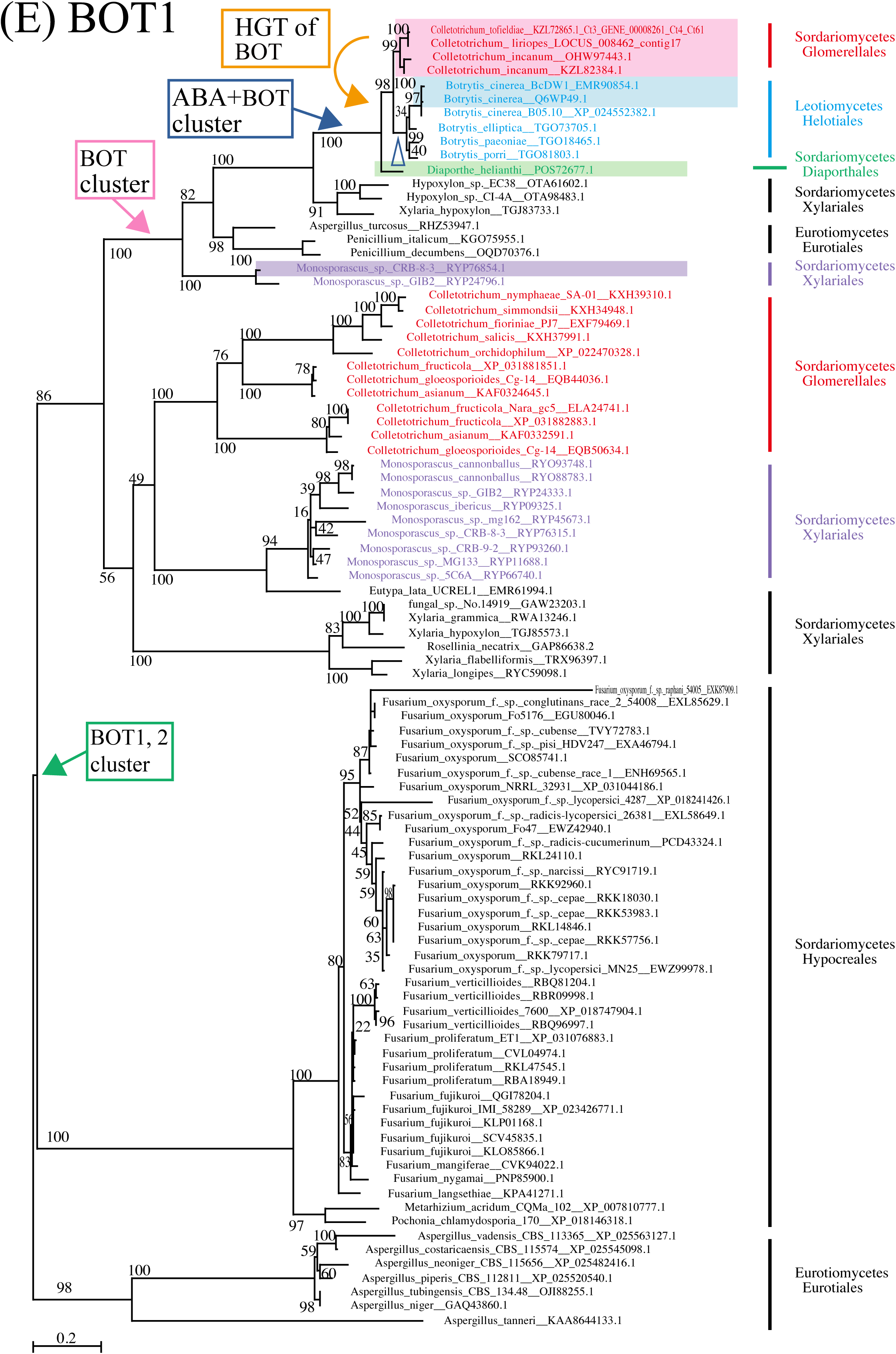
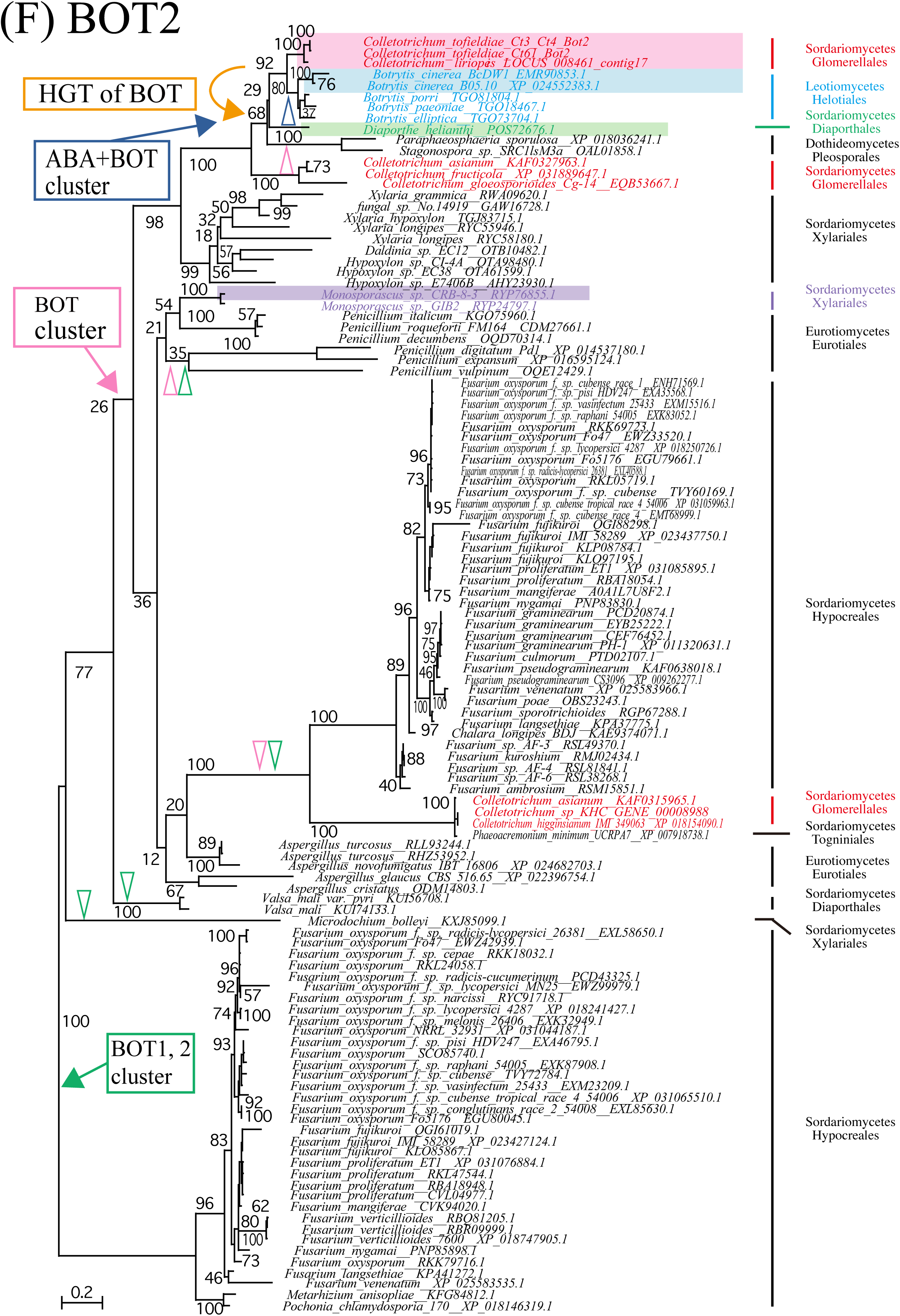
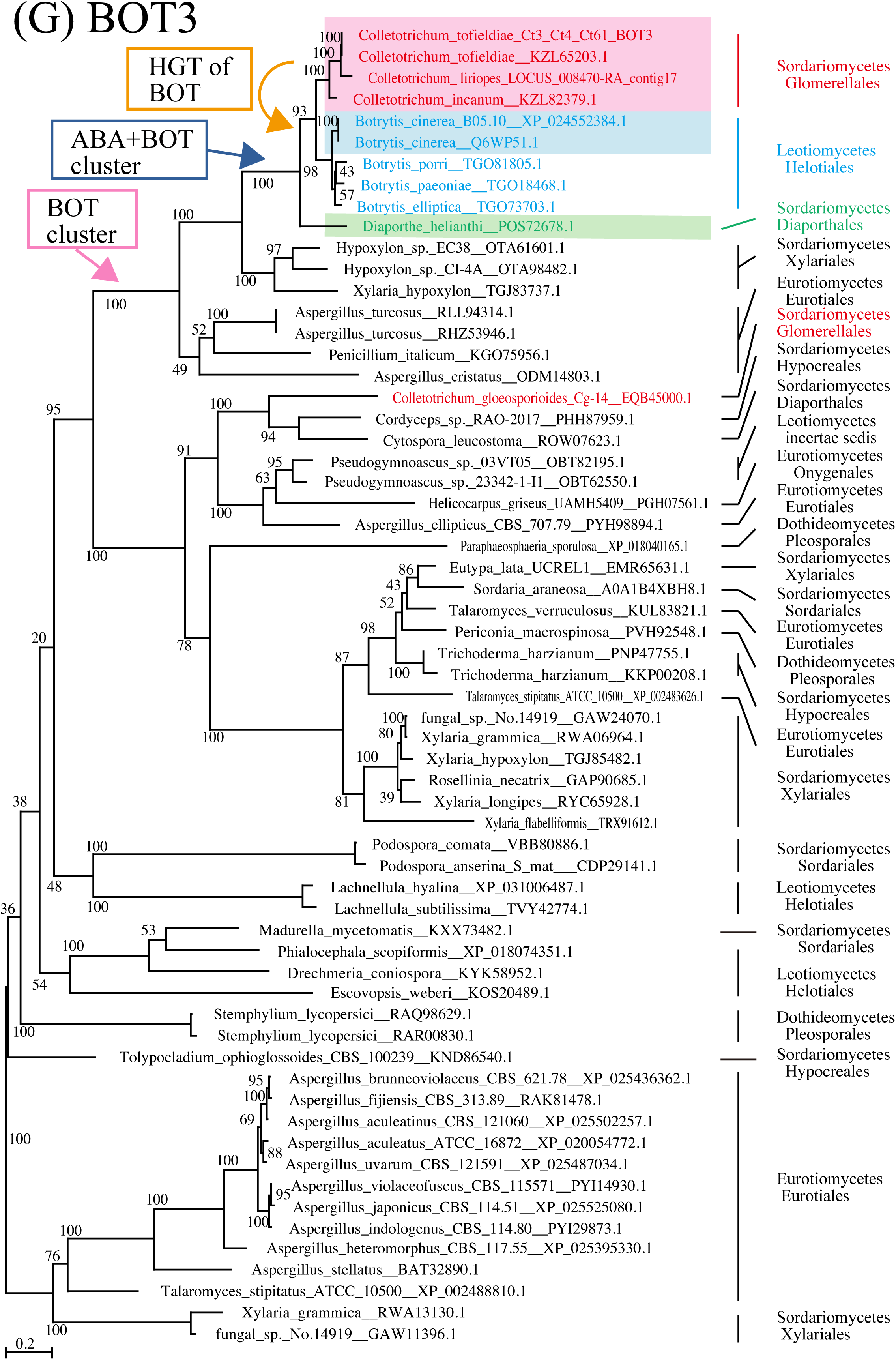
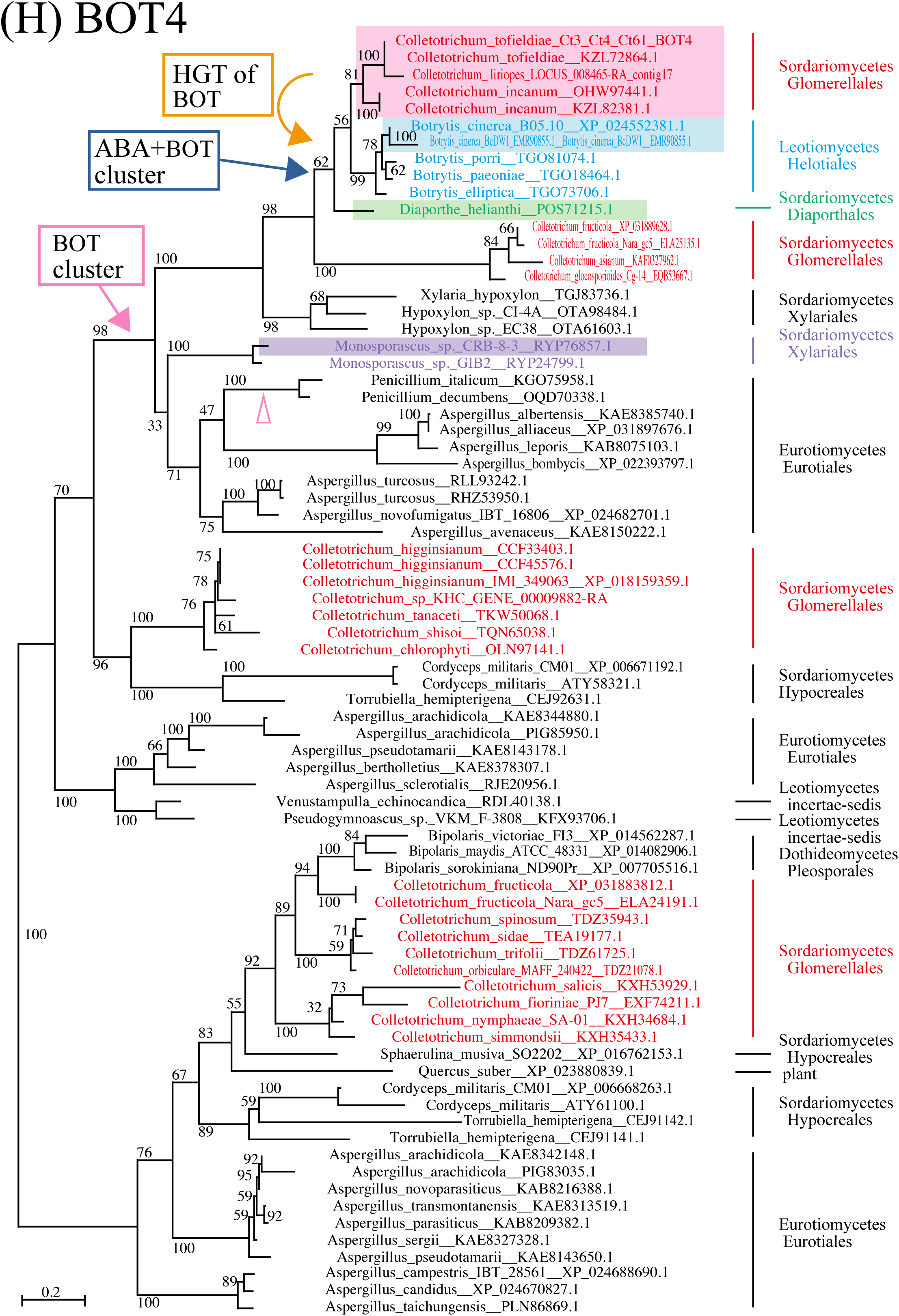
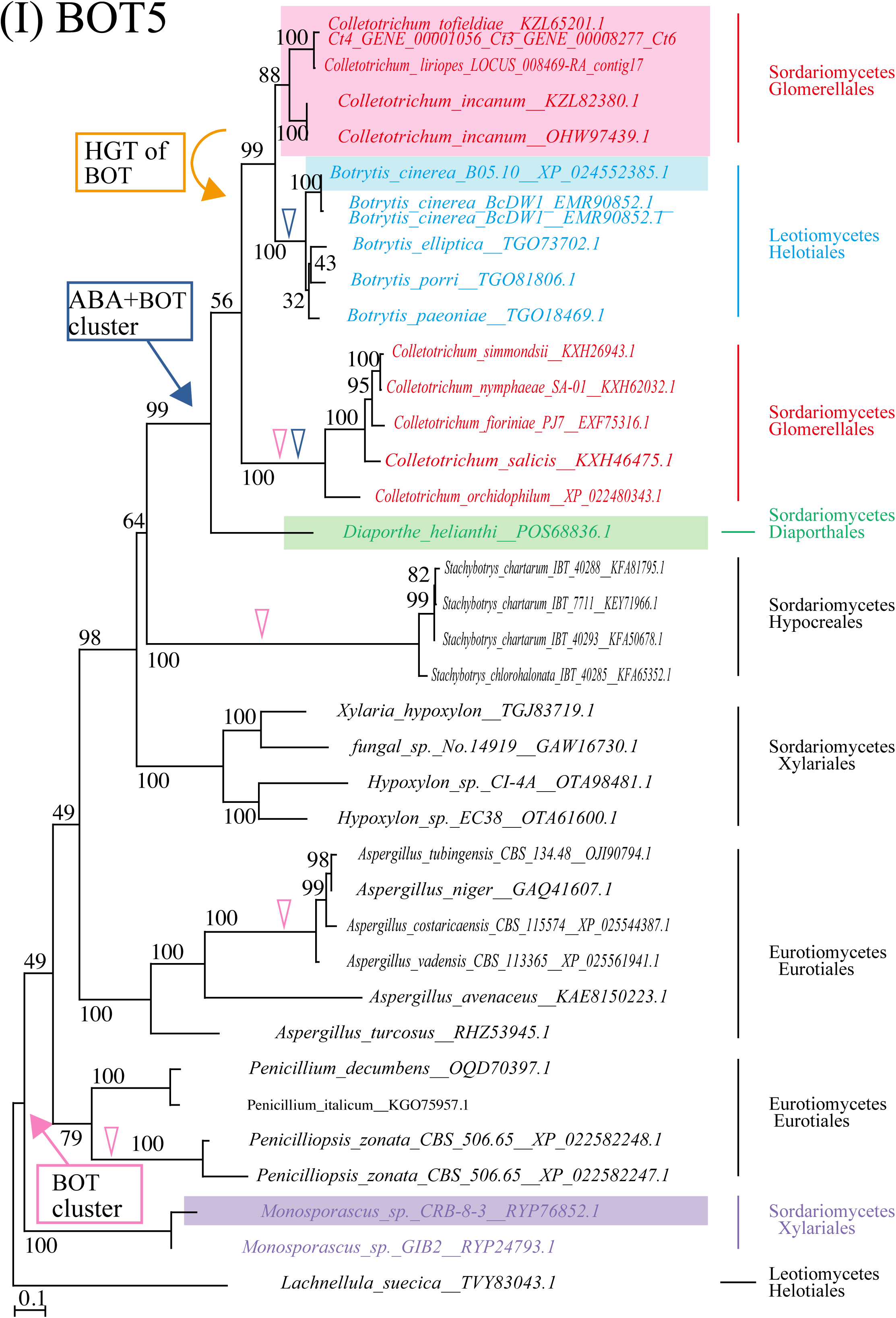
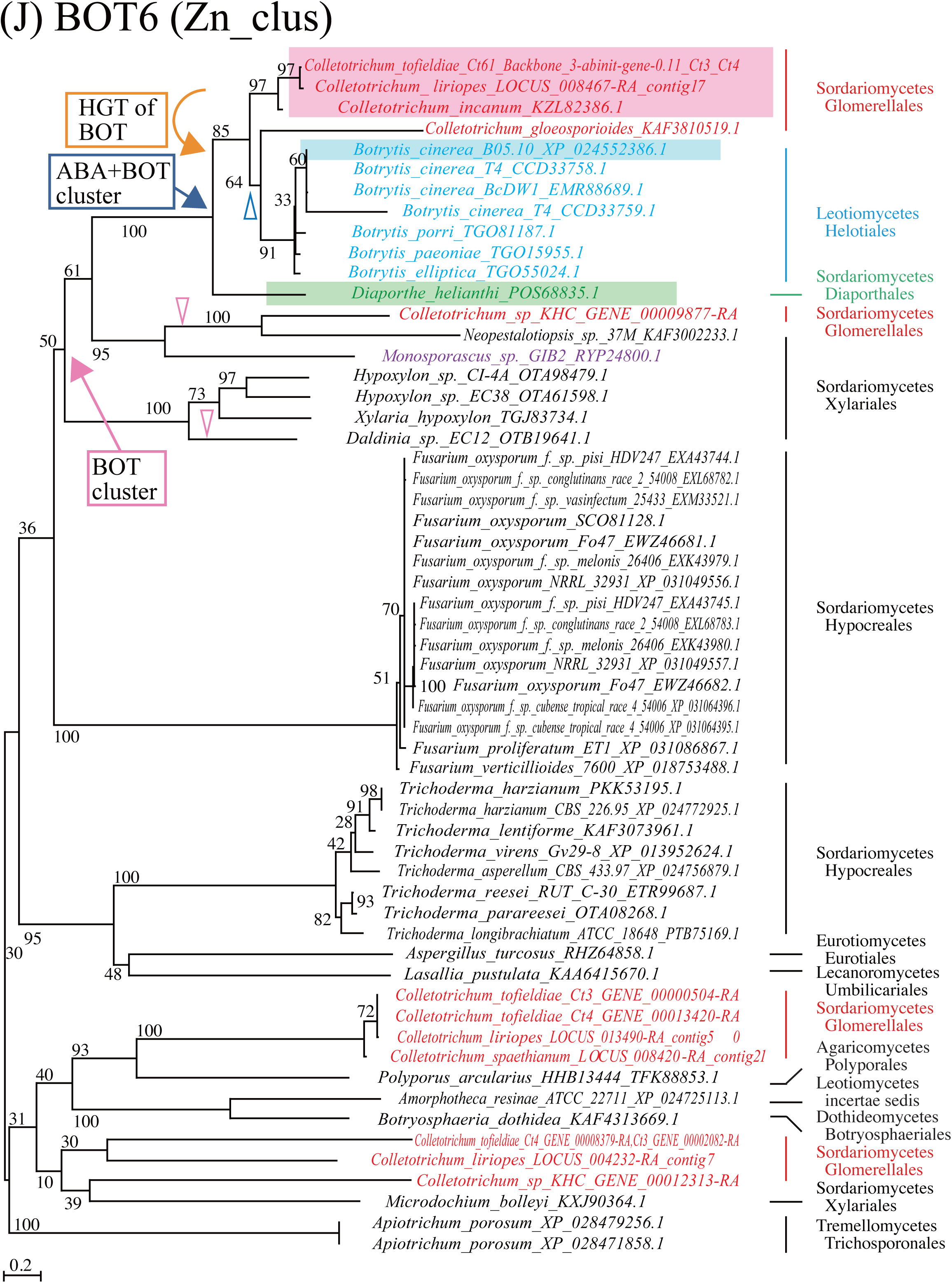
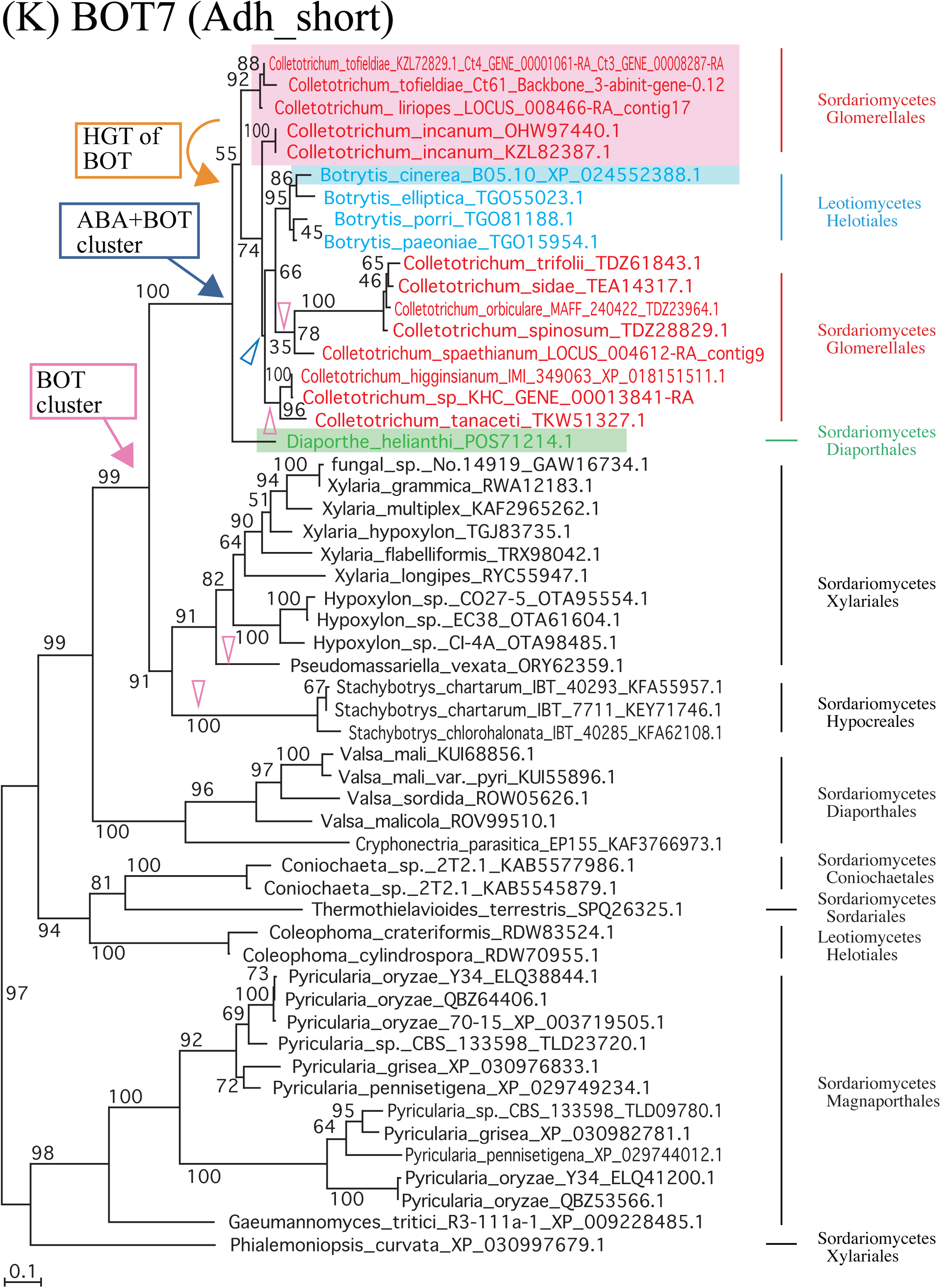
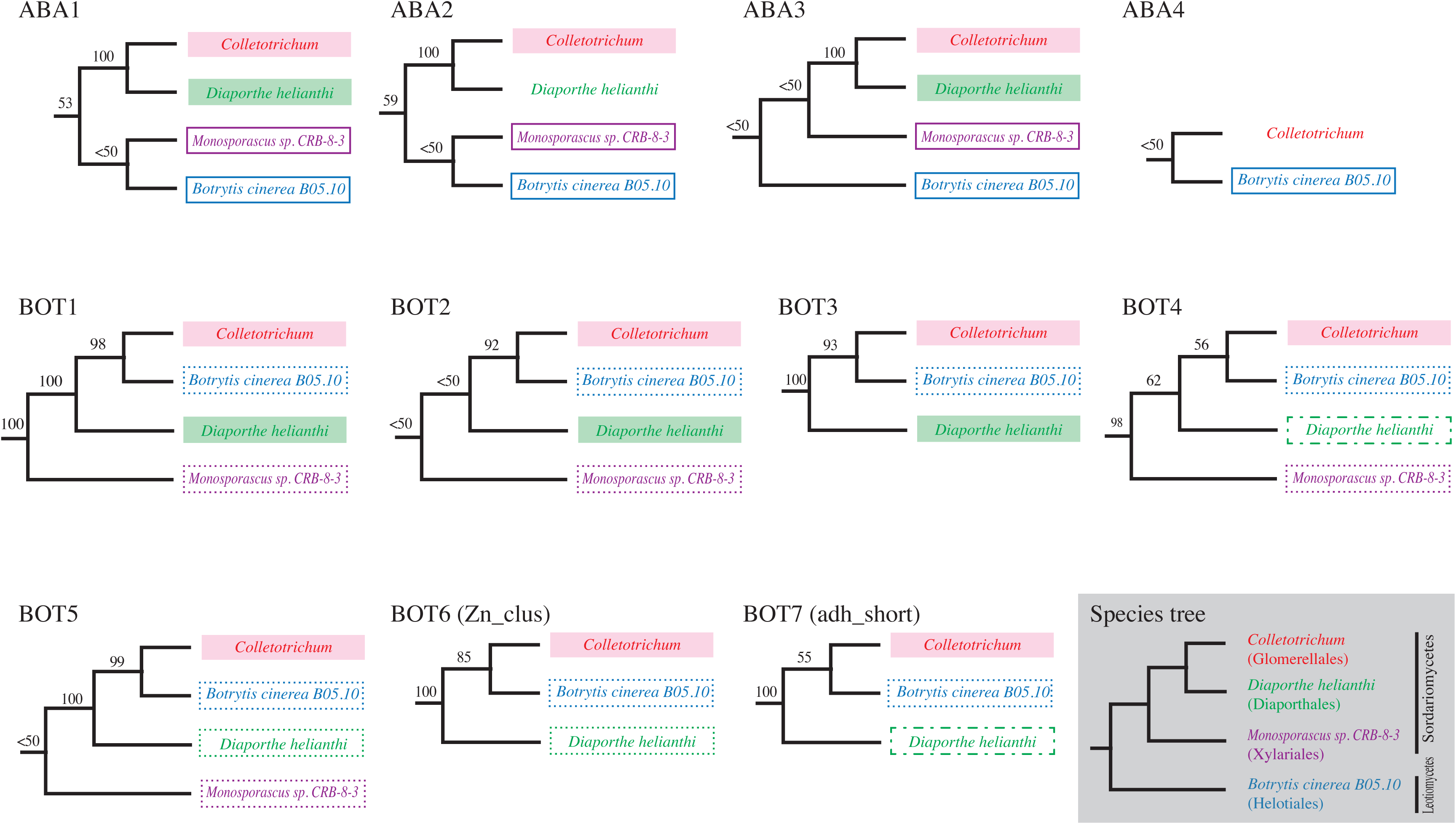
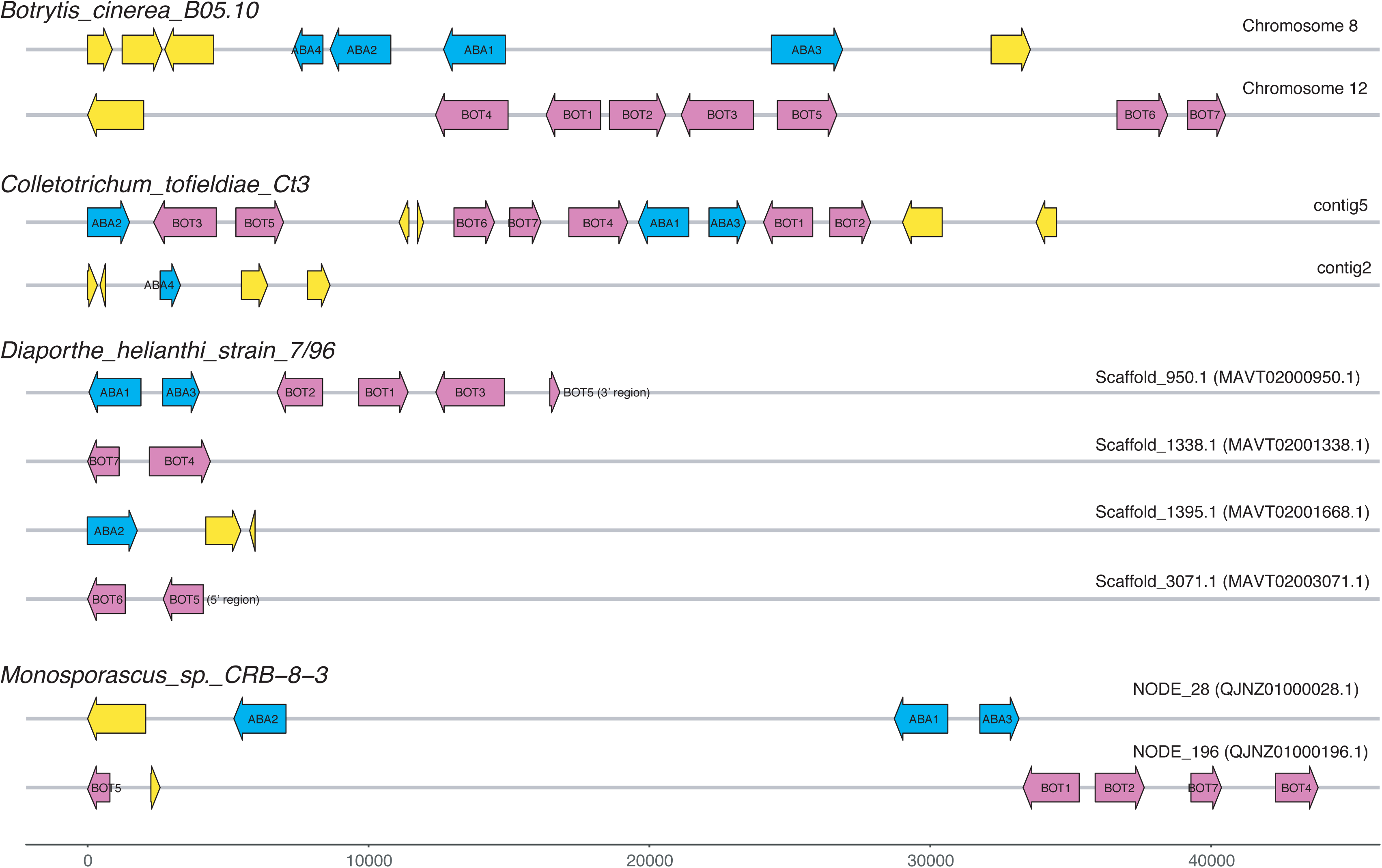
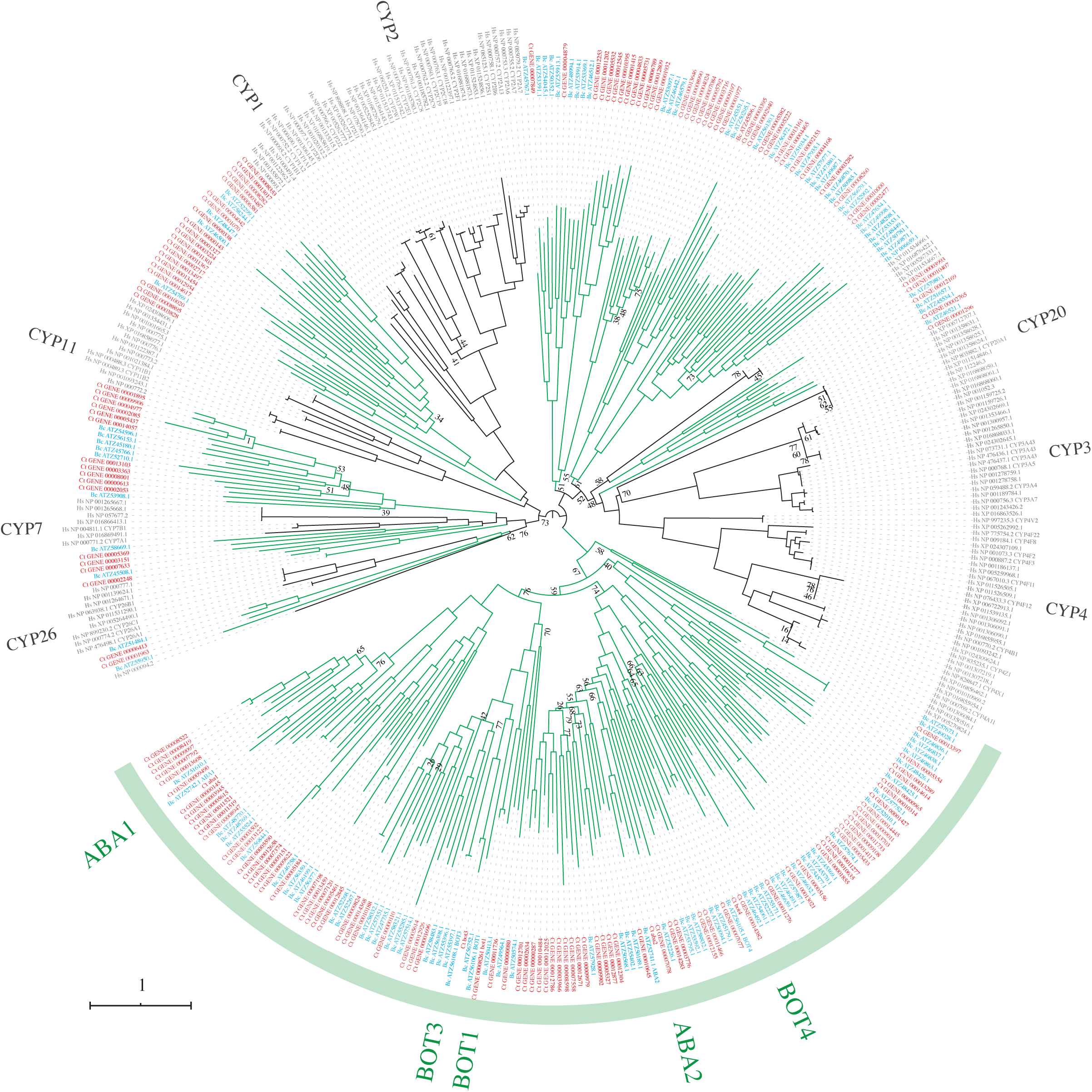
Fungal ABA and BOT biosynthesis genes are frequently transferred among plant-associated fungi via horizontal transfer. Related to Figure 4. **a–k.** Phylogenies of ABA and BOT genes using IQ-TREE version 1.6.11. Arrow indicates the origins of various gene clusters. Triangles with the same color as arrows, shows the deletion of clusters. Genes of *Colletotrichum*, *Botrytis*, *Diaporthe*, and *Monosporascus* are indicated by red, blue, green, and purple, respectively. Organism strains with both ABA and BOT gene clusters in their genome are labeled with the same color background as the letters. The bootstrap probability is shown on the branches. Scale bar represents substitutions per site. a–d *ABA1-4* genes. e–k *BOT1-7* genes. **l.** Phylogenetic relationship of ABA and BOT genes in the four organisms with both ABA and BOT clusters in their genome. The relationship and branch support are based on (a)–(k). The species tree of the four organisms is shown in the rectangle with a gray background. Organisms with an ABA+BOT cluster are labeled with red or green backgrounds. Organism names of each gene on the same molecule (but not on the ABA+BOT cluster) are boxed with the same solid or dotted line. **m.** Schematic drawing of genetic structures of ABA and BOT gene clusters. Genes on the same molecule (chromosome or scaffold) are drawn on a line side by side. Color coding: blue, ABA genes; red, BOT genes; yellow, other genes. Bar indicates base pairs. **n.** Phylogenetic tree of genes in cytochrome C P450 families. 414 genes of *C. tofieldiae Ct3*, *B. cinerea* B05.10, and *H. sapiens* were used for the ML analysis using IQ-TREE v. 1.6.11. Branches with green and black indicate fungi and human lineages, respectively. Red letters indicate *C. tofieldiae* genes, blue indicates *B. cinerea*, and gray indicates *H. sapiens*. ABA, BOT, and CYP genes are highlighted. All ABA and BOT genes are included in a large monophyletic clade of fungi (green arc). The number at each node corresponds to the bootstrap values supported from the ML analysis using IQ-TREE (only less than 80% were indicated). Scale bar represents substitutions per site. The tree’s root is based on the option of root mid-point of iTOL v. 5.6.

**Supplementary Figure 5.**
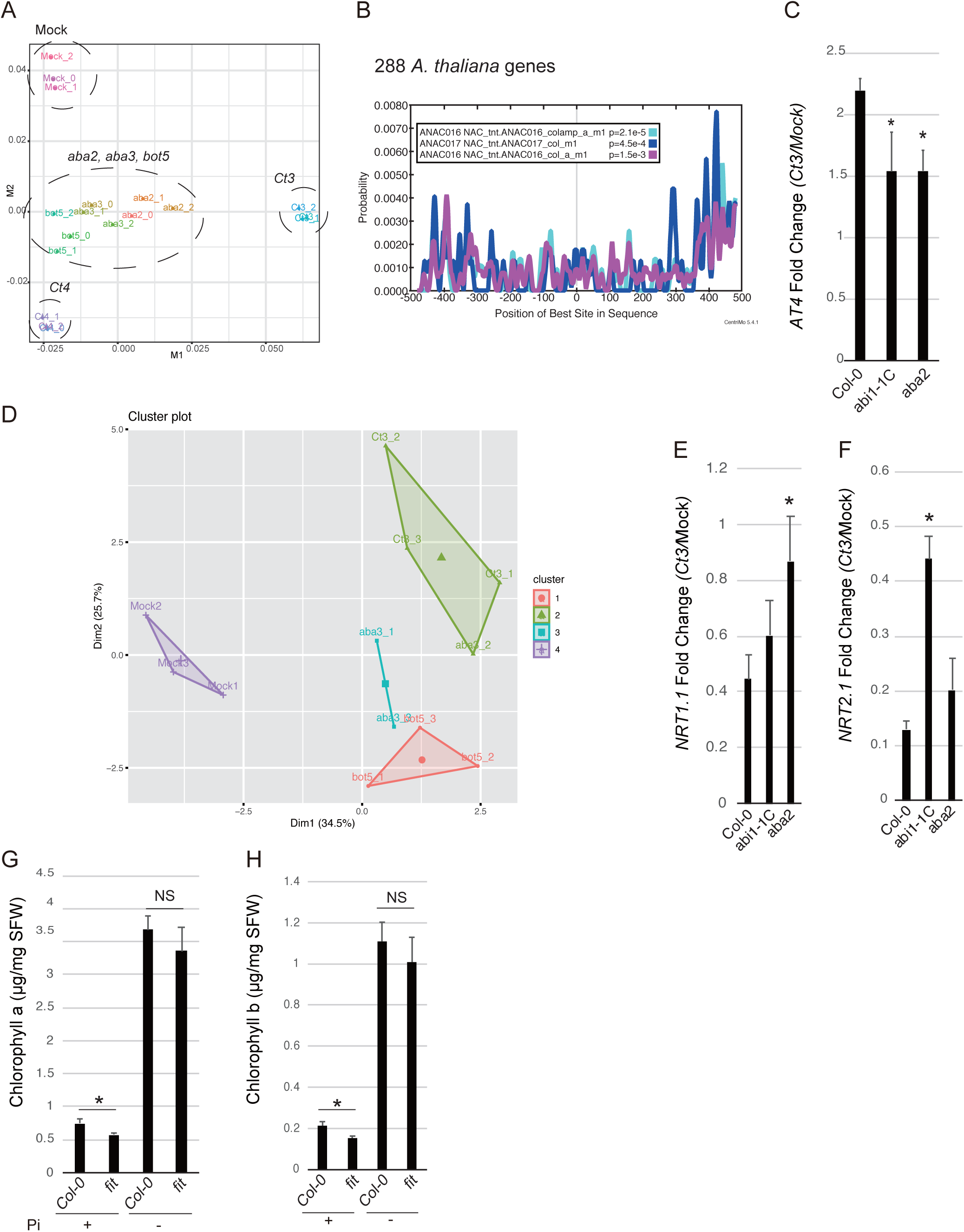
Fungal ABA and BOT biosynthesis genes suppress the expression of nutrient uptake-related *A. thaliana* genes while activating PSR genes. Related to Figure 5. **a.** MDS chart generated from plant transcriptome profile during association with mock (M), *Ct3*, *aba2*, *aba3, bot5,* and *Ct4* under low Pi. The three independent biological replicates are shown. **b.** Motif analysis against 1,000 bp upstream sequences of 288 upregulated *A. thaliana* genes during root colonization by *Ct3* compared to *aba2*, *aba3, bot5,* and *Ct4*. **c.** *AT4* gene fold change under low Pi. The relative expression of *AT4* estimates the fold change (*Ct3*/Mock). Asterisks indicate significantly different means between Col-0 and the corresponding mutants (SD, n = 3, p < 0.05, two-tailed t-test). **d.** PCA analysis of shoot mineral nutrient composition (ionome) in plants grown in normal Pi treated with either mock, *Ct3*, *aba2*, *aba3*, or *bot5*. The samples were separated into four different clusters. **e.** *NRT1.1* gene fold change under low Pi. Asterisks indicate significantly different means between Col-0 and the corresponding mutants (SD, n = 3, p < 0.05, two-tailed t-test). **f.** *NRT2.1* gene fold change under low Pi. Asterisks indicate significantly different means between Col-0 and the corresponding mutants (SD, n = 3, p < 0.05, two-tailed t-test). **g.h** Quantitative data of plant shoot chlorophyll a or b in Col-0 and *fit* under normal and low Pi. Asterisks indicate significantly different means between Col-0 and *fit* plants (SD, p < 0.01, two-tailed t-test).

**Table S1:**
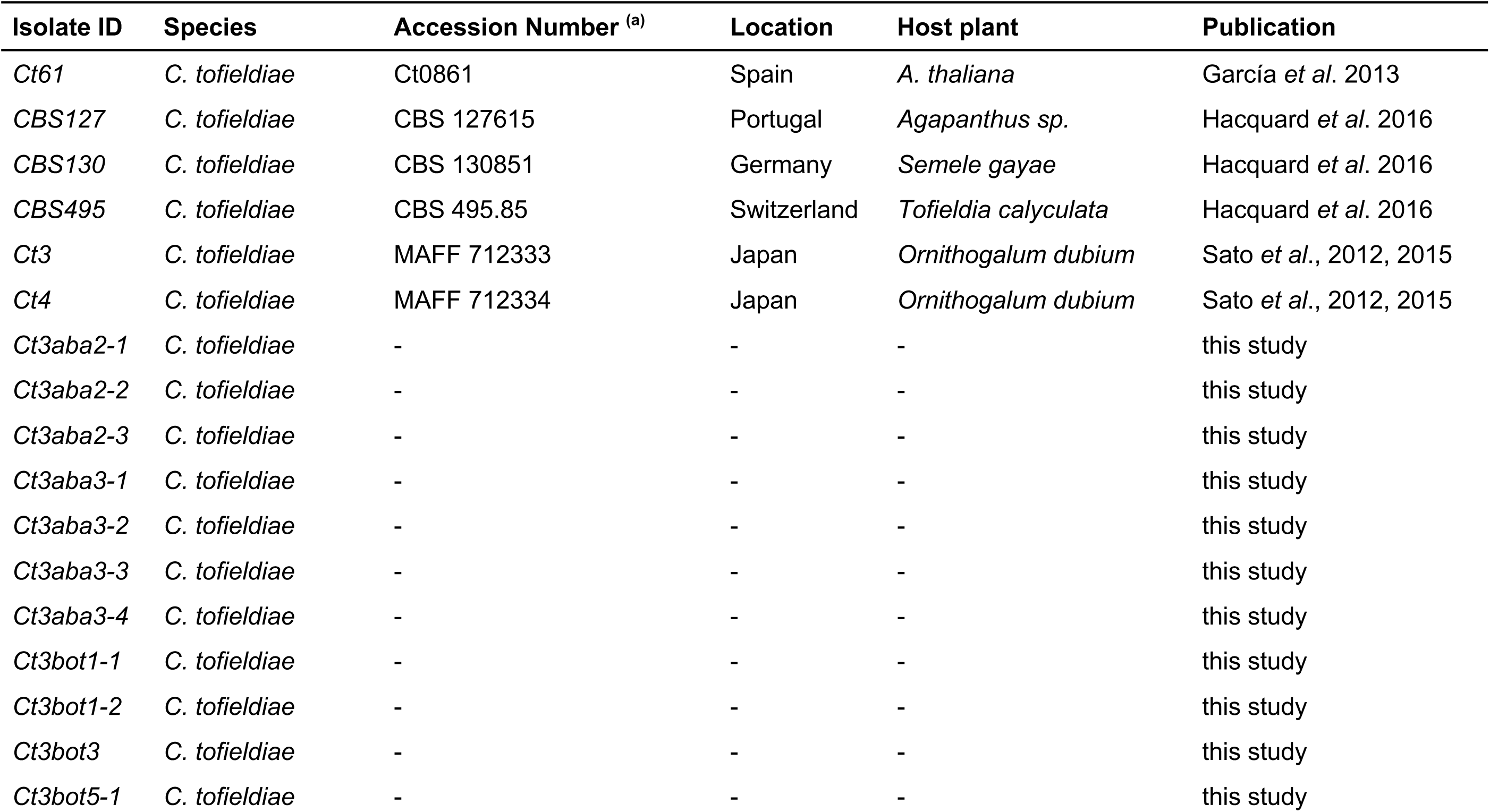
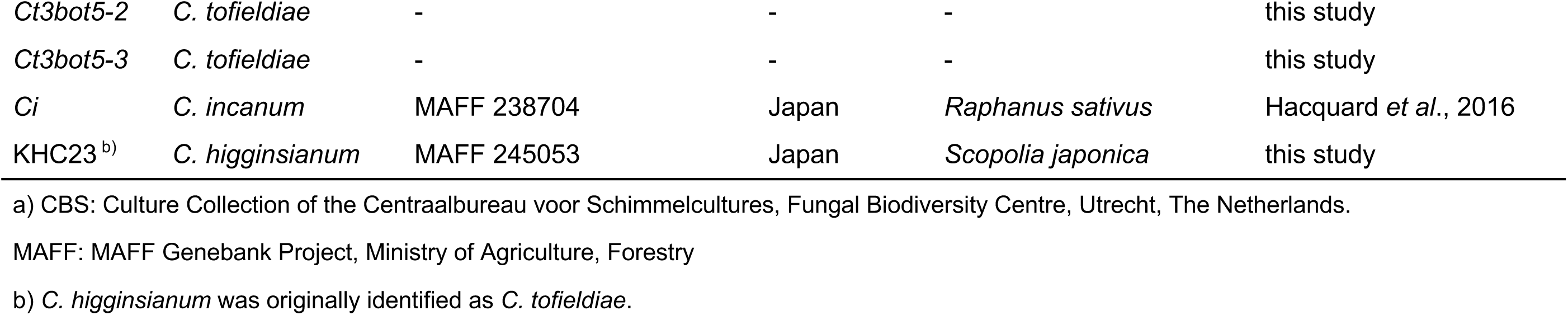
Background information regarding Colletotrichum tofieldiae and KHC23 (KHC) isolates or the Ct3 mutants selected or generated in this study.

**Table S2:** Genome sequencing and assembly statistics of C. tofieldiae isolates and KHC.

| Assembly Statistics | Illumina +454 | Pacbio | Pacbio + Illumina | Pacbio | Pacbio |
| --- | --- | --- | --- | --- | --- |
| Isolate | *Ct0861*<sup>a</sup> | *Ct0861* (PacBio) | *Ct4* | *Ct3* | KHC |
| Genome (bp) | 52836184 | 52991954 | 53983402 | 54252643 | 49069799 |
| Number of Scaffolds | 1048 | 28 | 27 | 19 | 27 |
| Largest scaffold (bp) | 861708 | 6294542 | 6028009 | 6503204 | 5997240 |
| Smallest scaffold (bp) | 201 | 19730 | 323670 | 8543 | 4021 |
| Genome N50 (bp) | 209340 | 4211846 | 3719715 | 6011218 | 4389193 |
| GC content (%) | 45.7 | 50.3 | 52.1 | 48.6 | 50.6 |
| Number of CDS | 12501 | 15617 | 14573 | 14916 | 14359 |
| Number of tRNA | 389 | 390 | 370 | 418 | 380 |
| Number of rRNA | 73 | 124 | 70 | 86 | 113 |
| Candidate secreted proteins | 1391 | 1397 | 1446 | 1481 | 1426 |
| Candidate effector proteins | 430 | 490 | 496 | 507 | 475 |
| Carbohydrate-active enzymes | 720 | 710 | 729 | 733 | 679 |
| Candidate secondary metabolites | 59 | 59 | 59 | 56 | 77 |
| BUSCO genome mode | 97.6 | 88.5 | 97.4 | 97.2 | 93.6 |
| BUSCO protein mode | 94.7 | 86 | 95 | 94.4 | 90.5 |

a) This genome assembly has been reported in Hacquard *et al.*, 2016.

**Table S3:** List of *A. thaliana* genes whose expression was significantly influenced between sample_1 and sample_2 (q (FDR) < 0.05) at 10 days post-inoculation (dpi). P = normal Pi. mp = low Pi. Treatment: Mock, *Ct61*, *Ct4*, *Ct3*, or KHC.

**Table S4:** Gene ontology analyses for 758 A. thaliana genes specifically and significantly upregulated during root colonization by pathogenic Ct3 compared with beneficial Ct61 and Ct4 at 10 dpi (log2FC >1, q (FDR) < 0.05). Genes were separated into three different clusters by k-means.

**Table S5:** List of *Ct3* genes significantly influenced during *A. thaliana* root colonization between 10 dpi and 24 dpi (q (FDR) < 0.05). P = normal Pi. mp = low Pi. e = 10 dpi. L = 24 dpi.

**Table S6:**
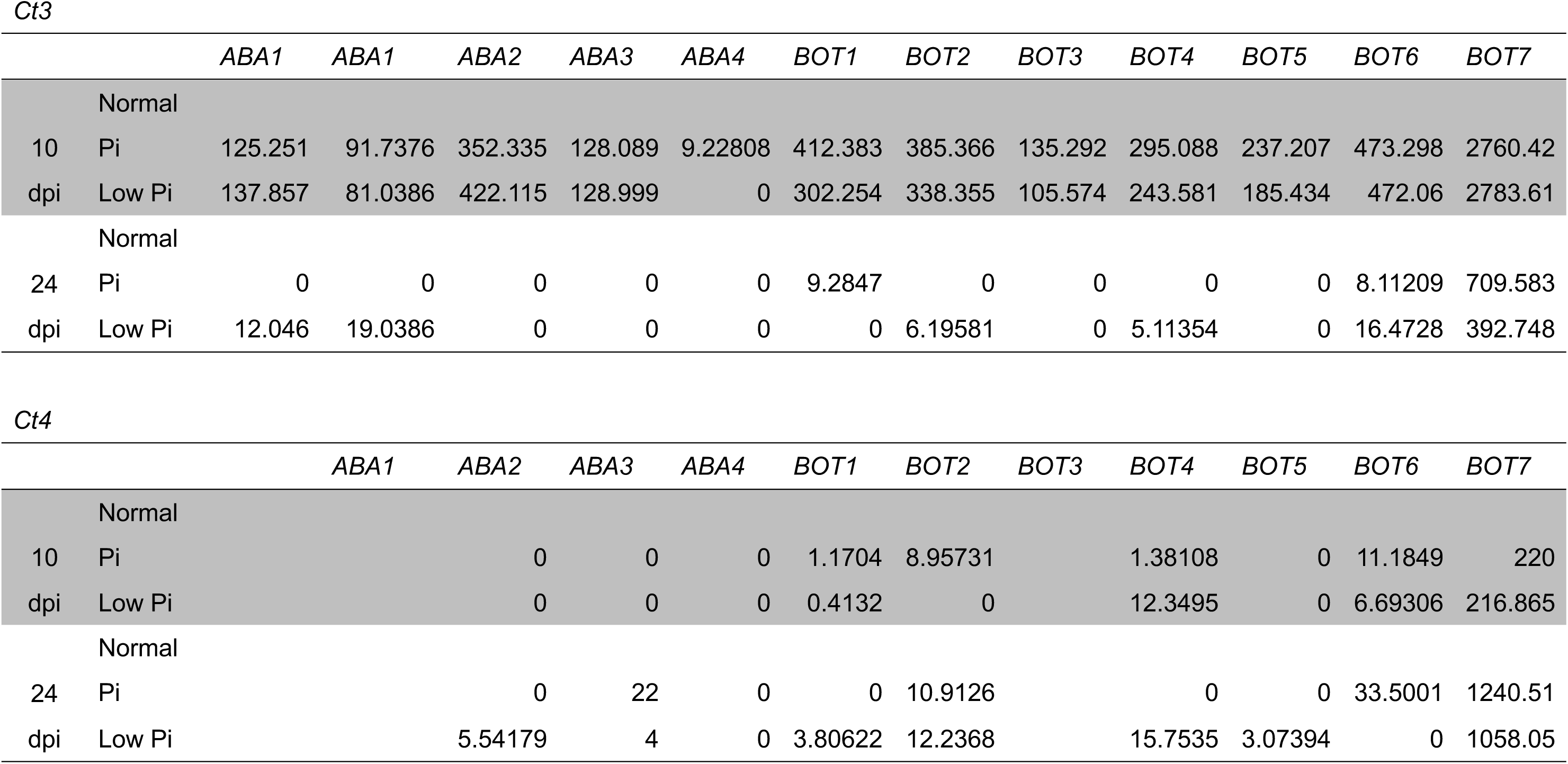
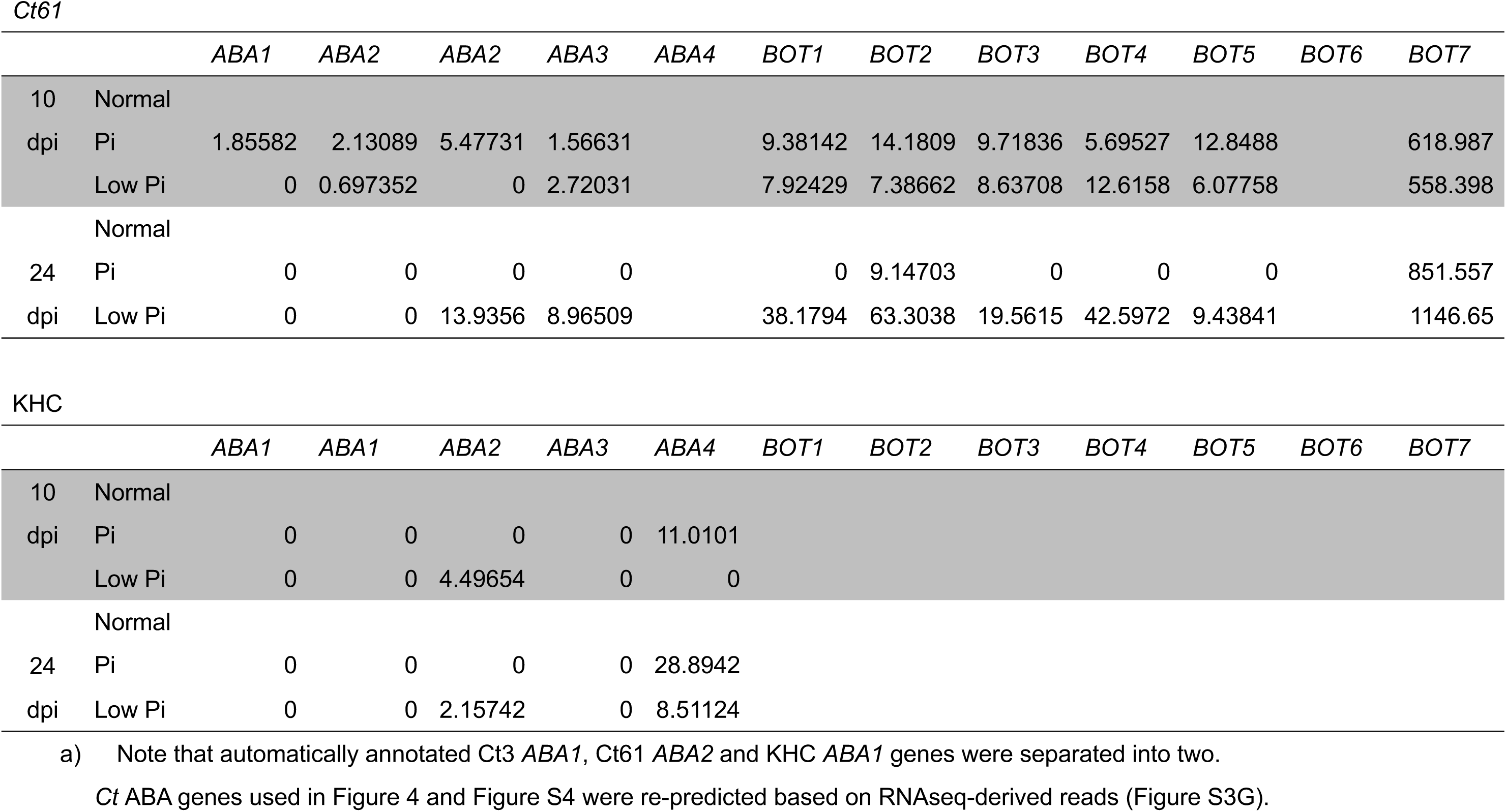
Expression of Ct3, Ct4, Ct61 or KHC ABA and BOT genes during root colonization. Values represent fpkm. This value was calculated based on Ct3, Ct4, Ct61, or KHC genome assemblies.

**Table S7:** ABA (1-4) or BOT(1-7) homologous genes used as the query of BLASTP search against GenBank NR database.

**Table S8:** Amino acid sequences used in Figures 4 and S4.

**Table S9:** List of *A. thaliana* genes whose expression was significantly influenced between sample_1 and sample_2 (q (FDR) < 0.05) at 10 days post-inoculation (dpi) under low Pi. Treatment: Mock, *Ct3*, *aba2*, *aba3*, *bot5*, or *Ct4*.

**Table S10:** A total of 288 A. thaliana genes were significantly induced by root colonization with Ct3WT than those with aba2, aba3, bot5, and Ct4 (log2FC >1, q (FDR) < 0.05).

**Table S11:** 375 *A. thaliana* genes were significantly suppressed by root colonization with *Ct3*WT compared to those with *aba2*, *aba3*, *bot5*, and *Ct4* (log2FC <1, q (FDR) < 0.05).

**Table S12:**
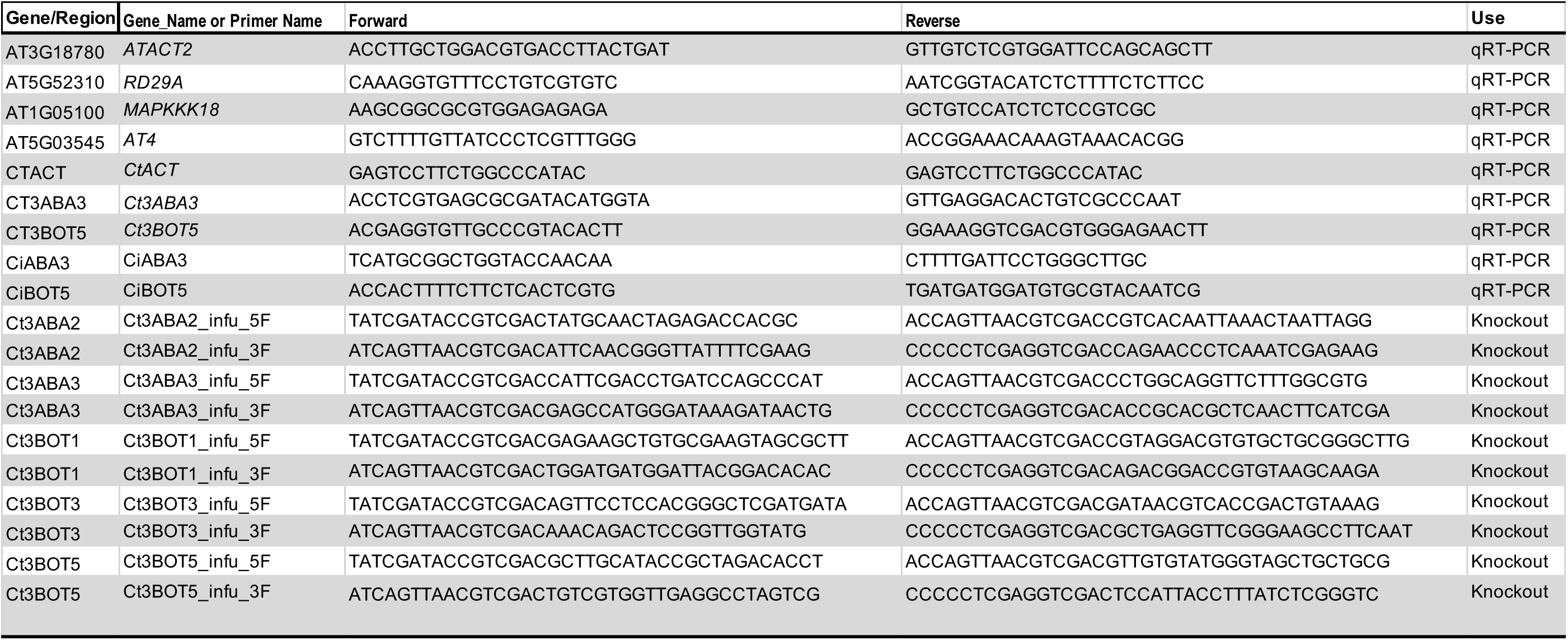
Primers used in this study.

CLUSTAL format alignment by MAFFT (v7.471)

**Figure.**
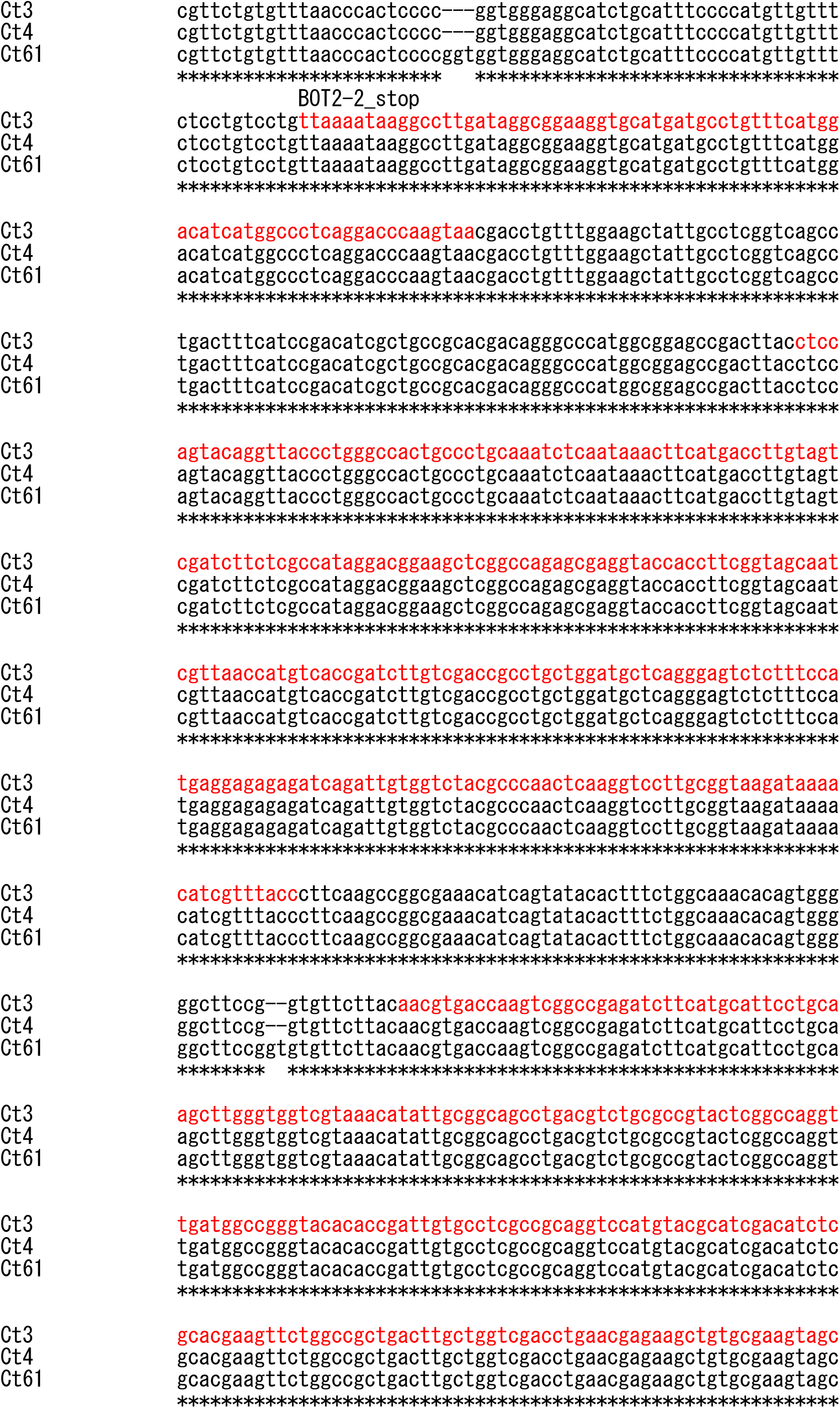
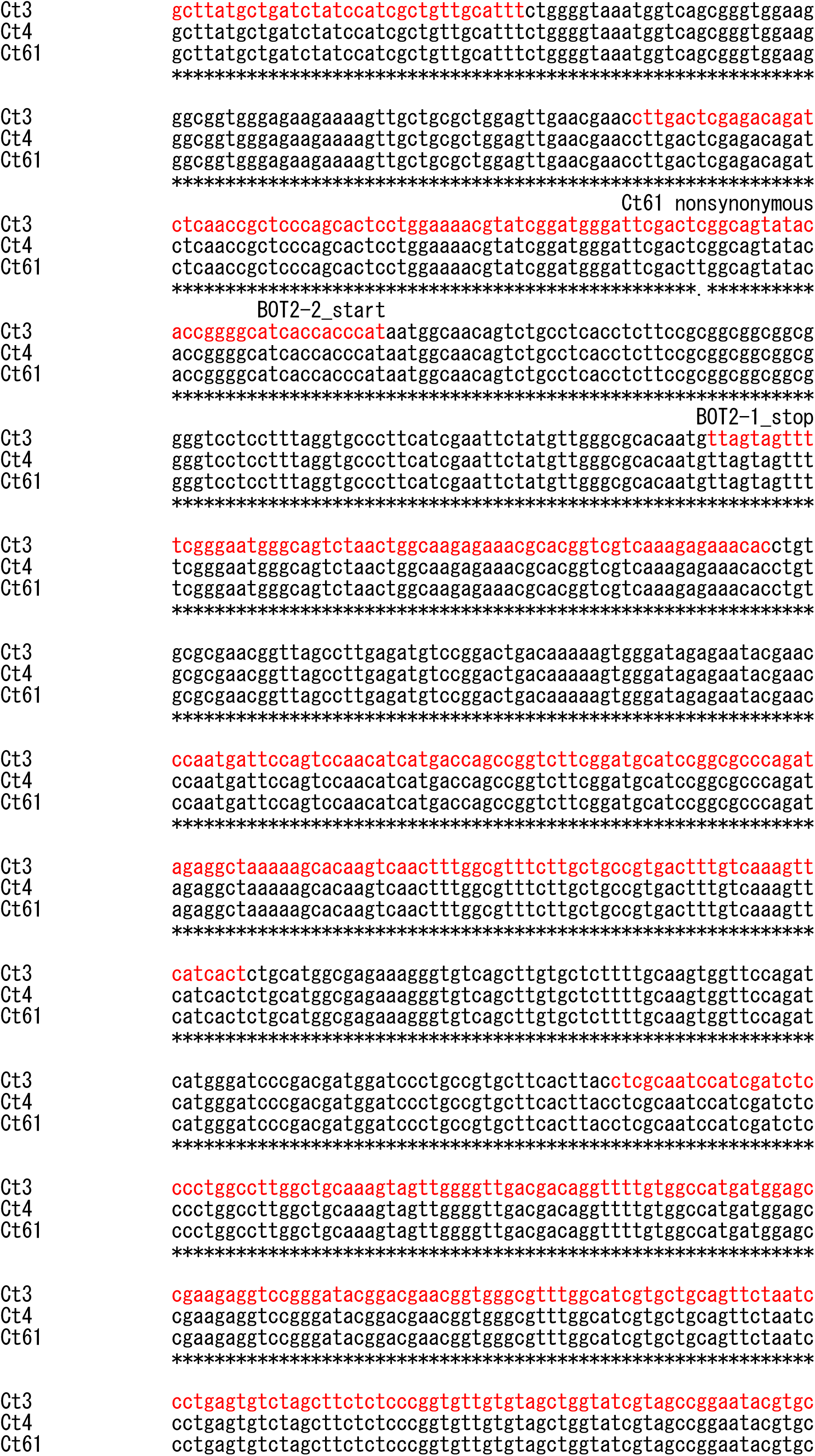
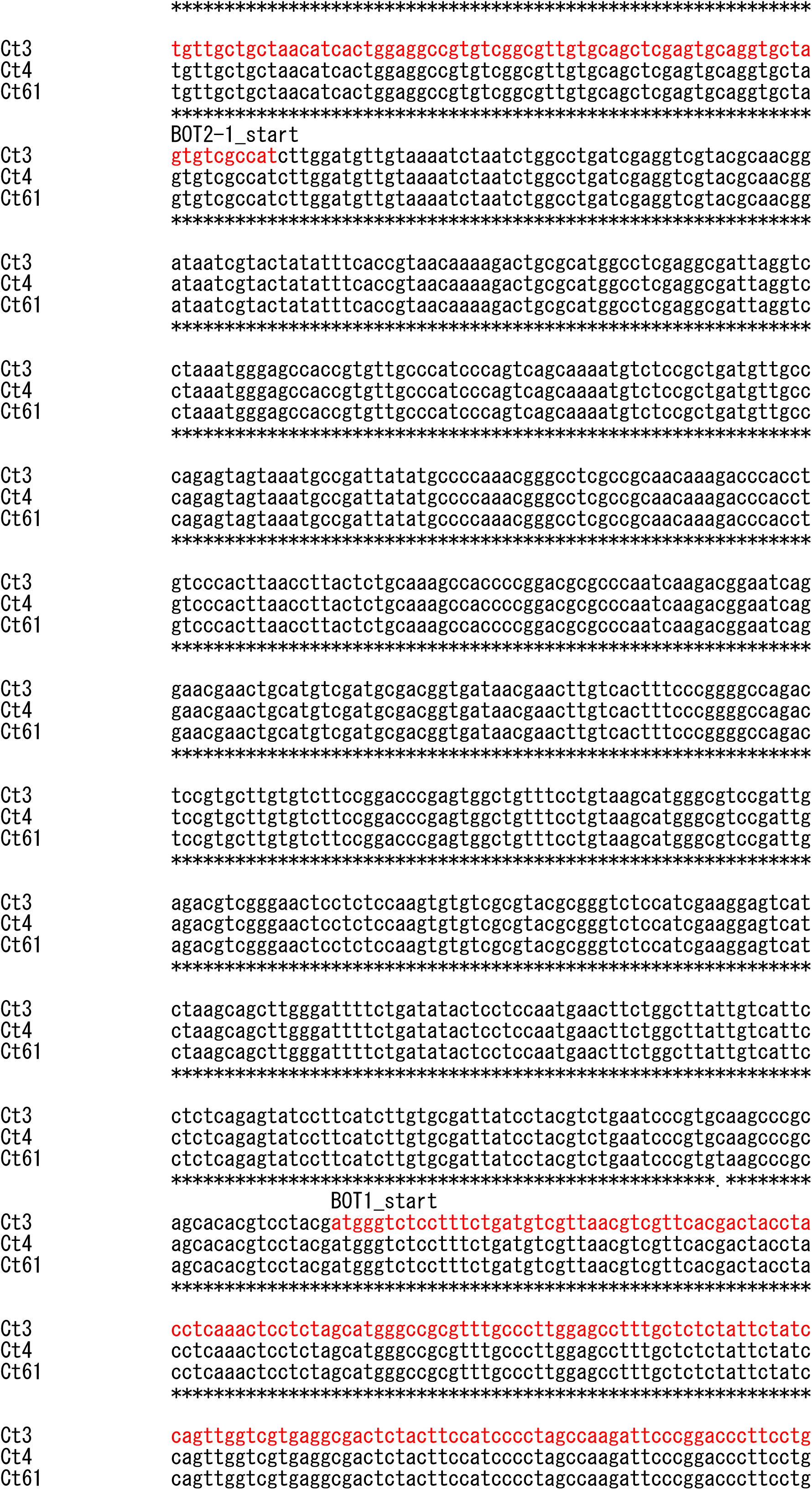
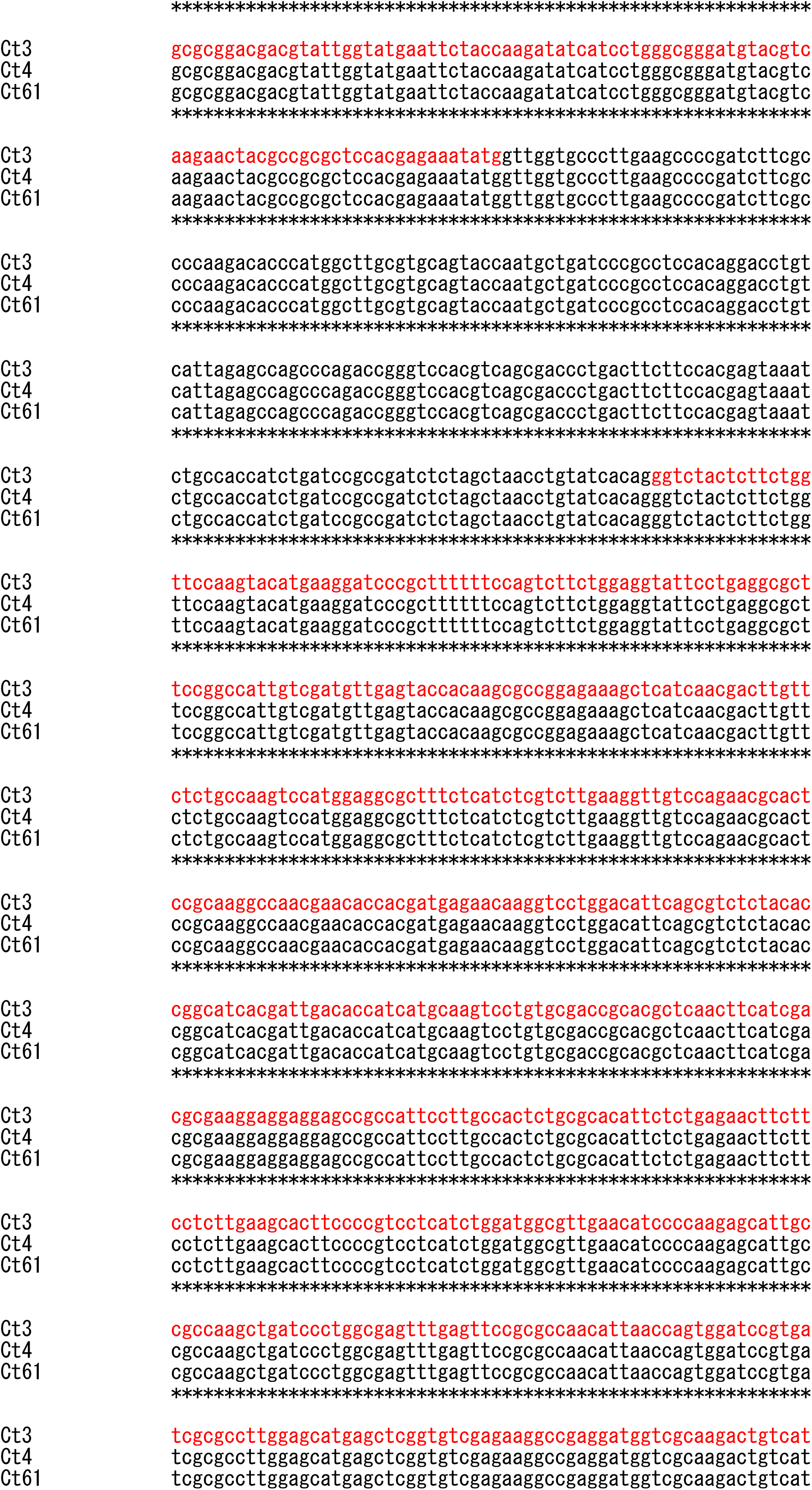
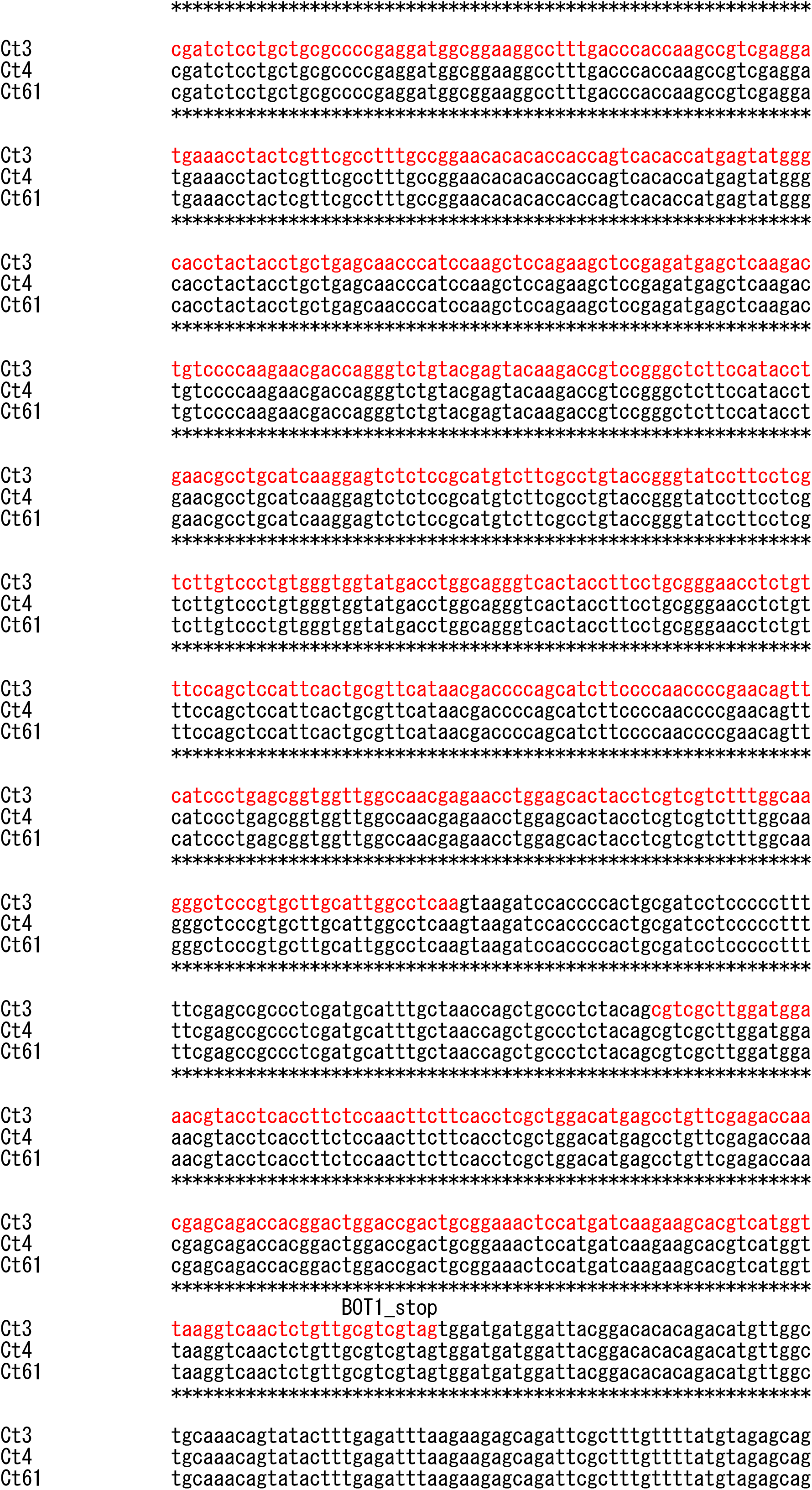
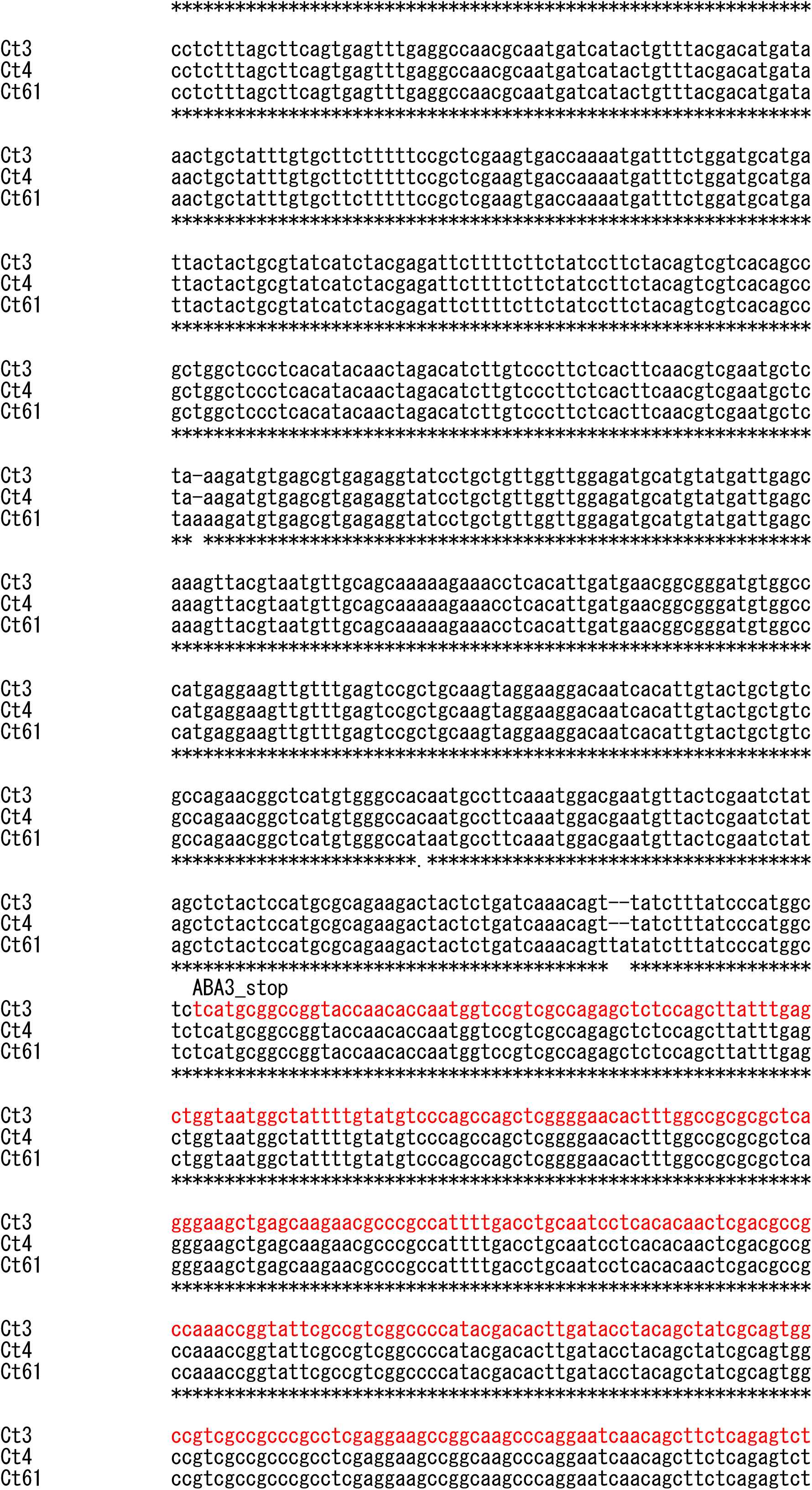
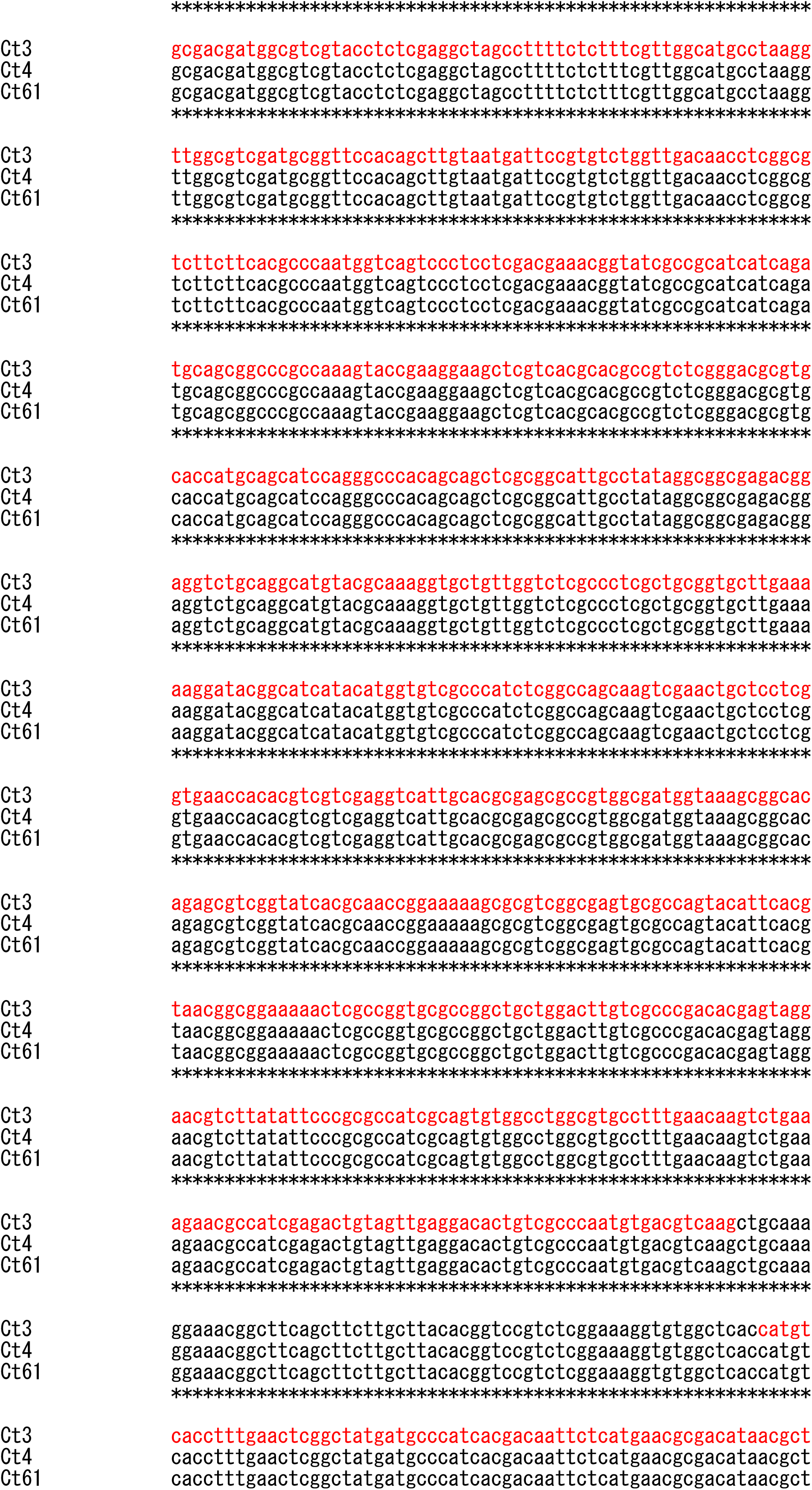
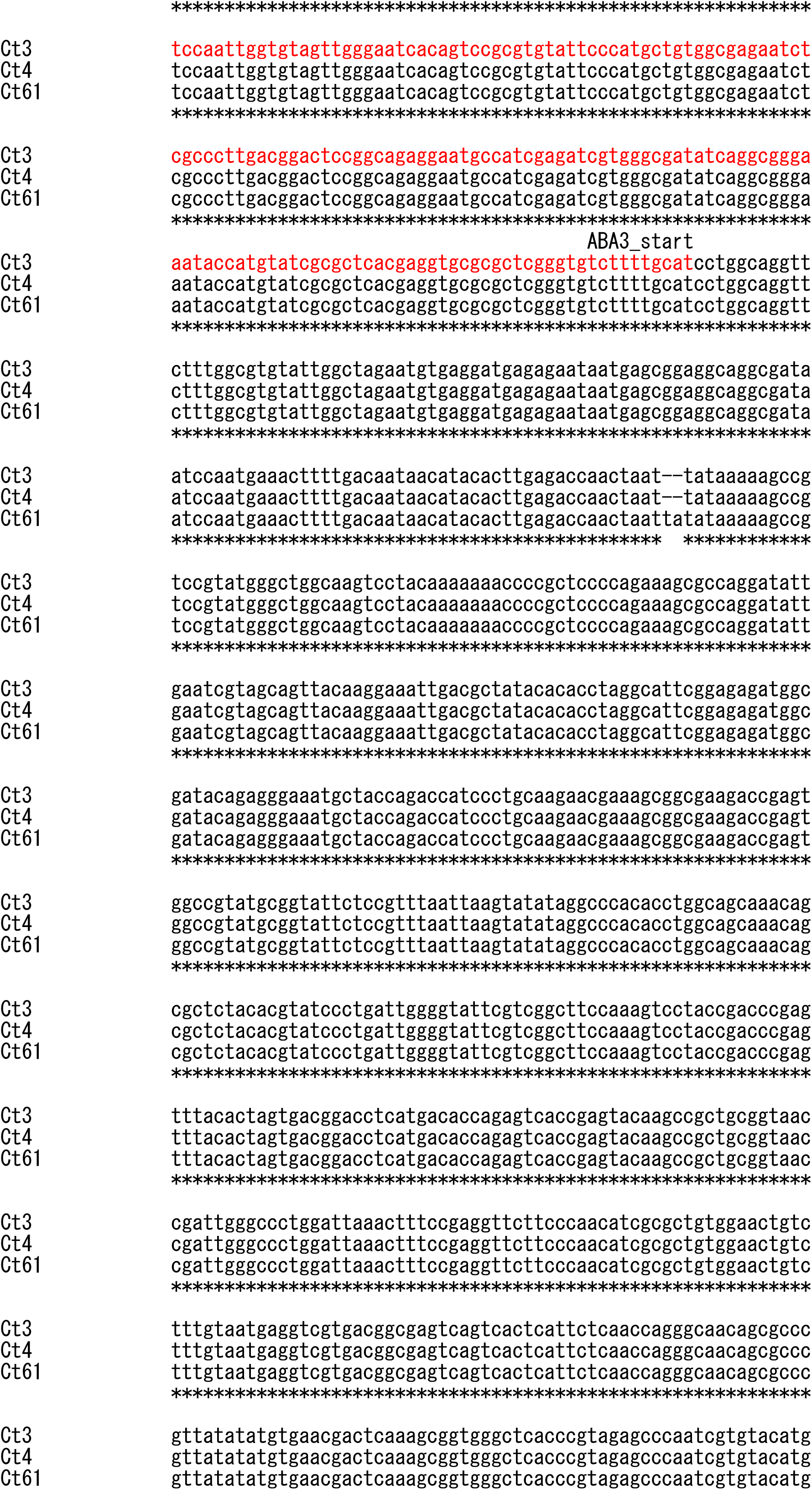
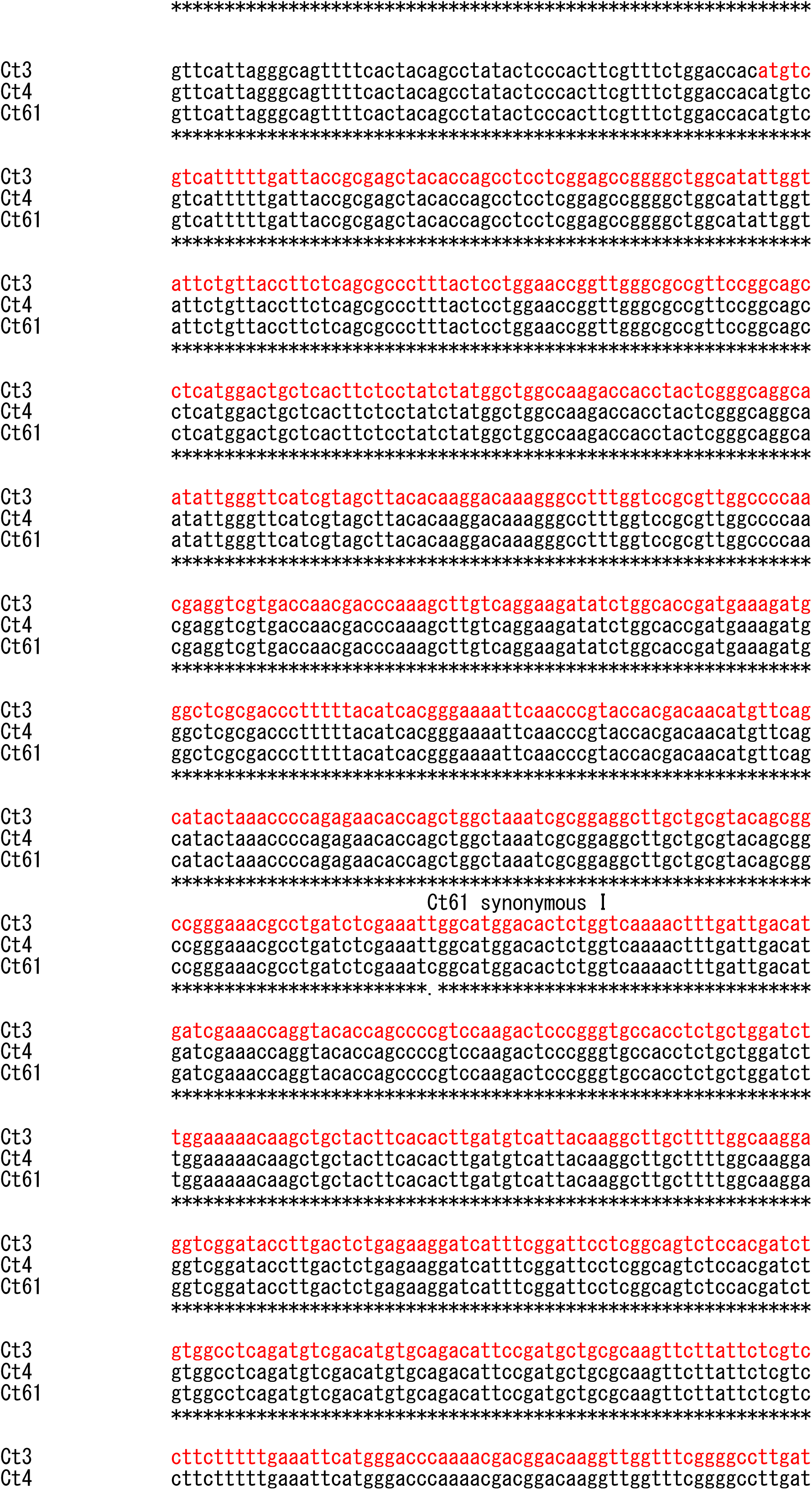
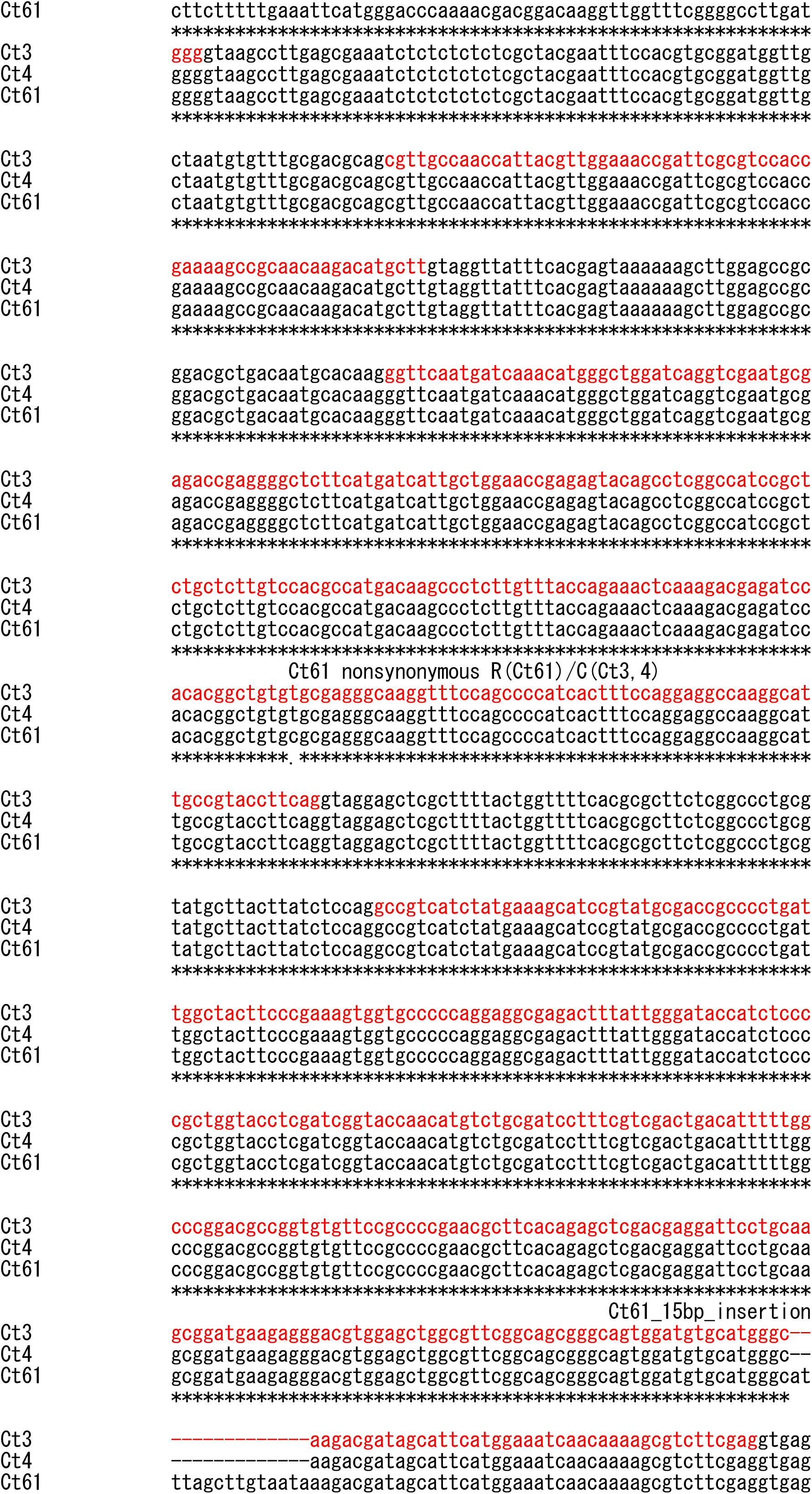
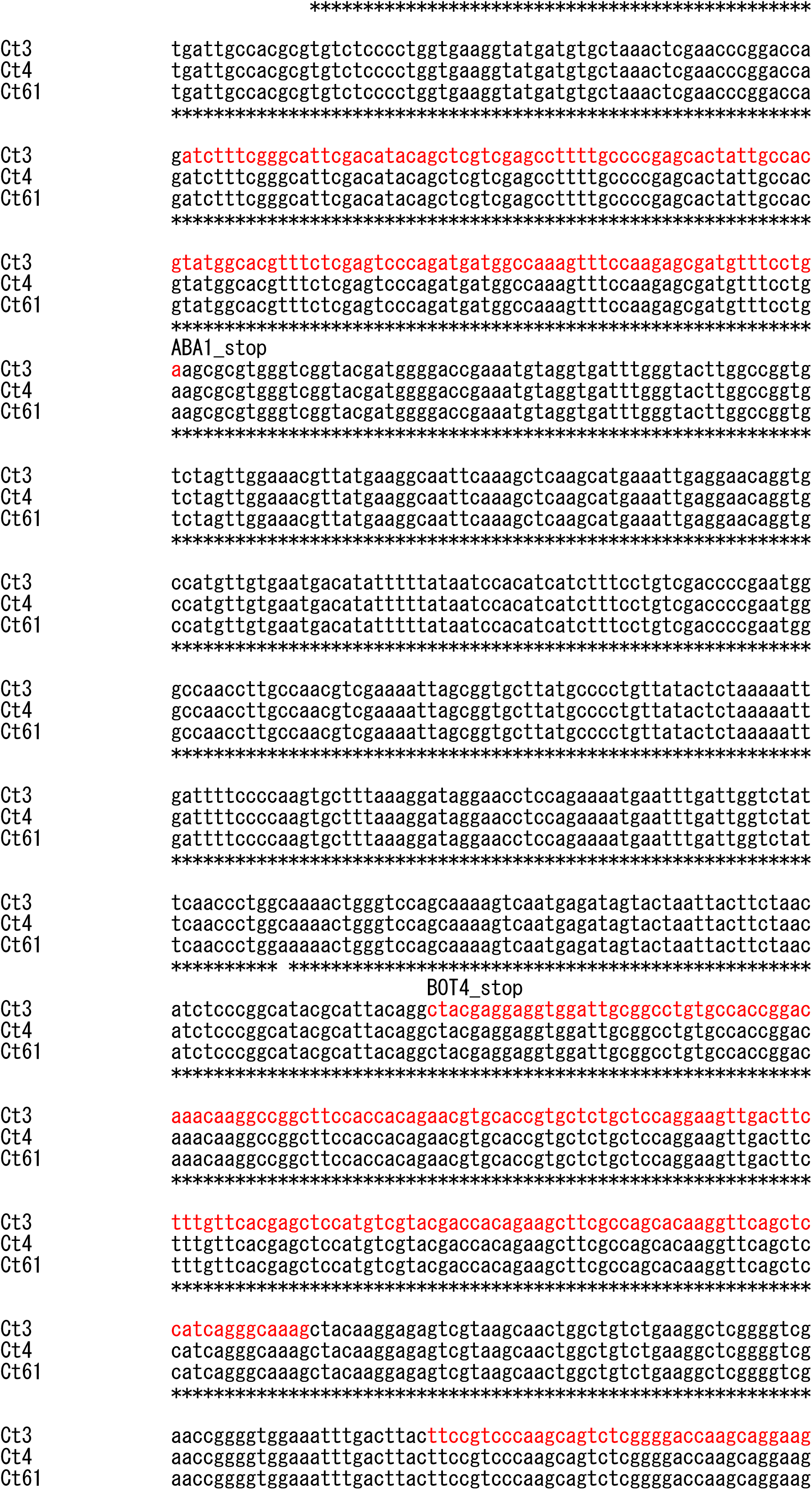
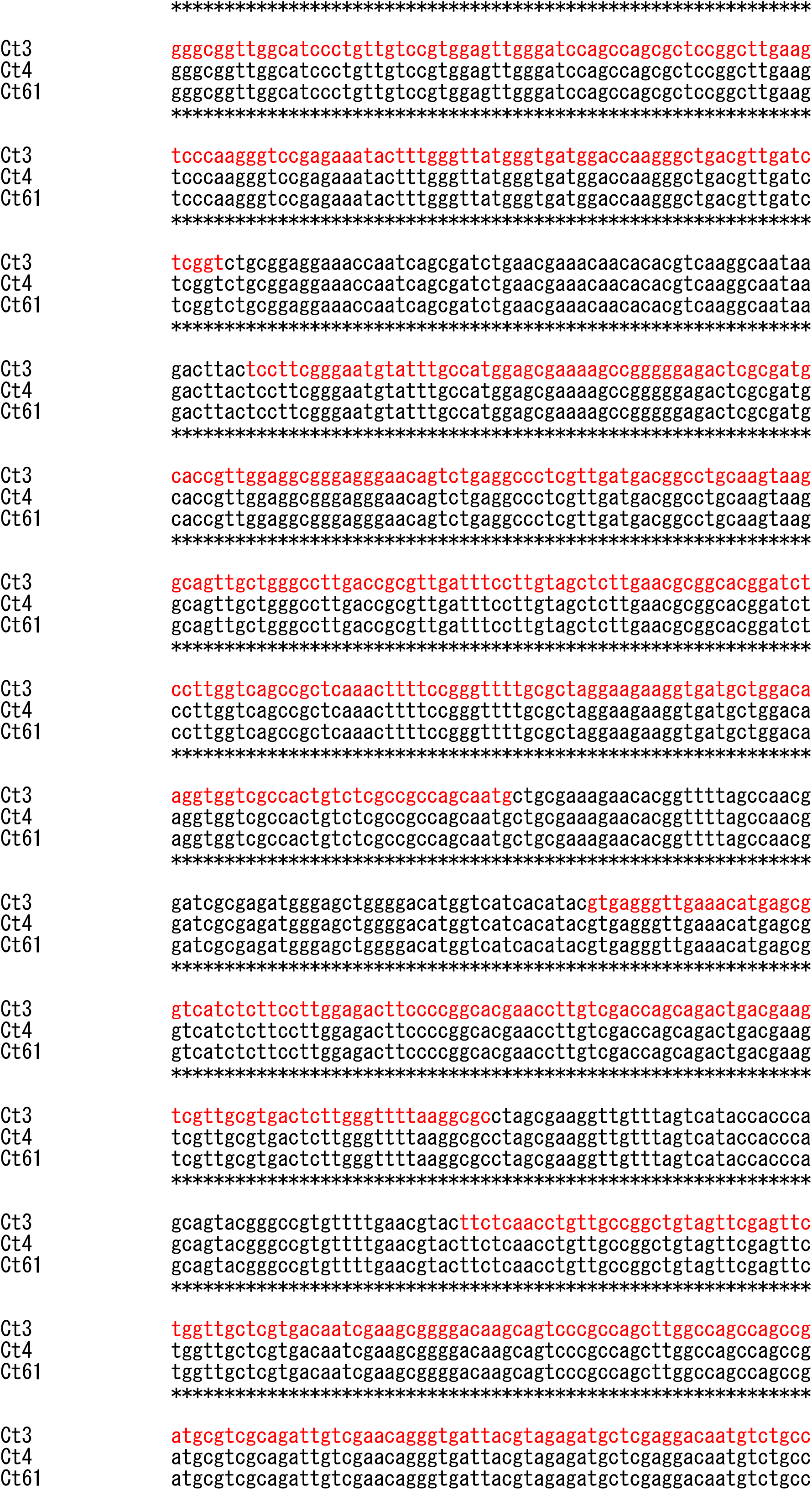
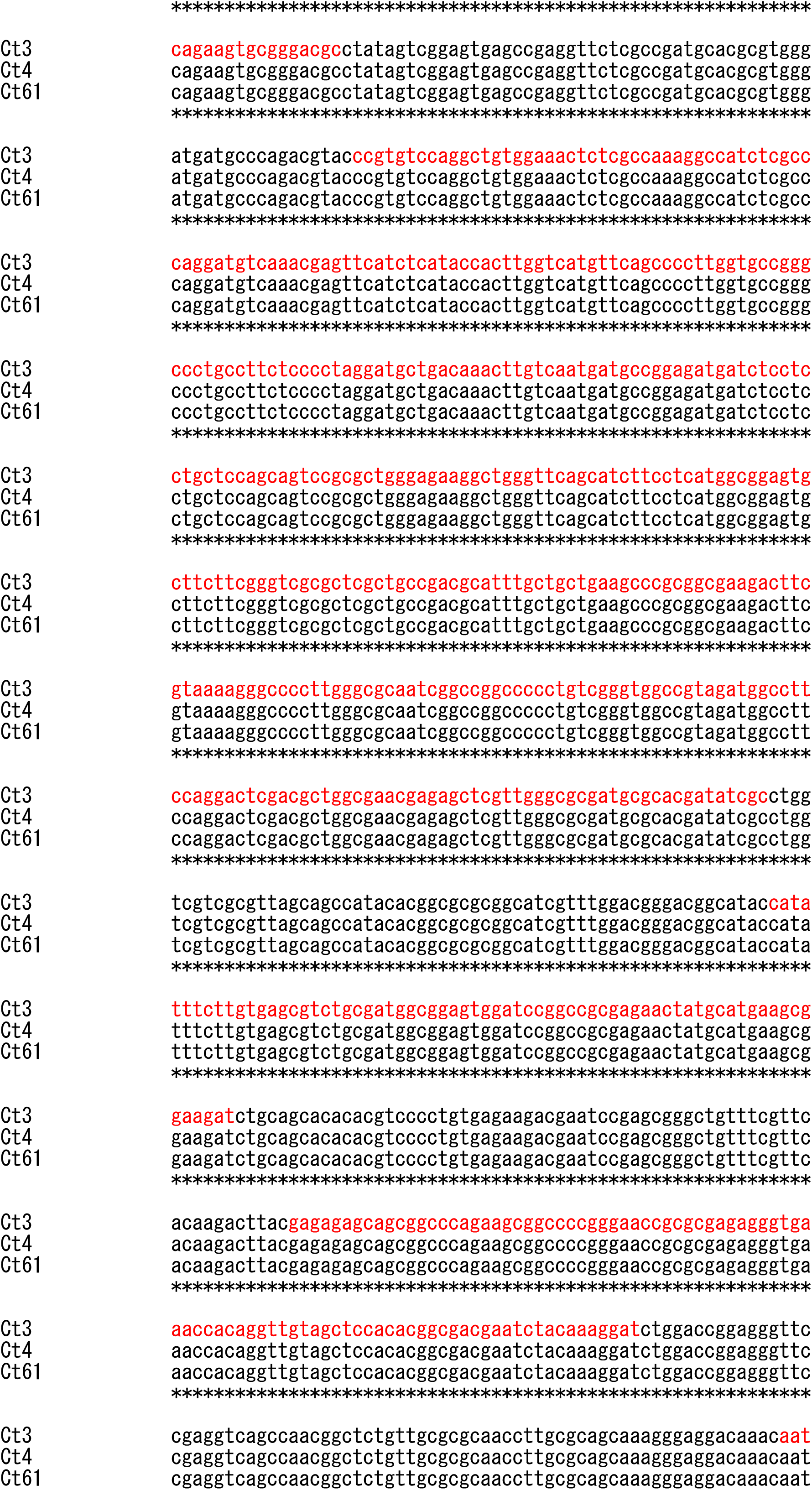
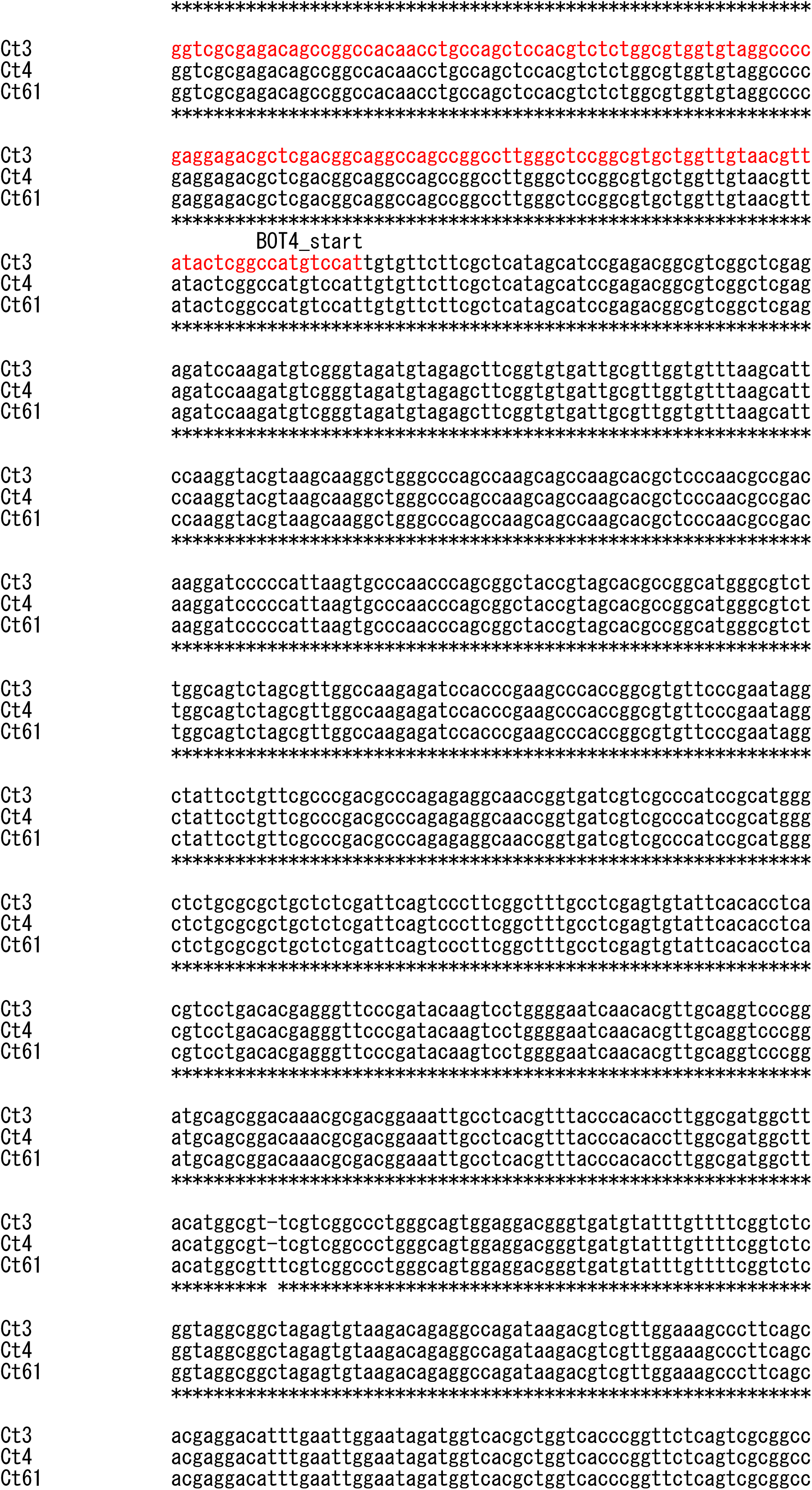
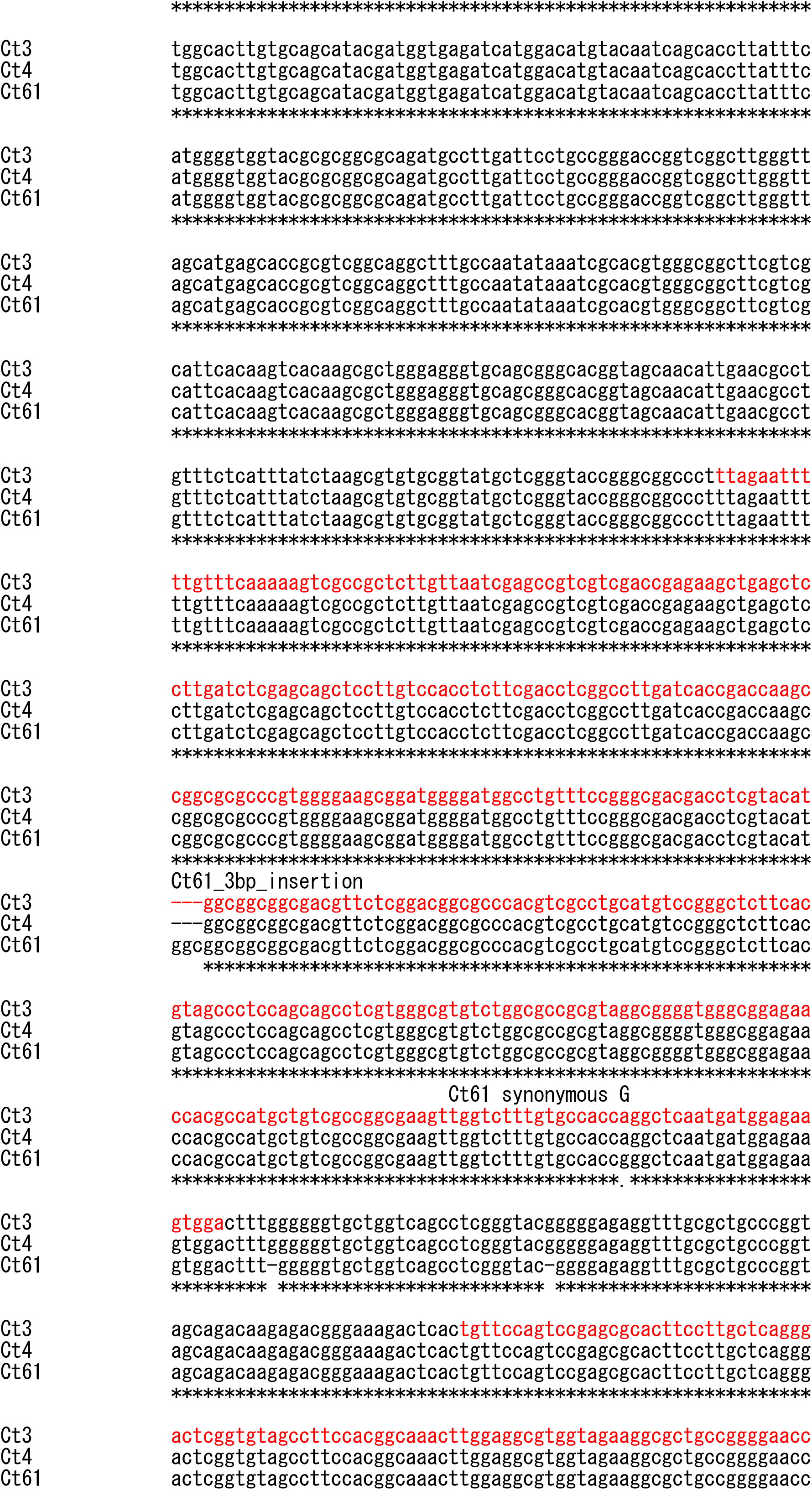
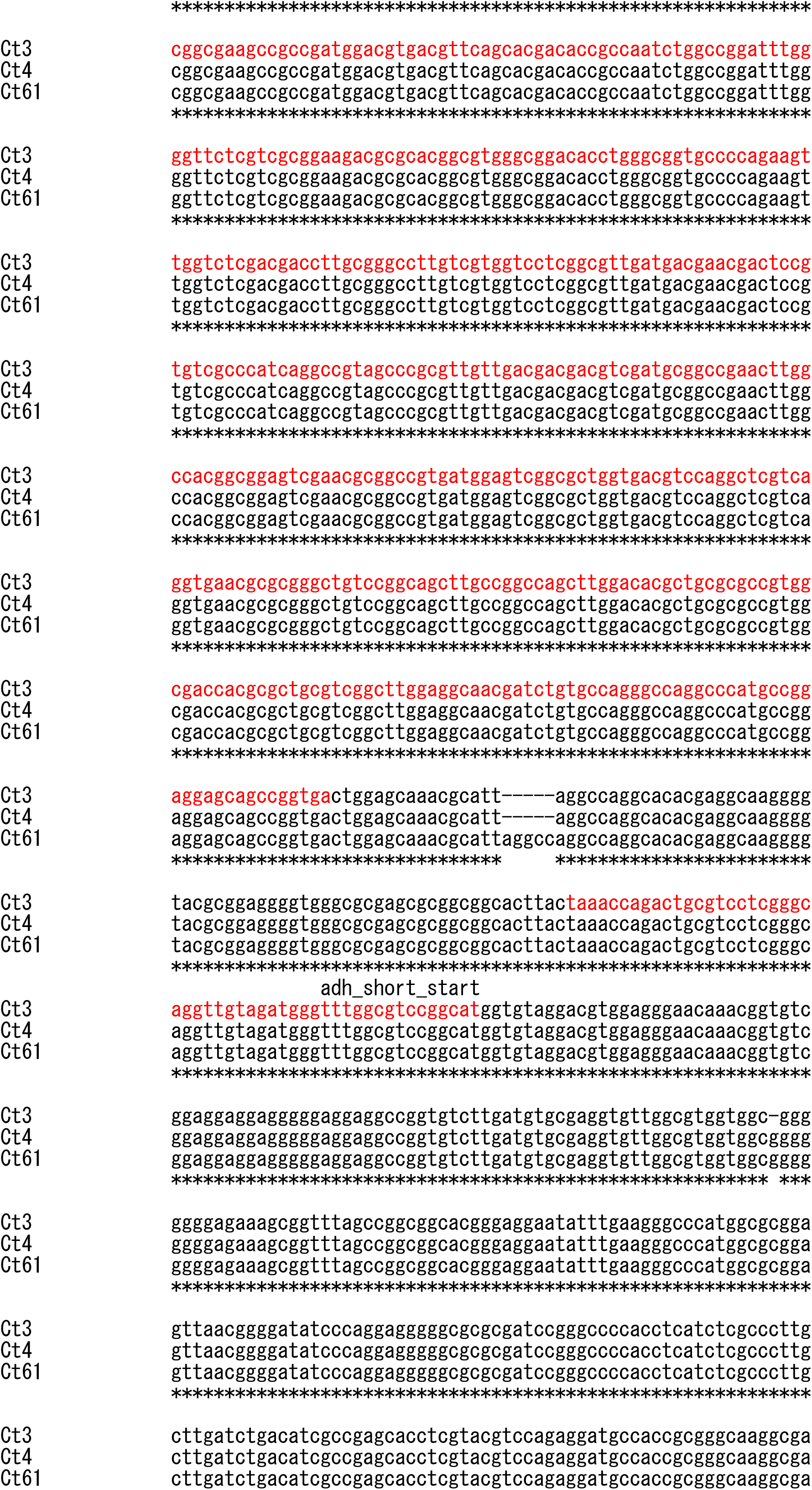
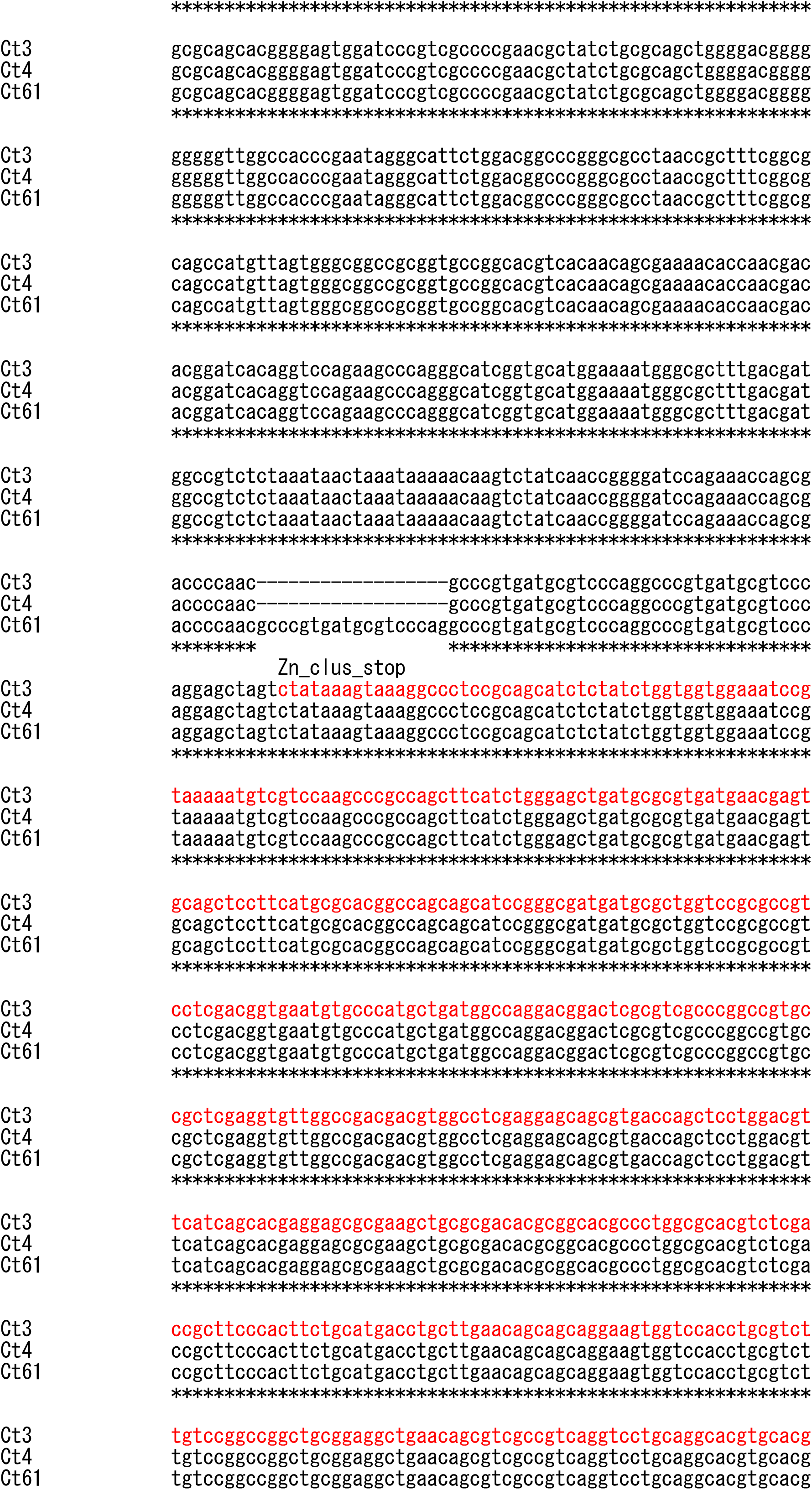
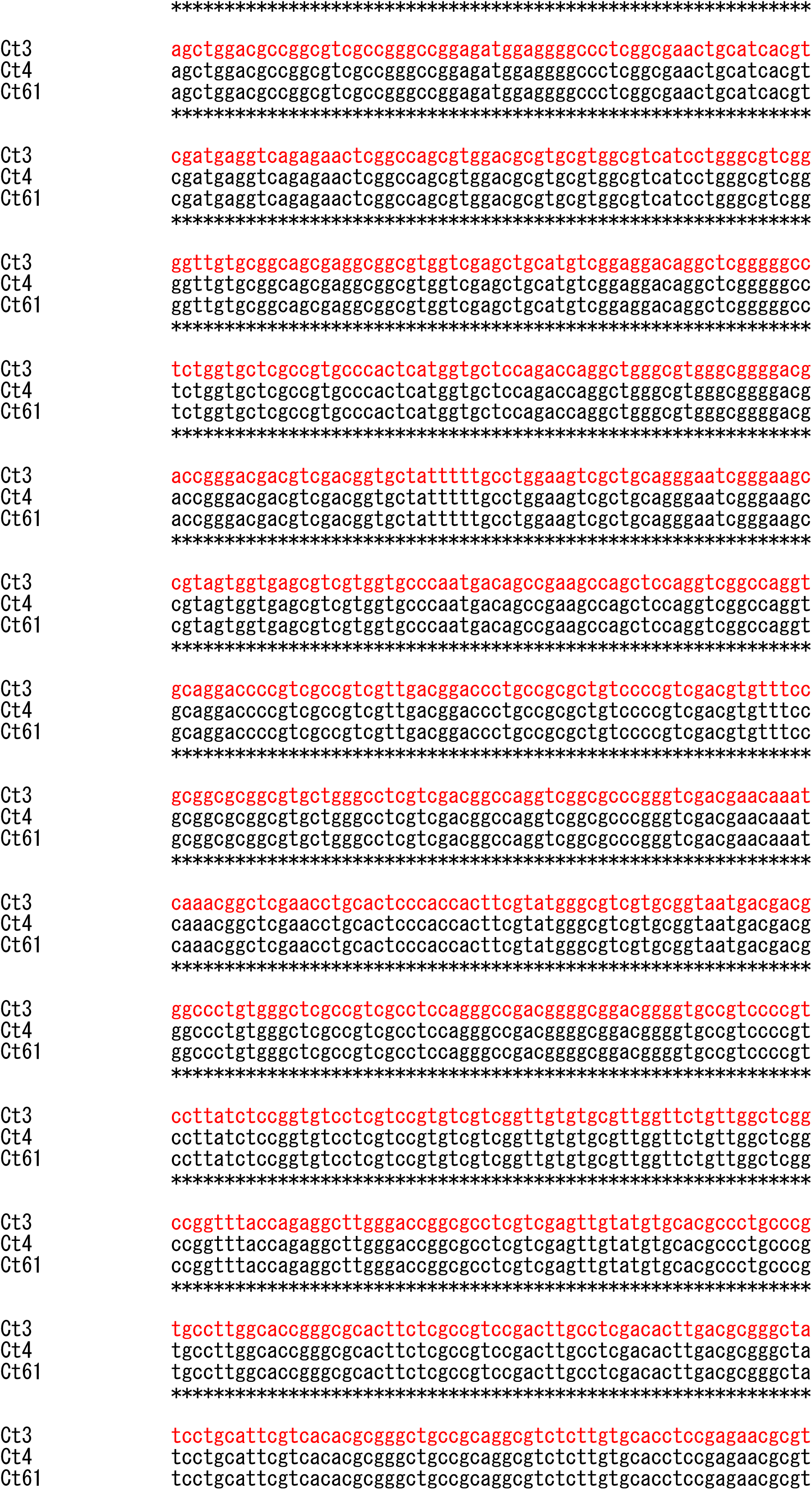
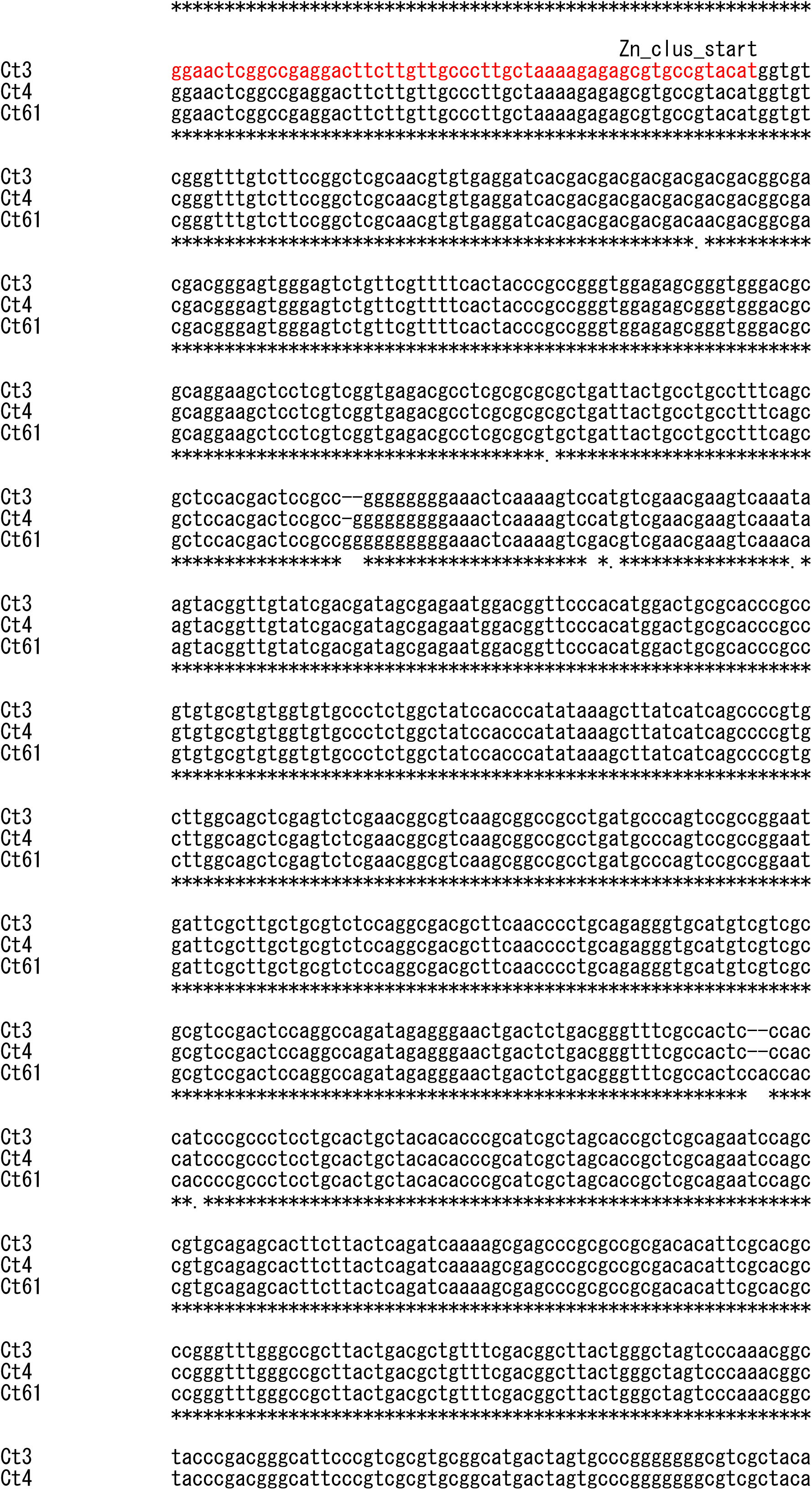
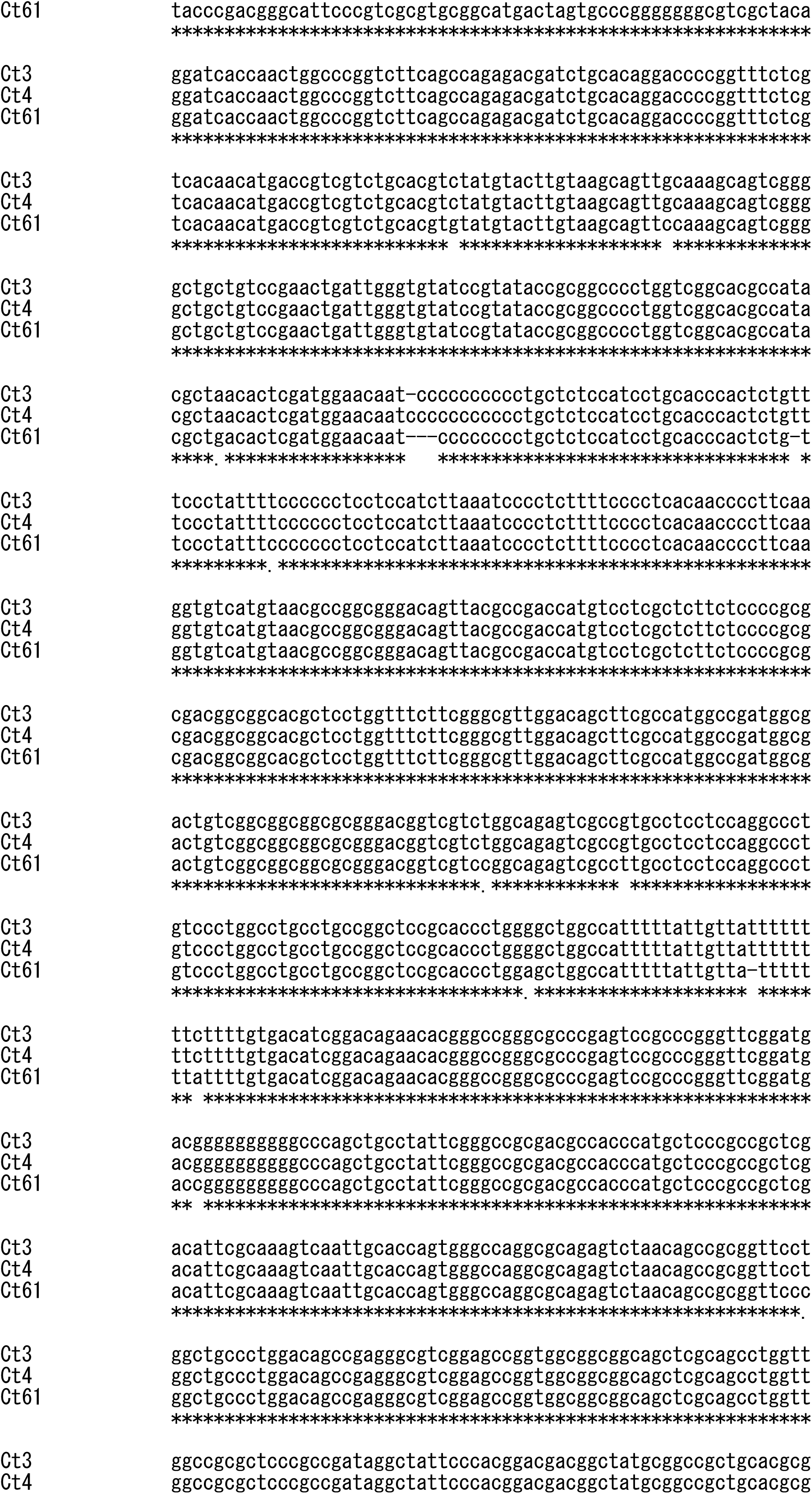
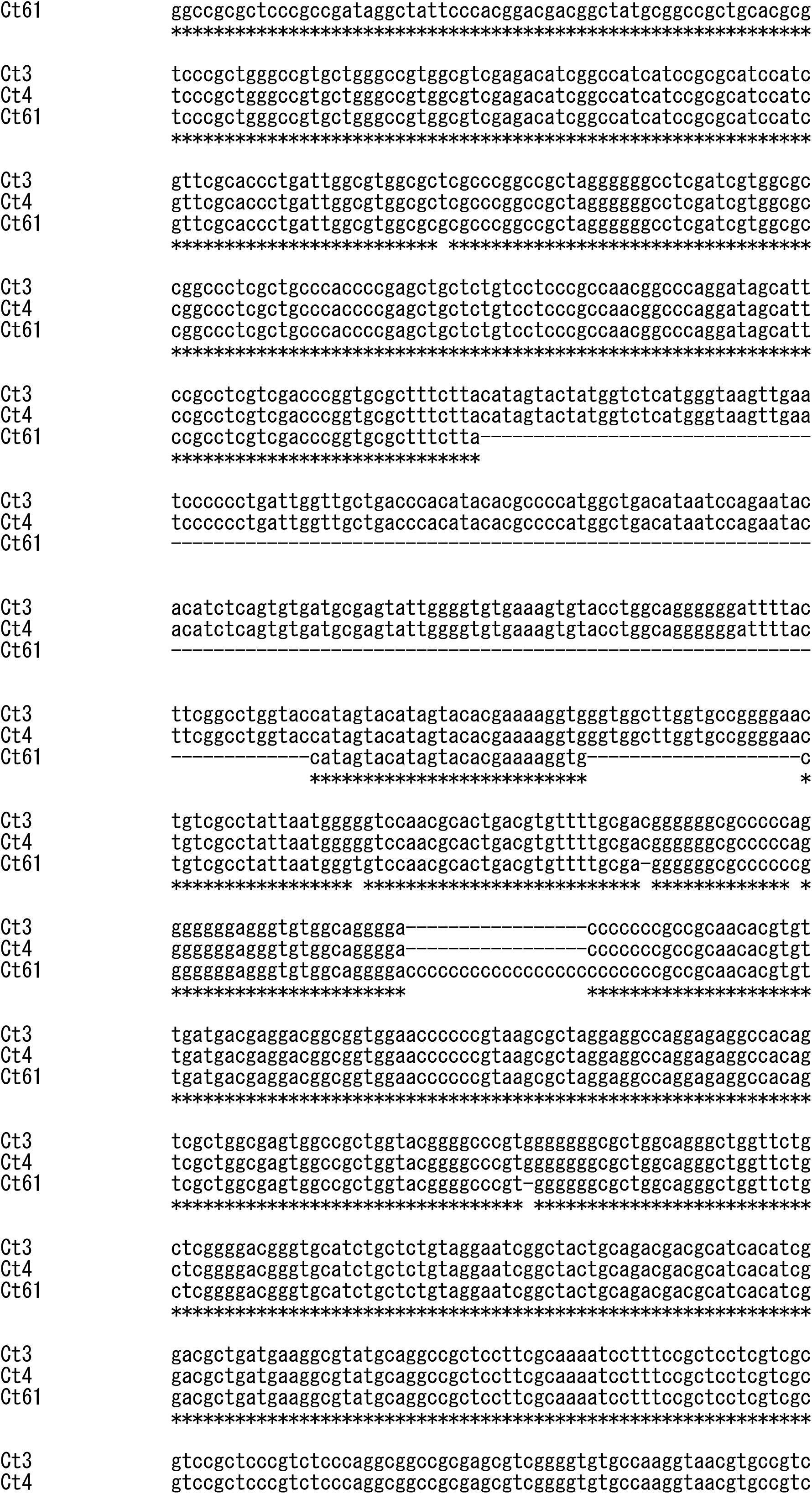
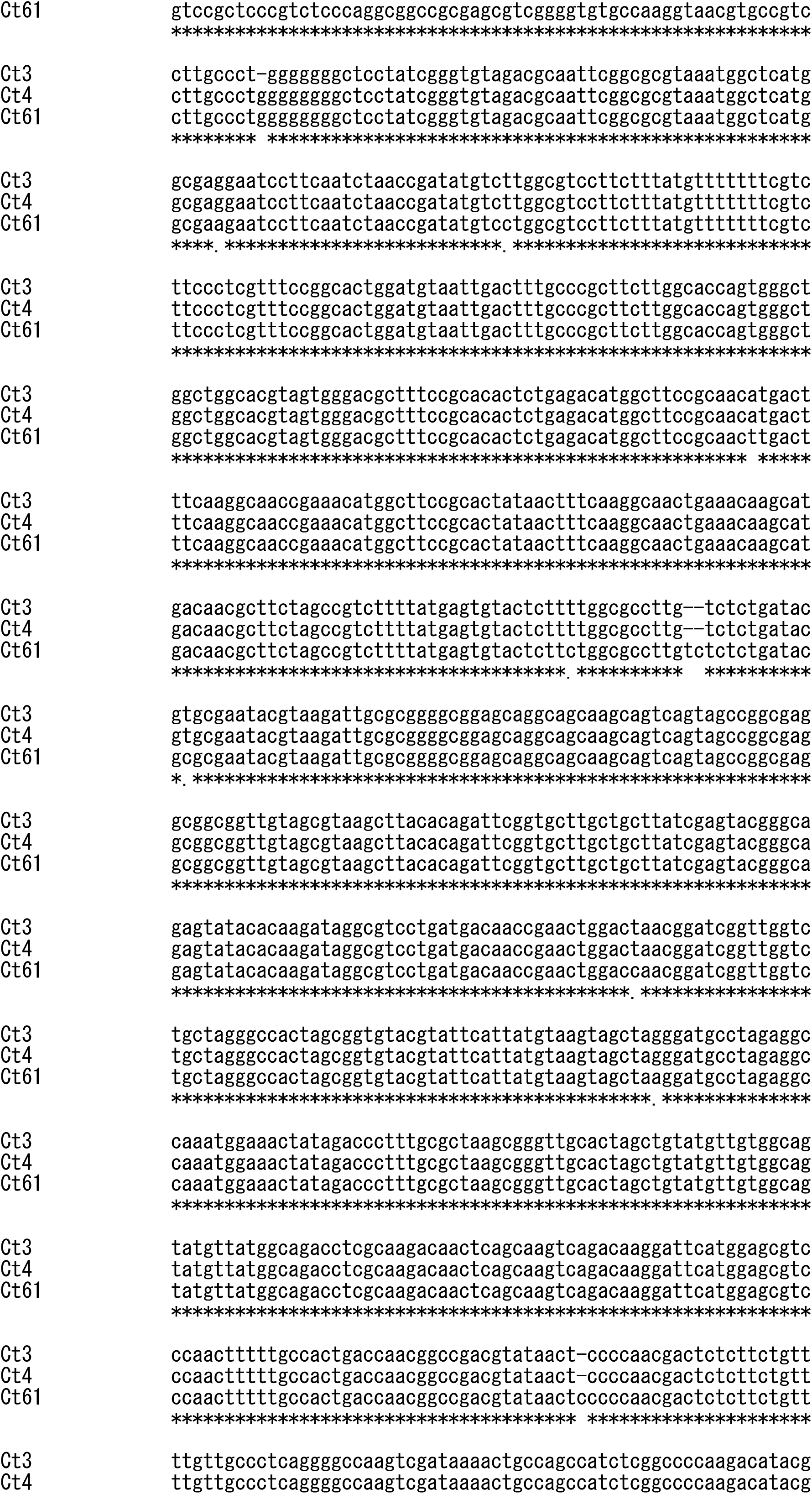
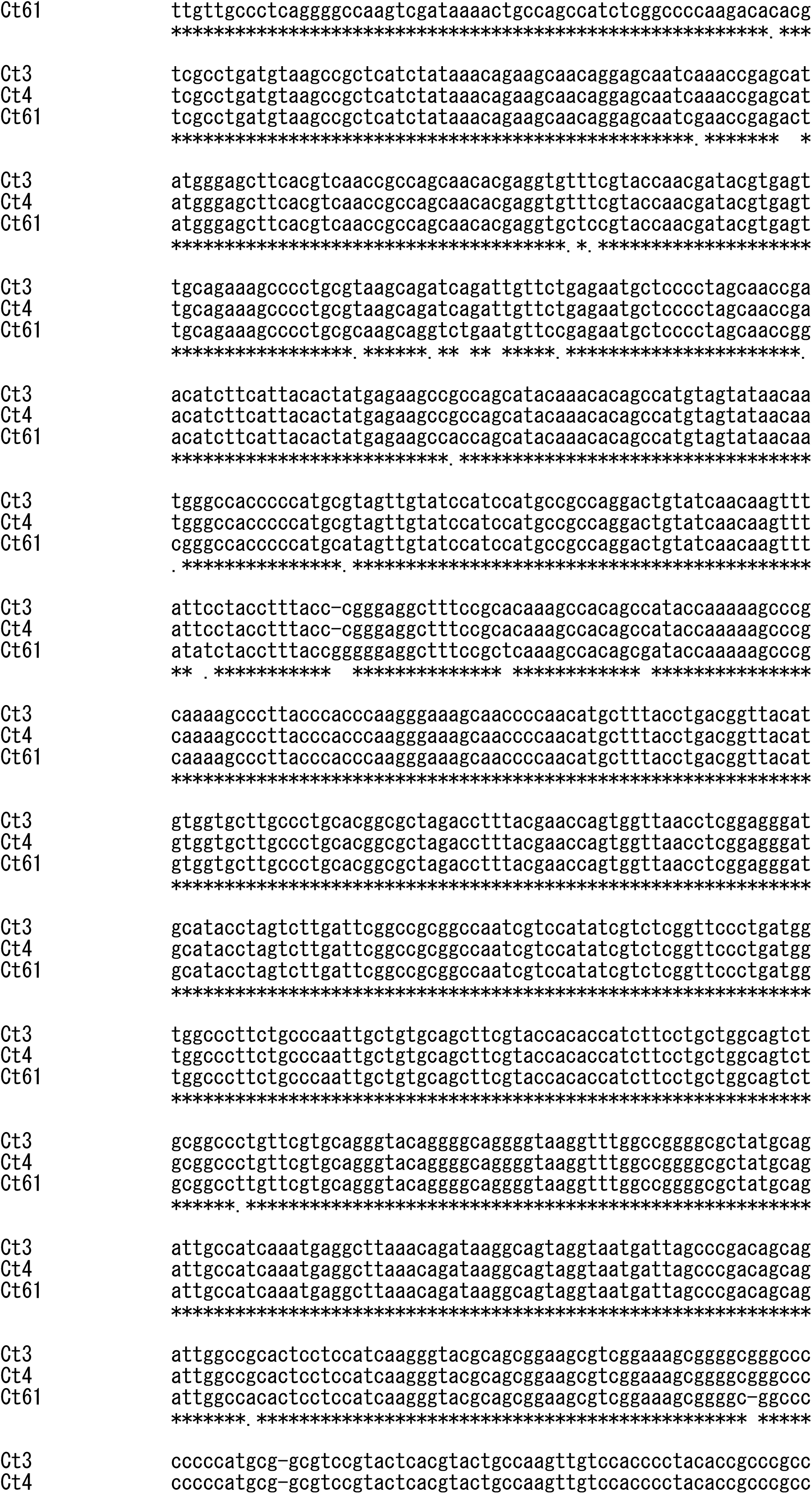
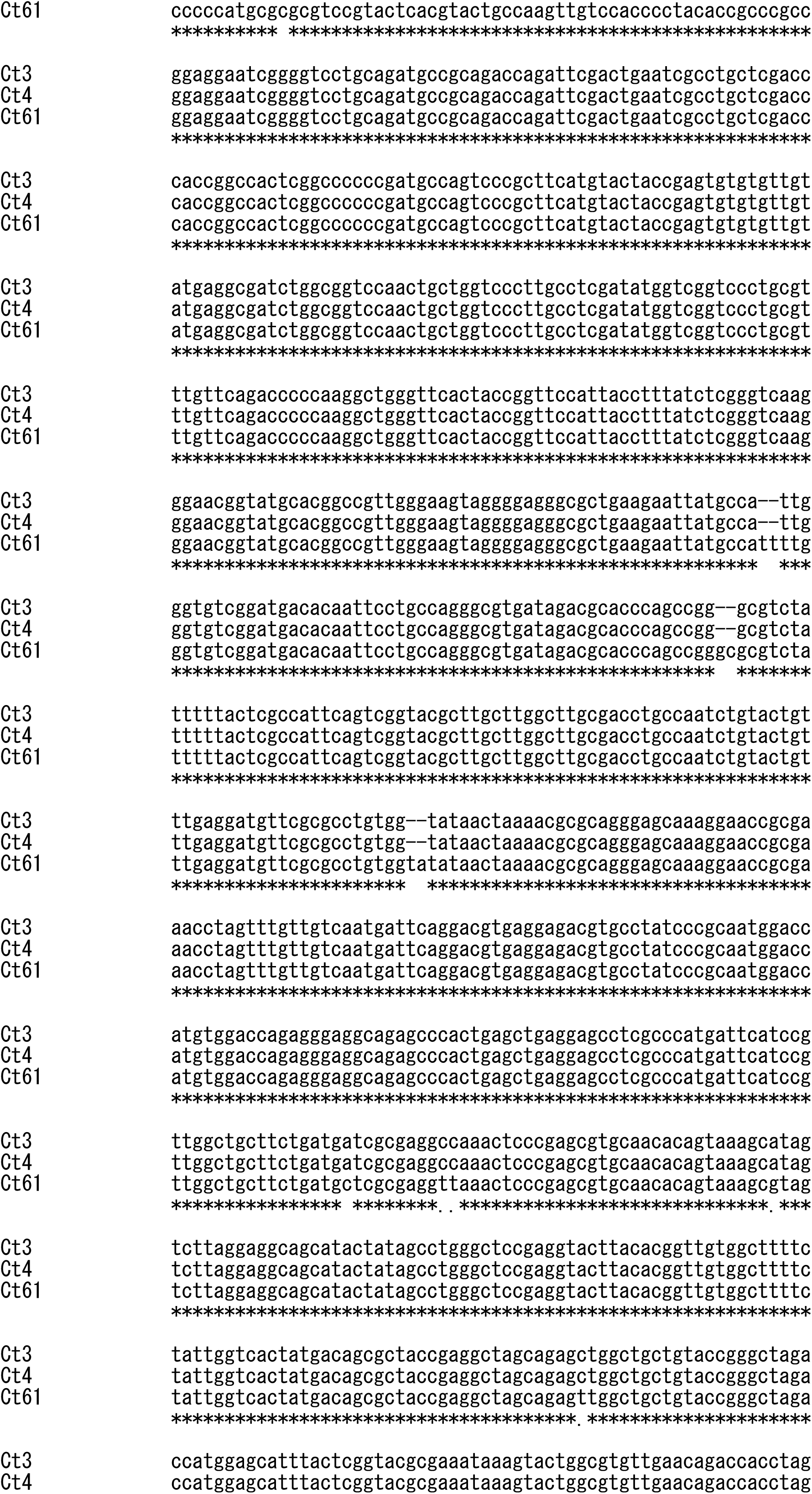
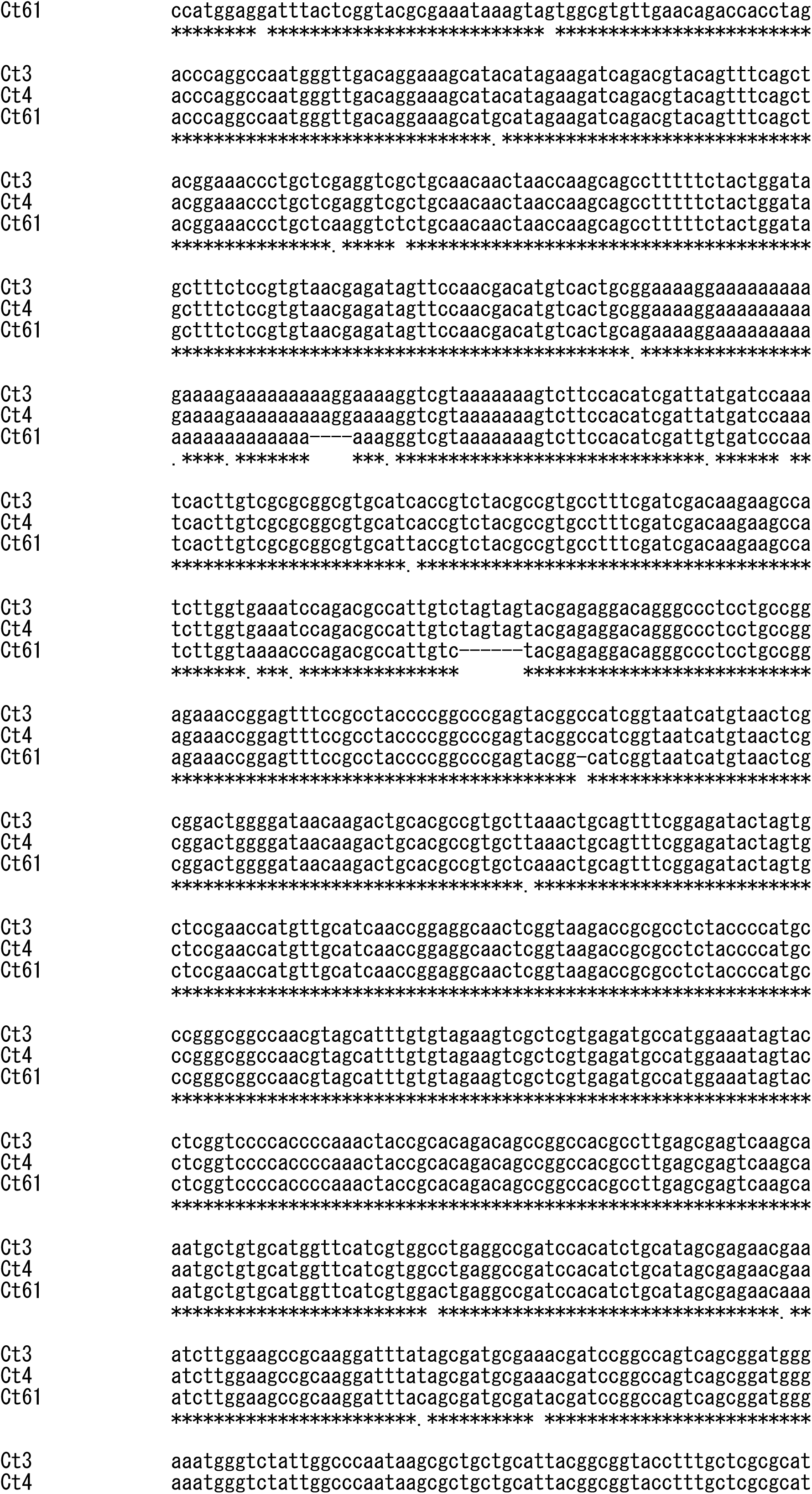
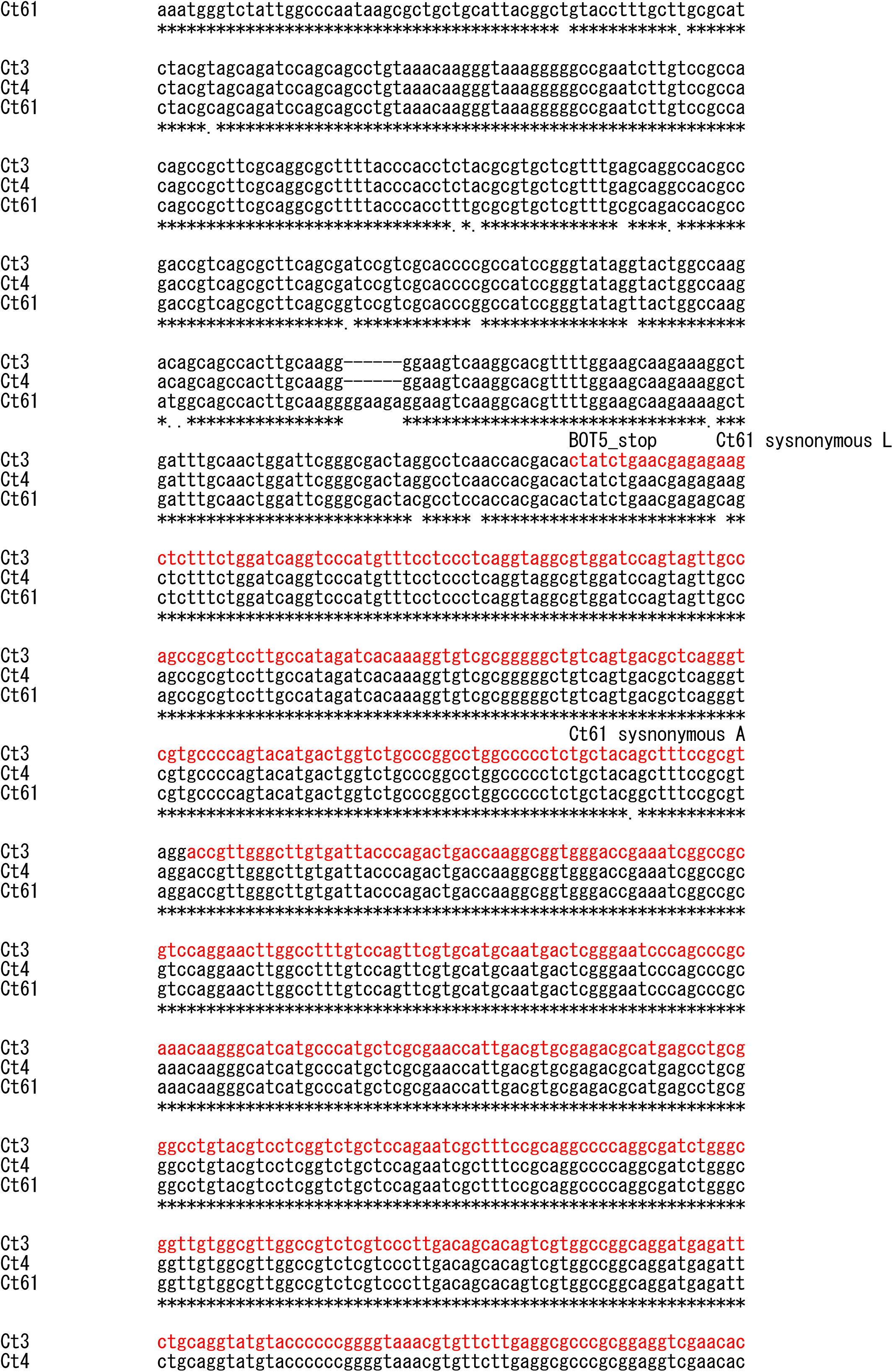
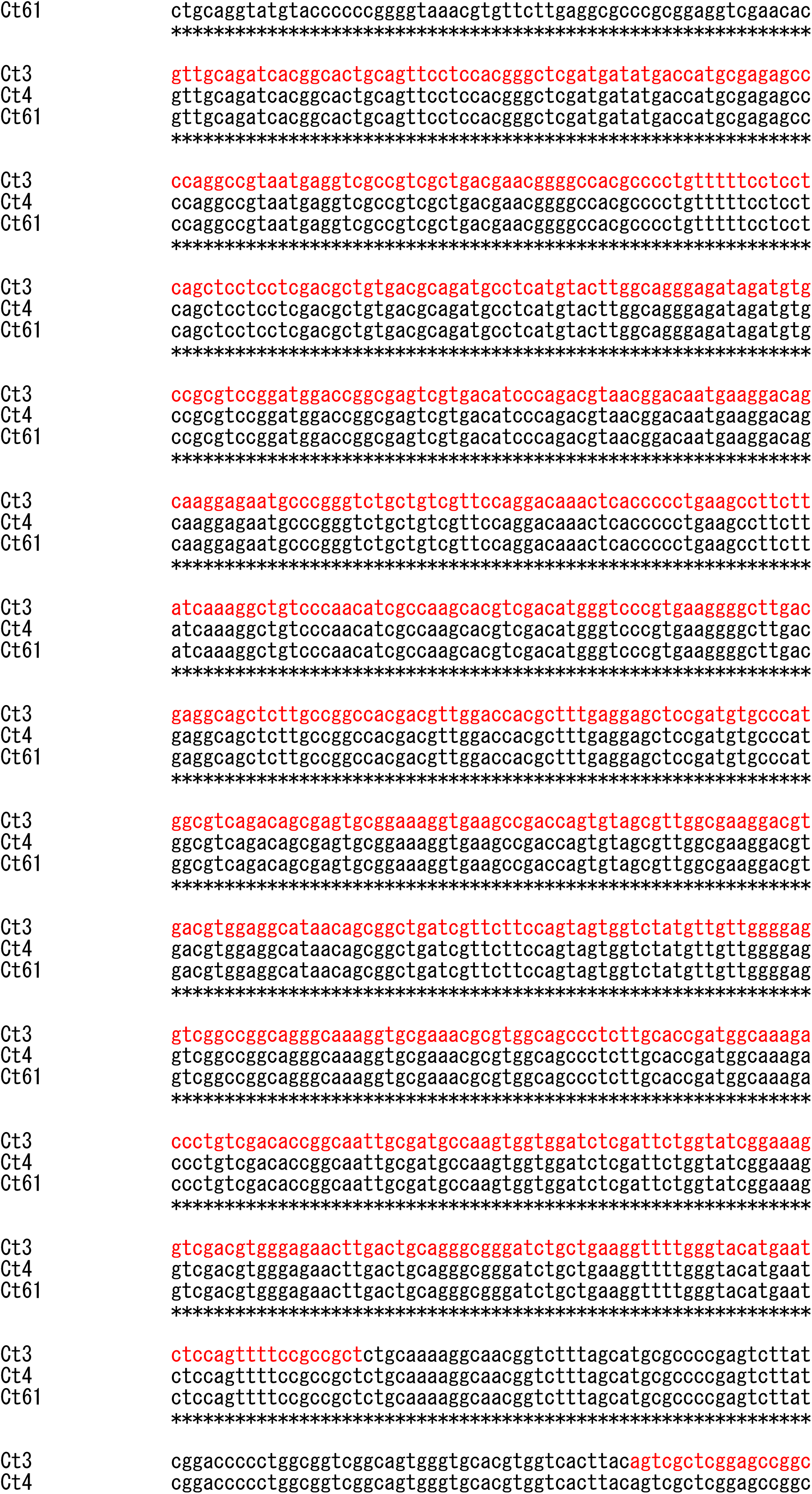
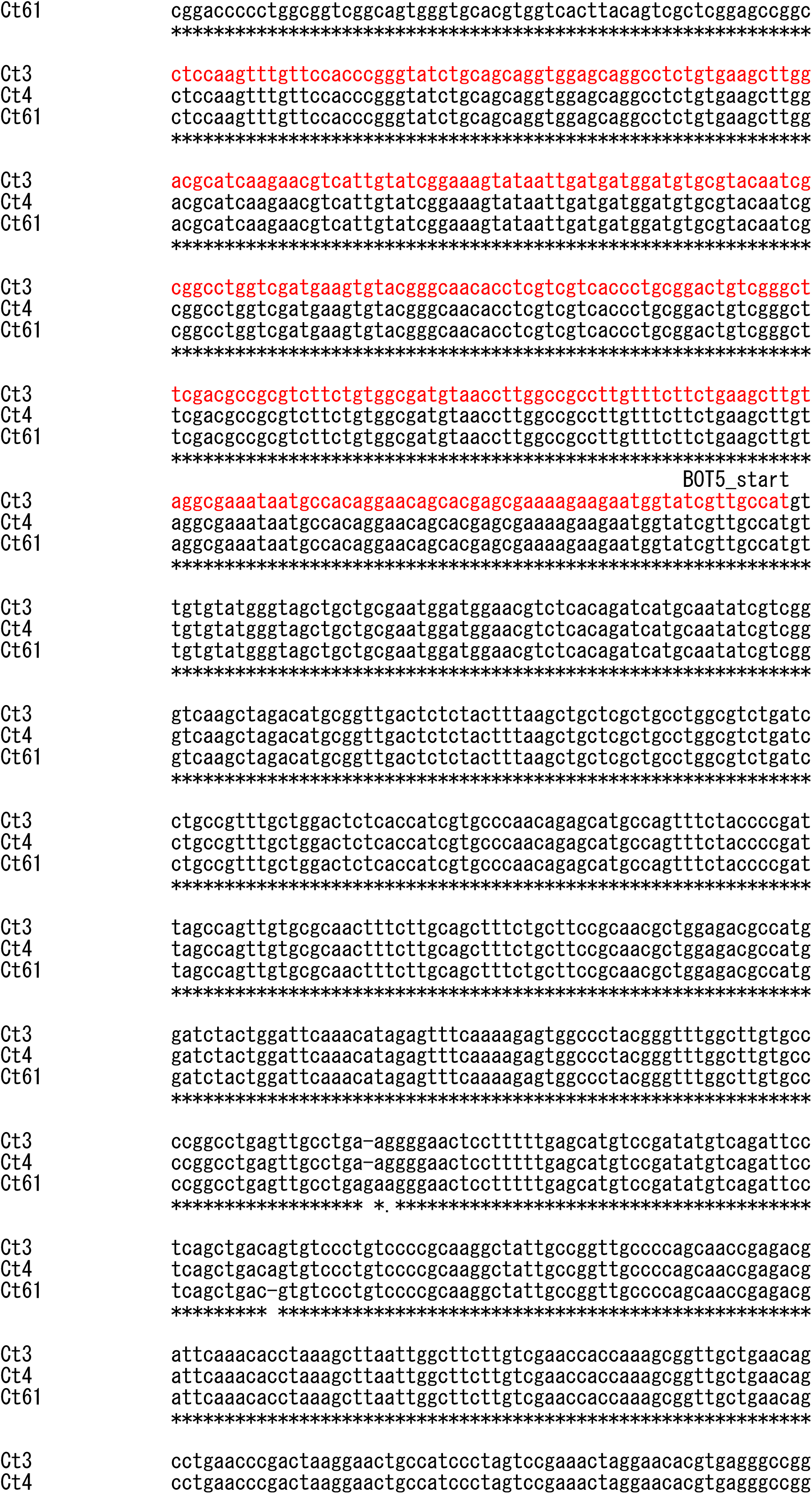
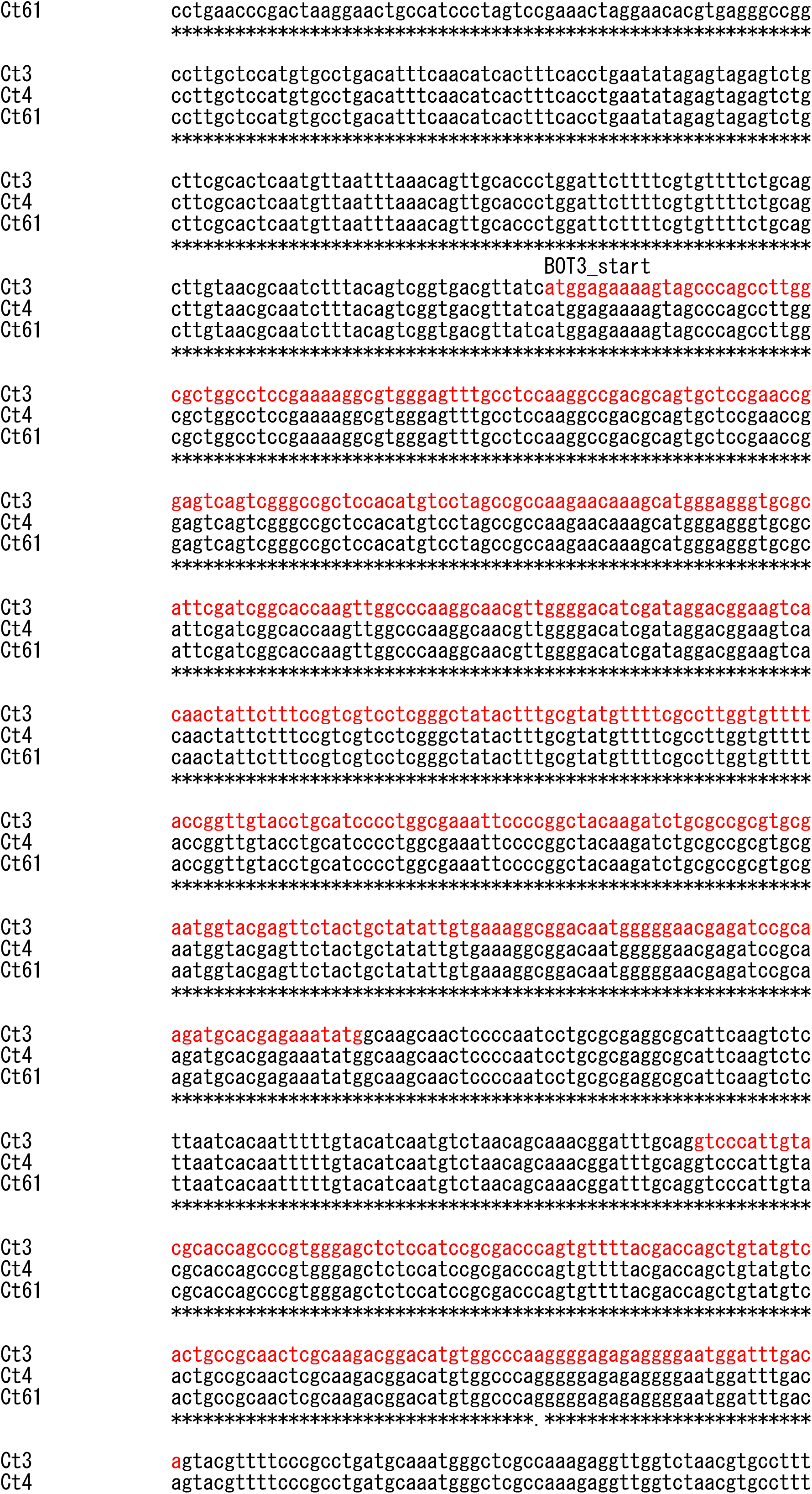
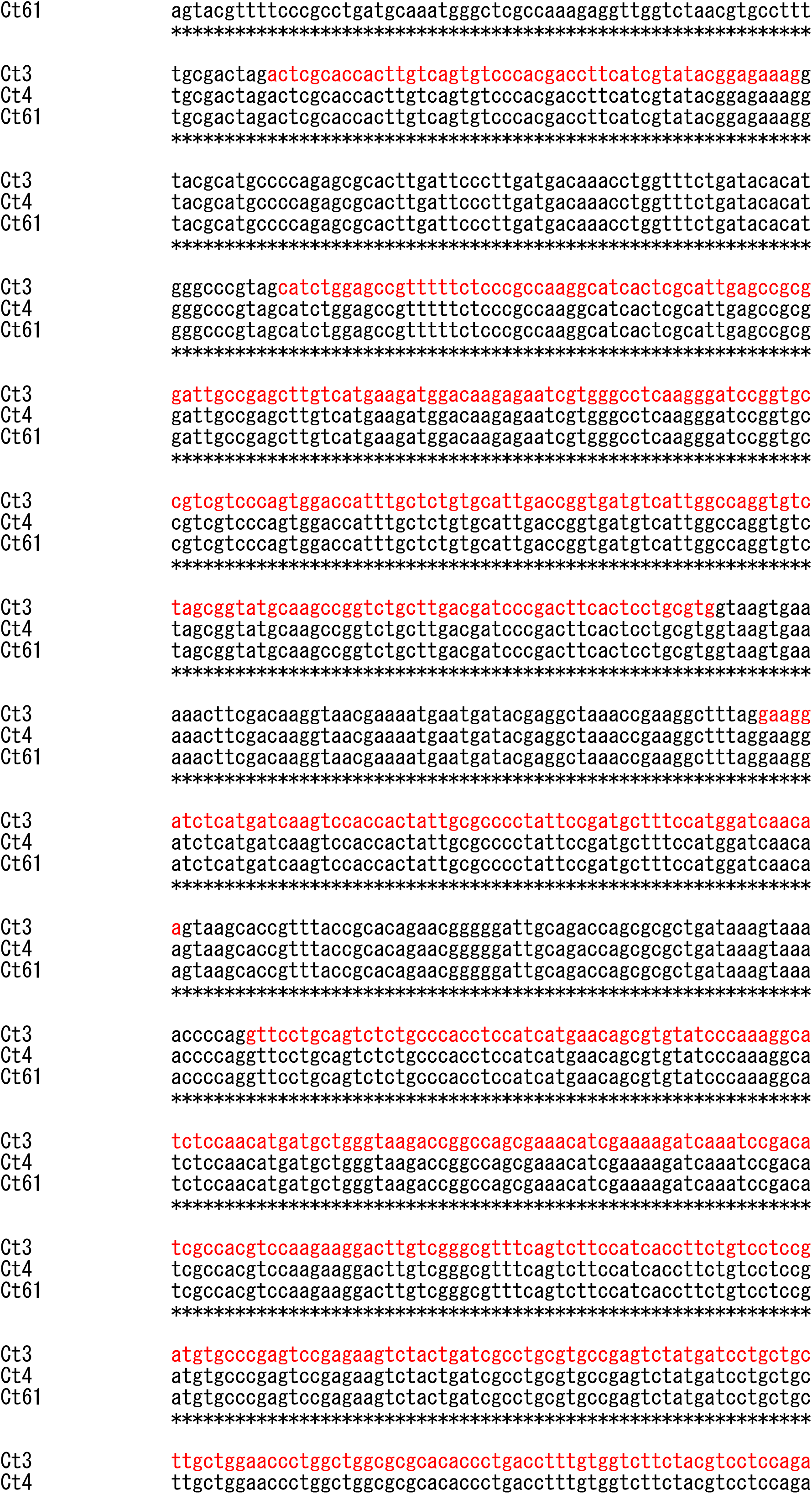
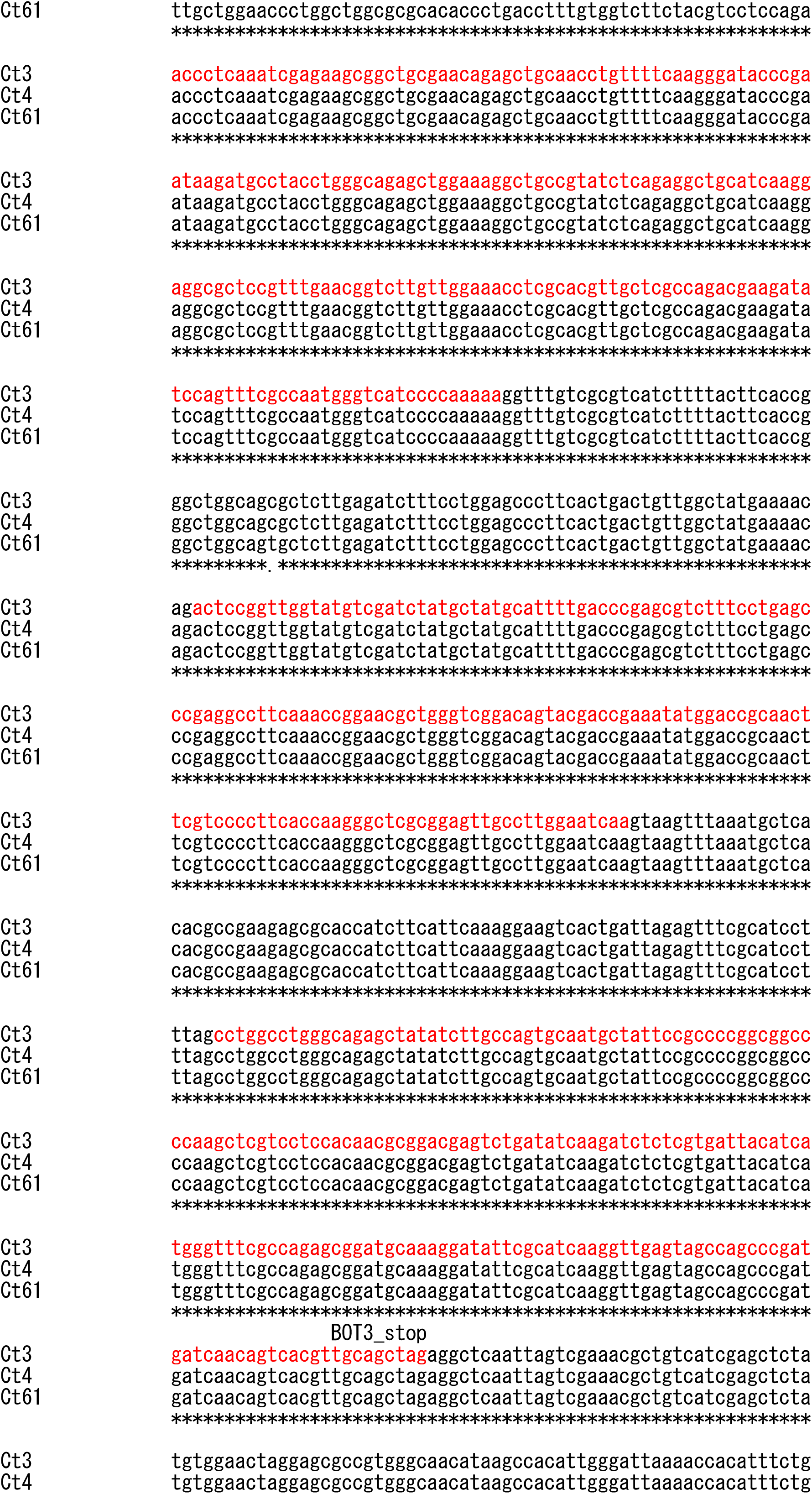
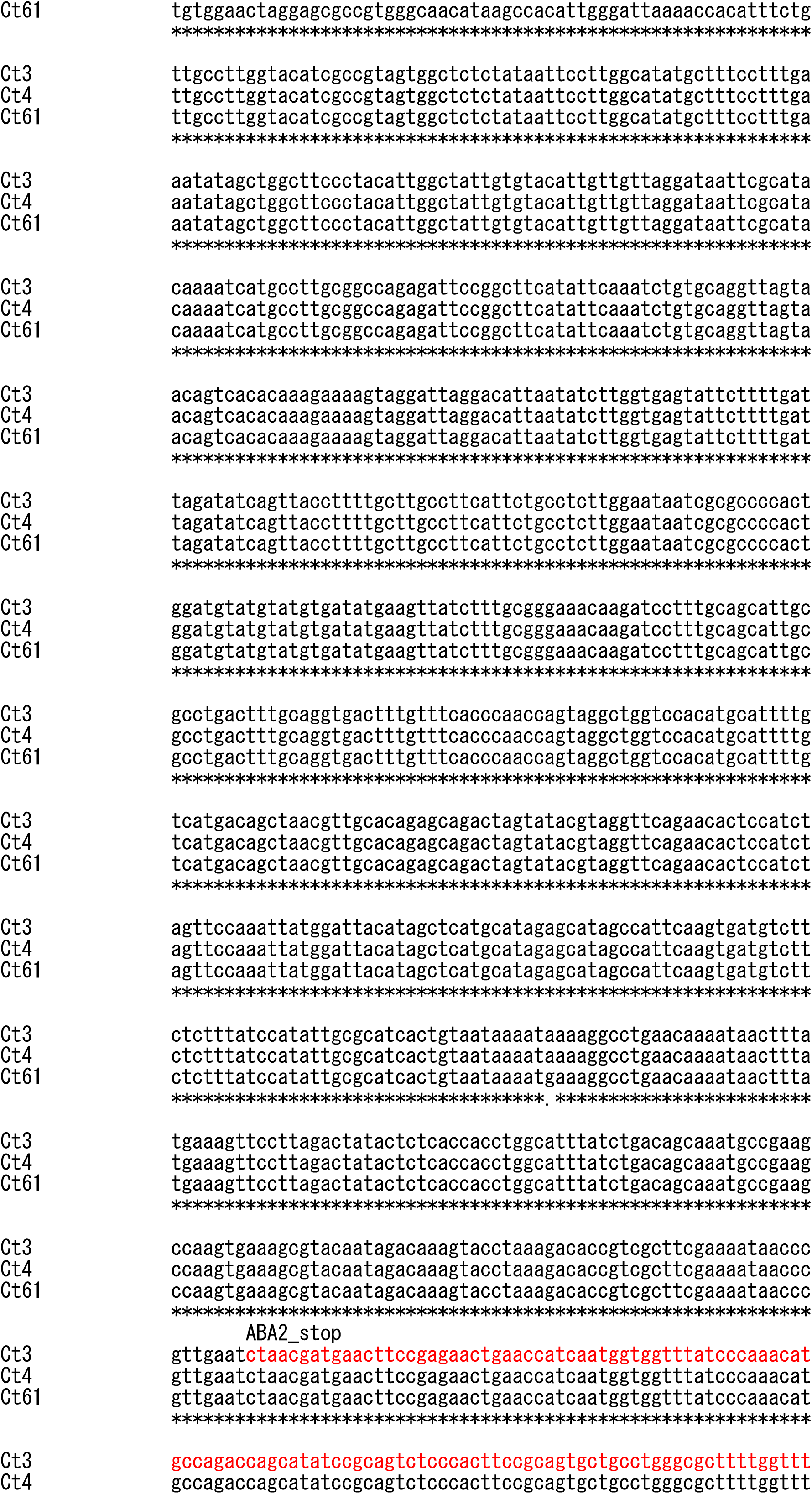
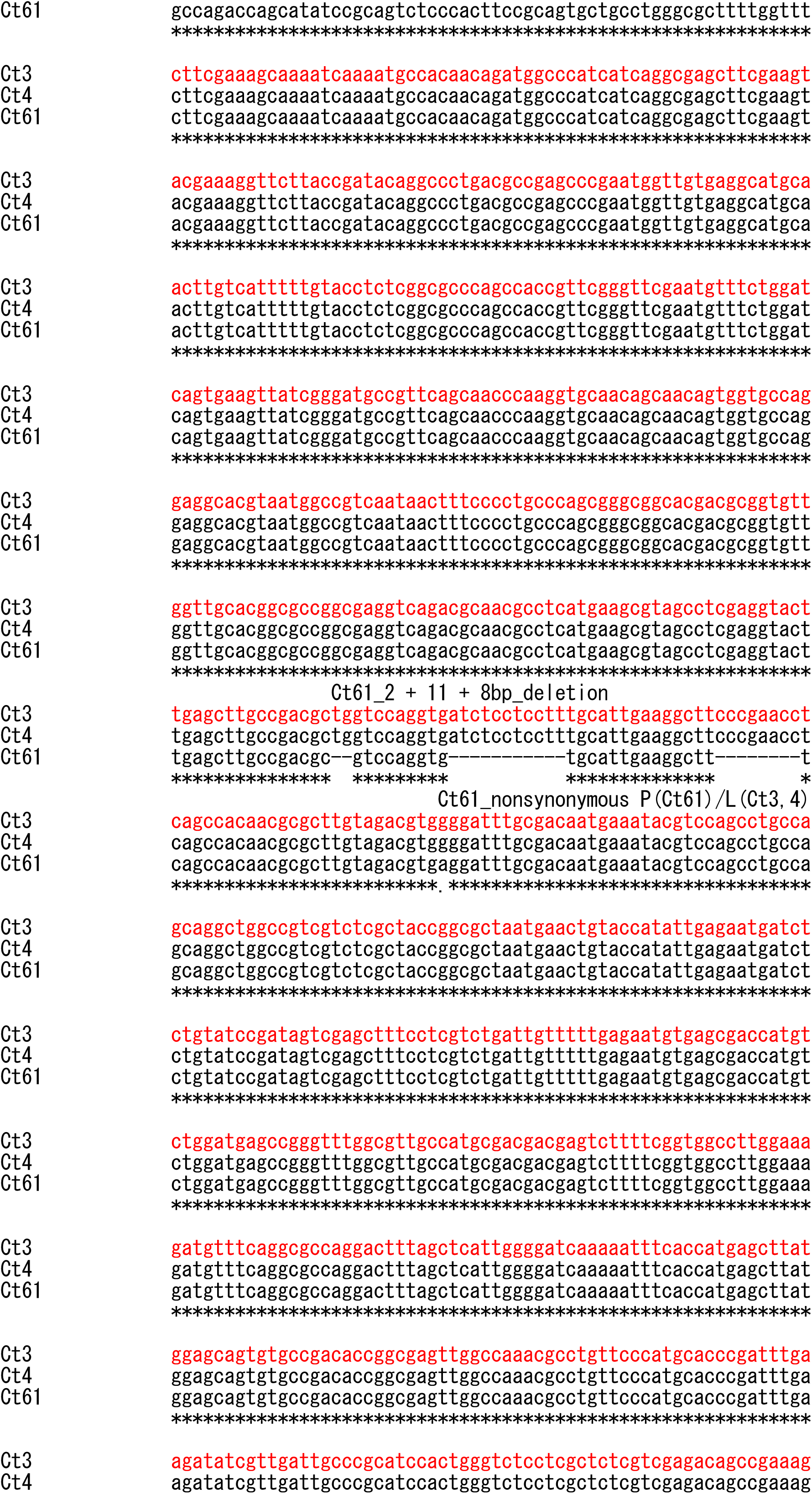
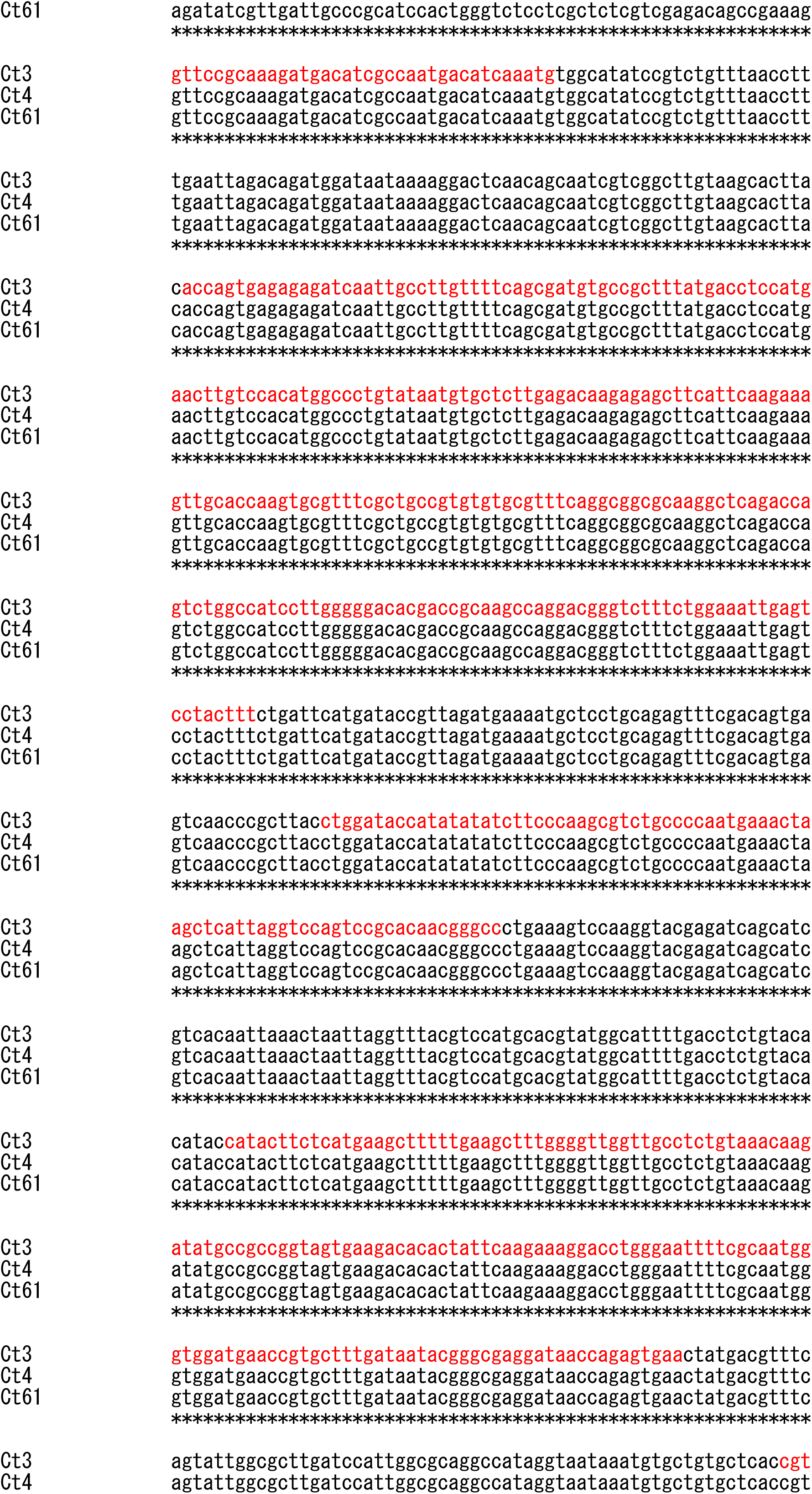
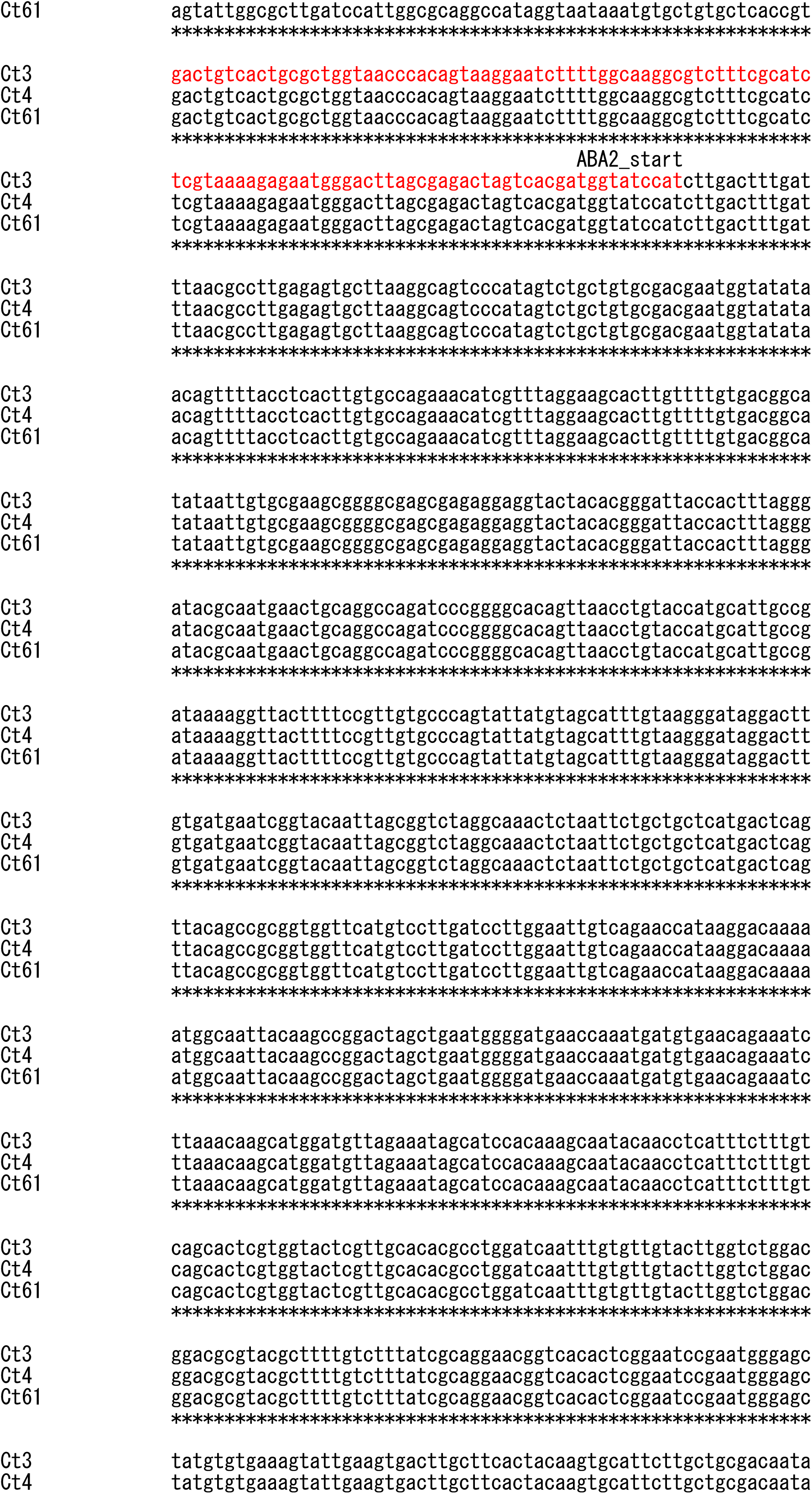
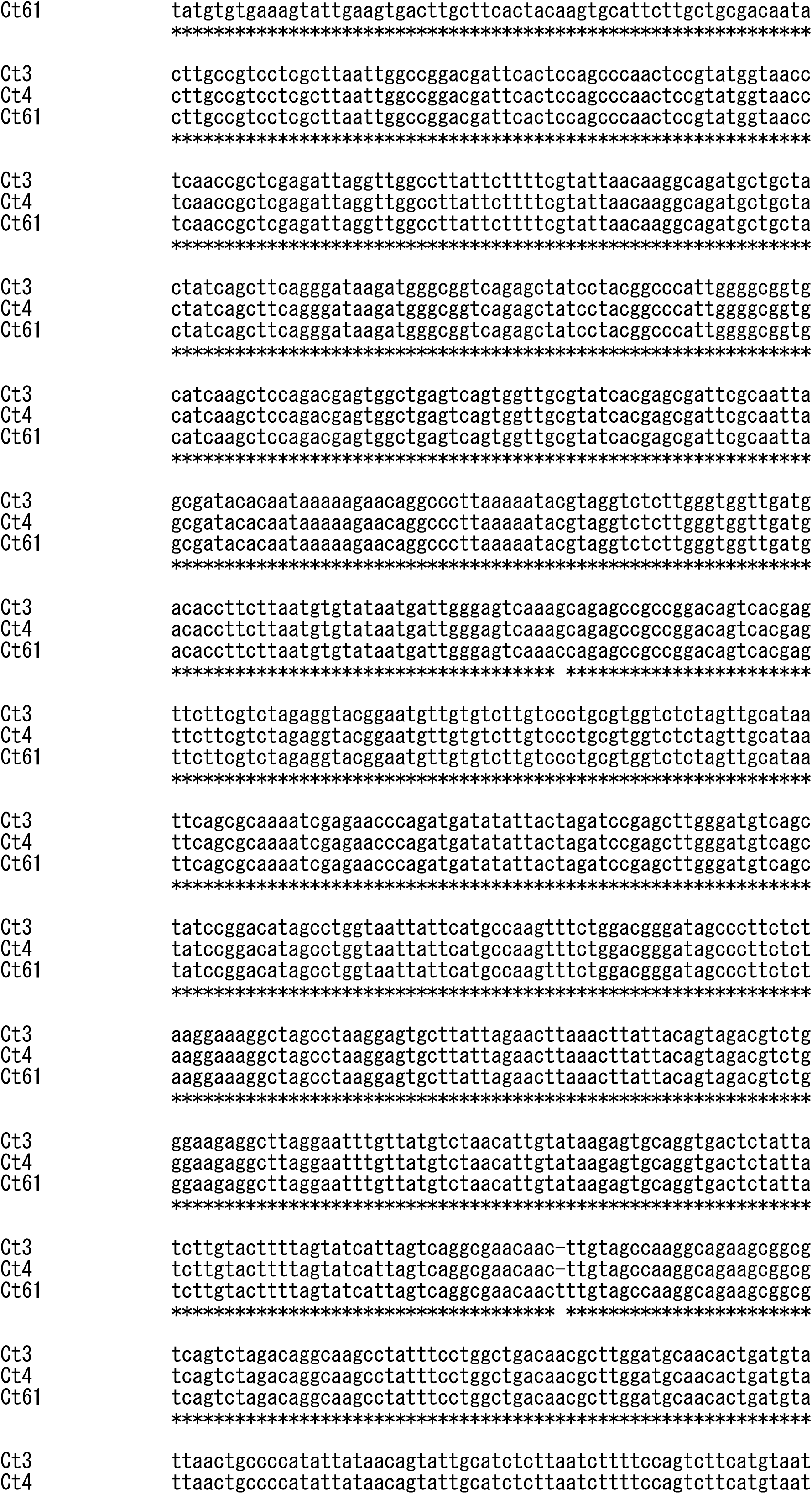
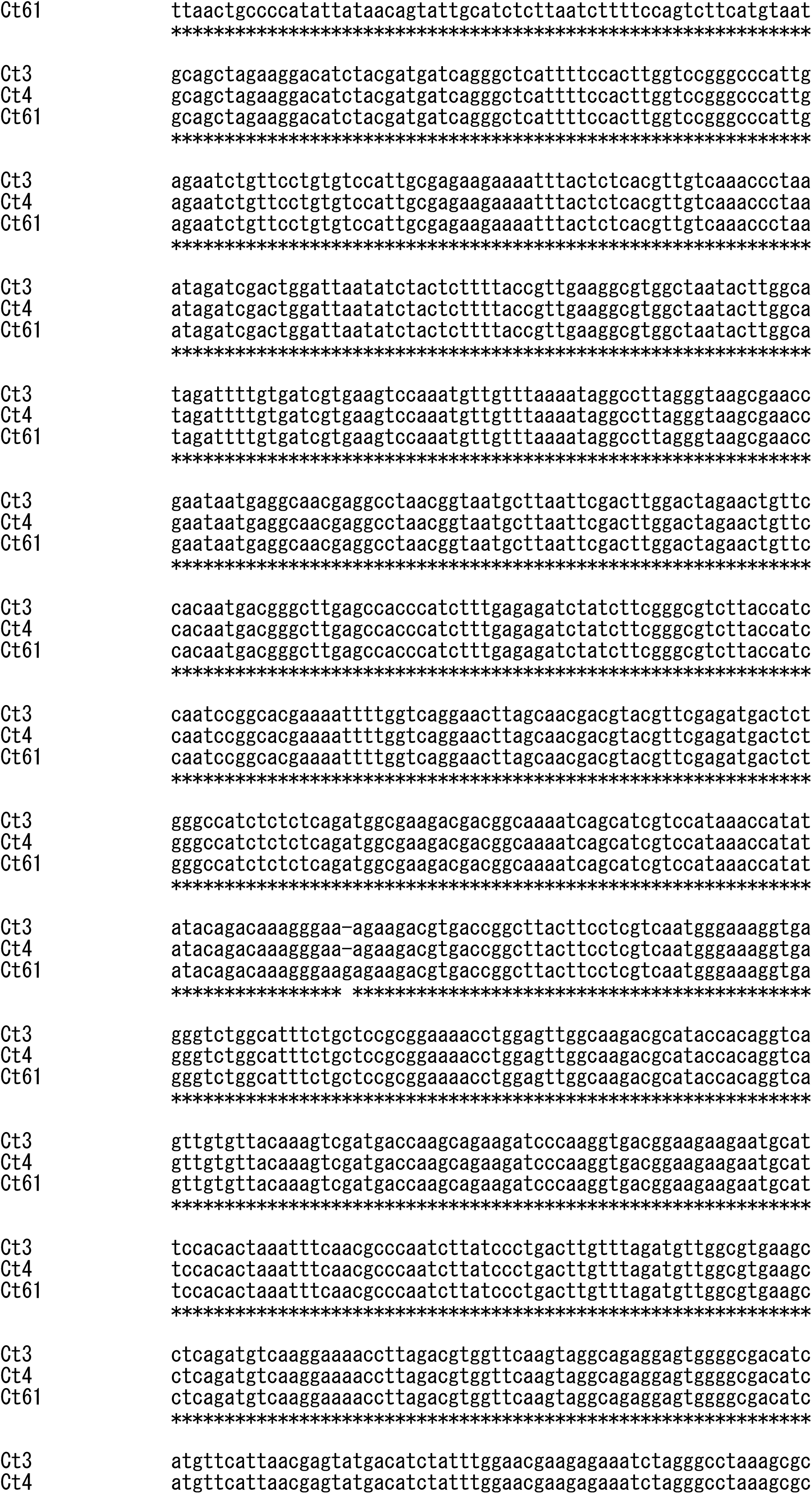
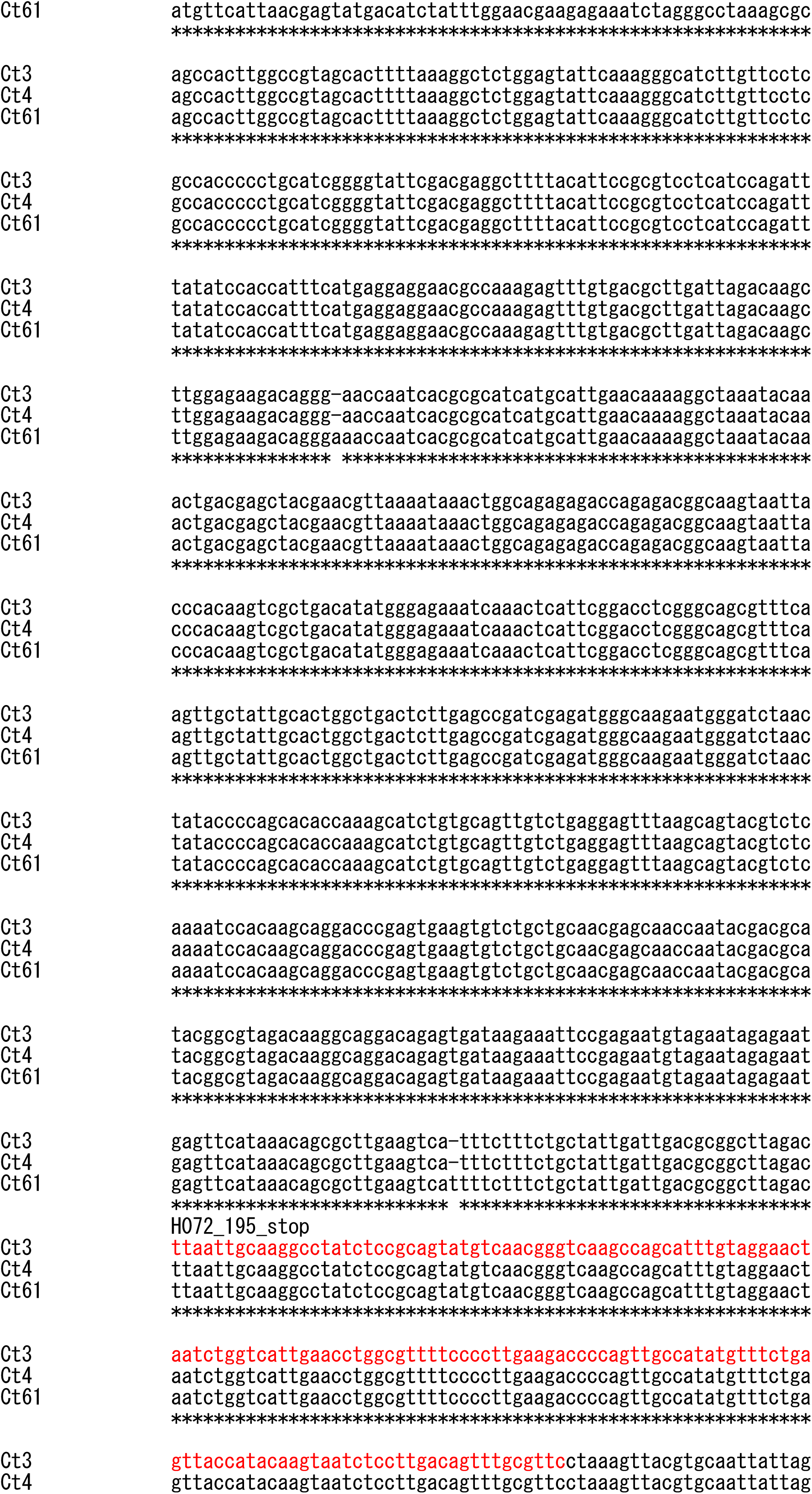
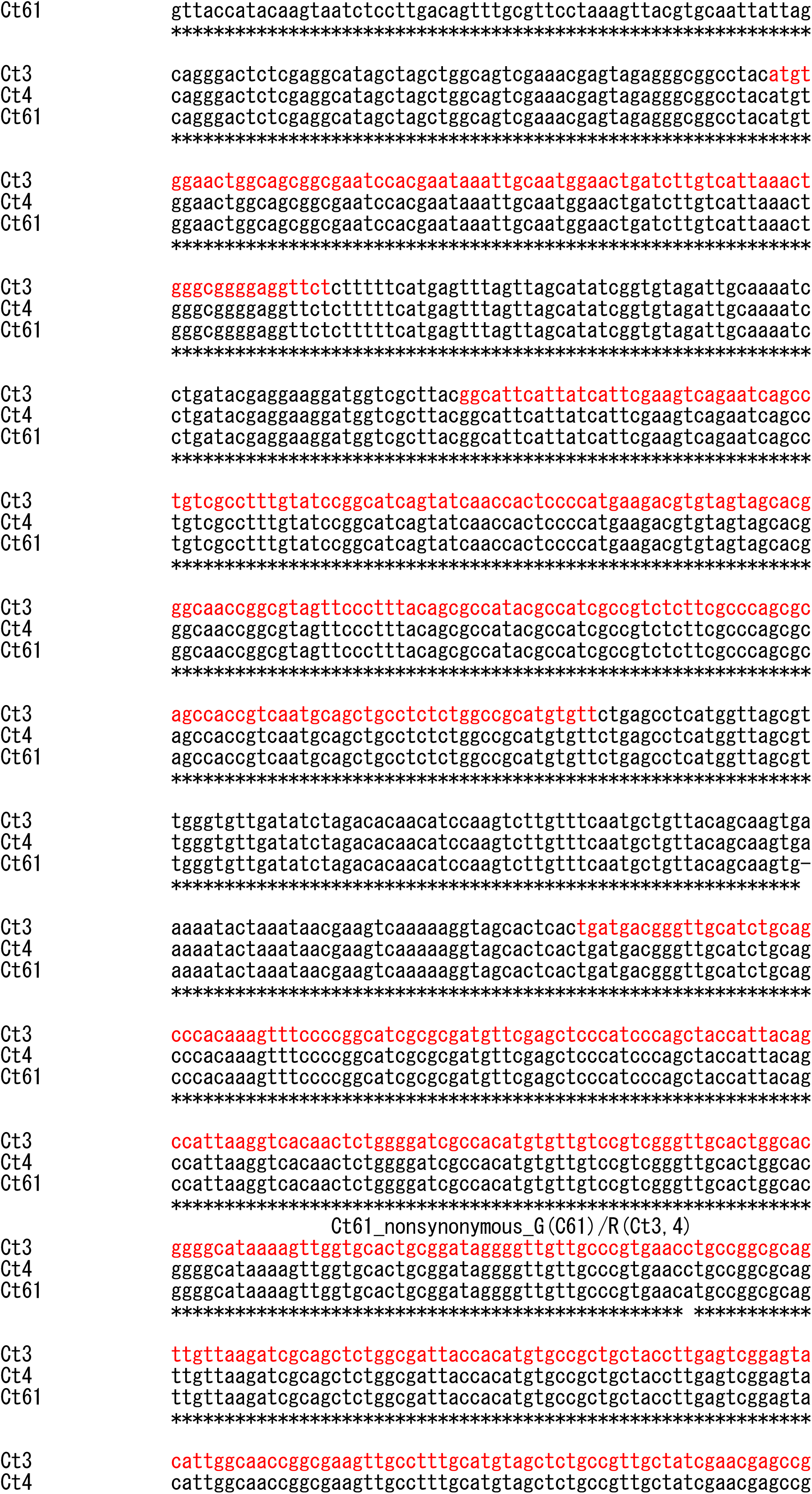
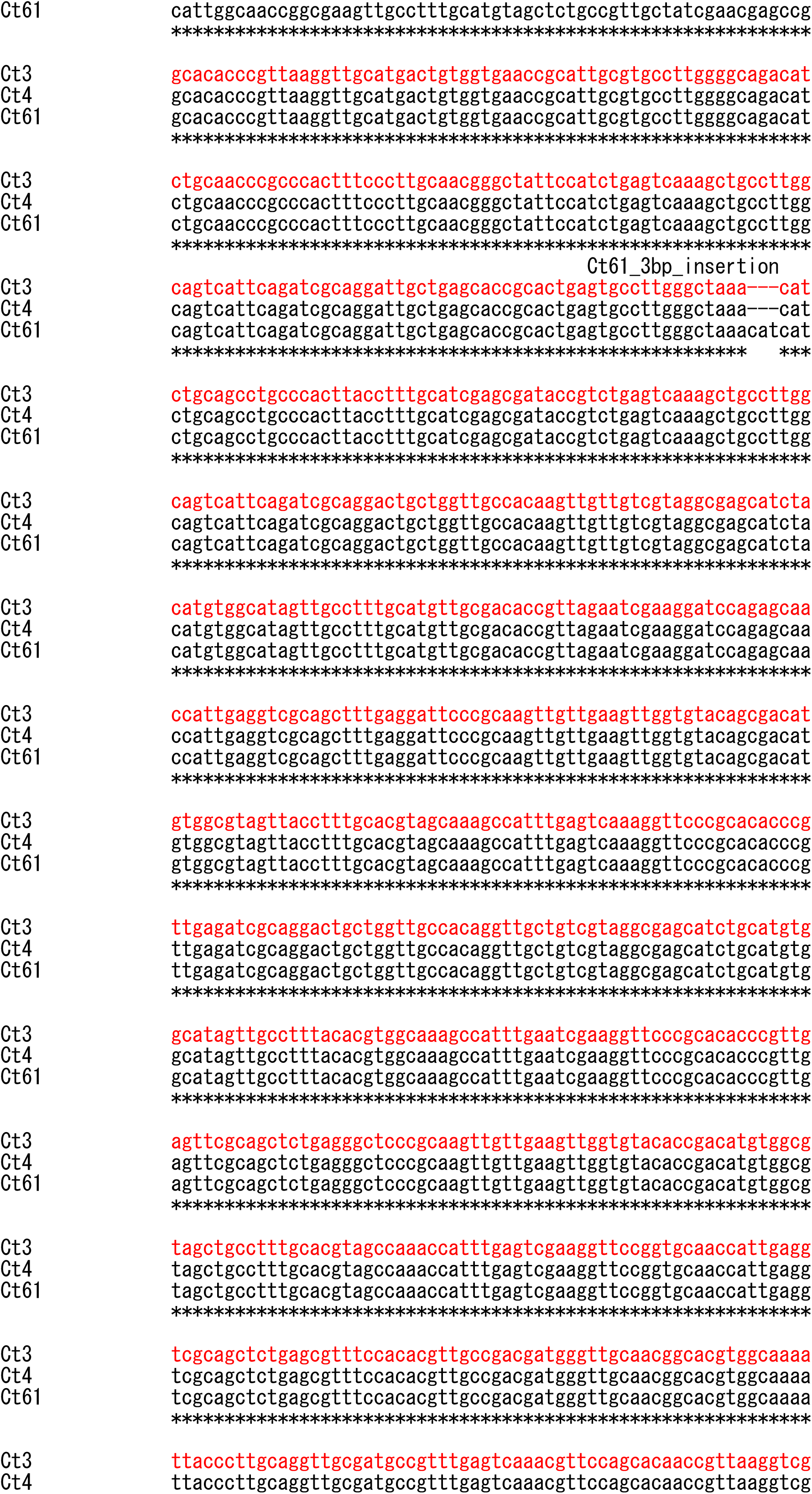
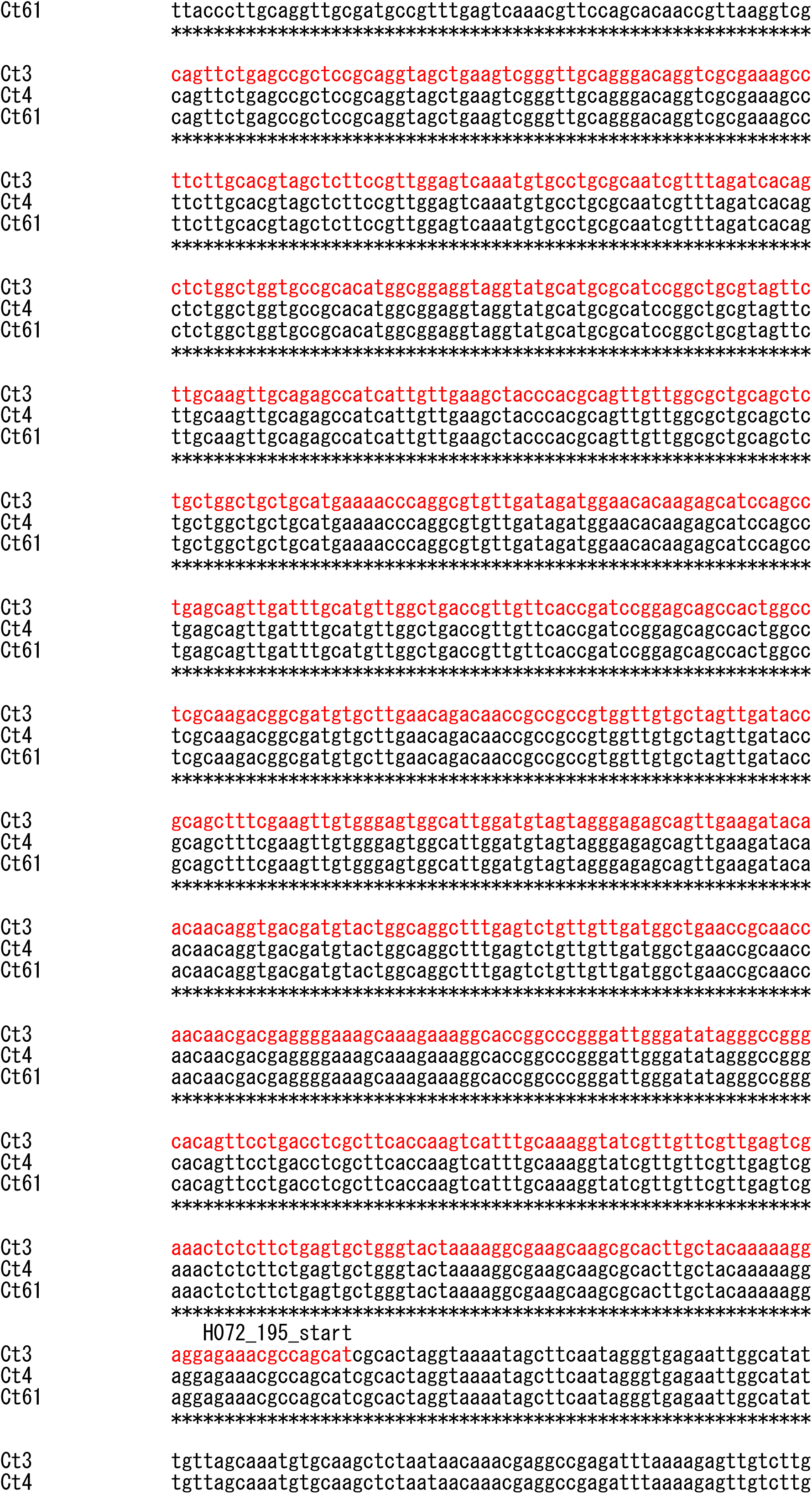
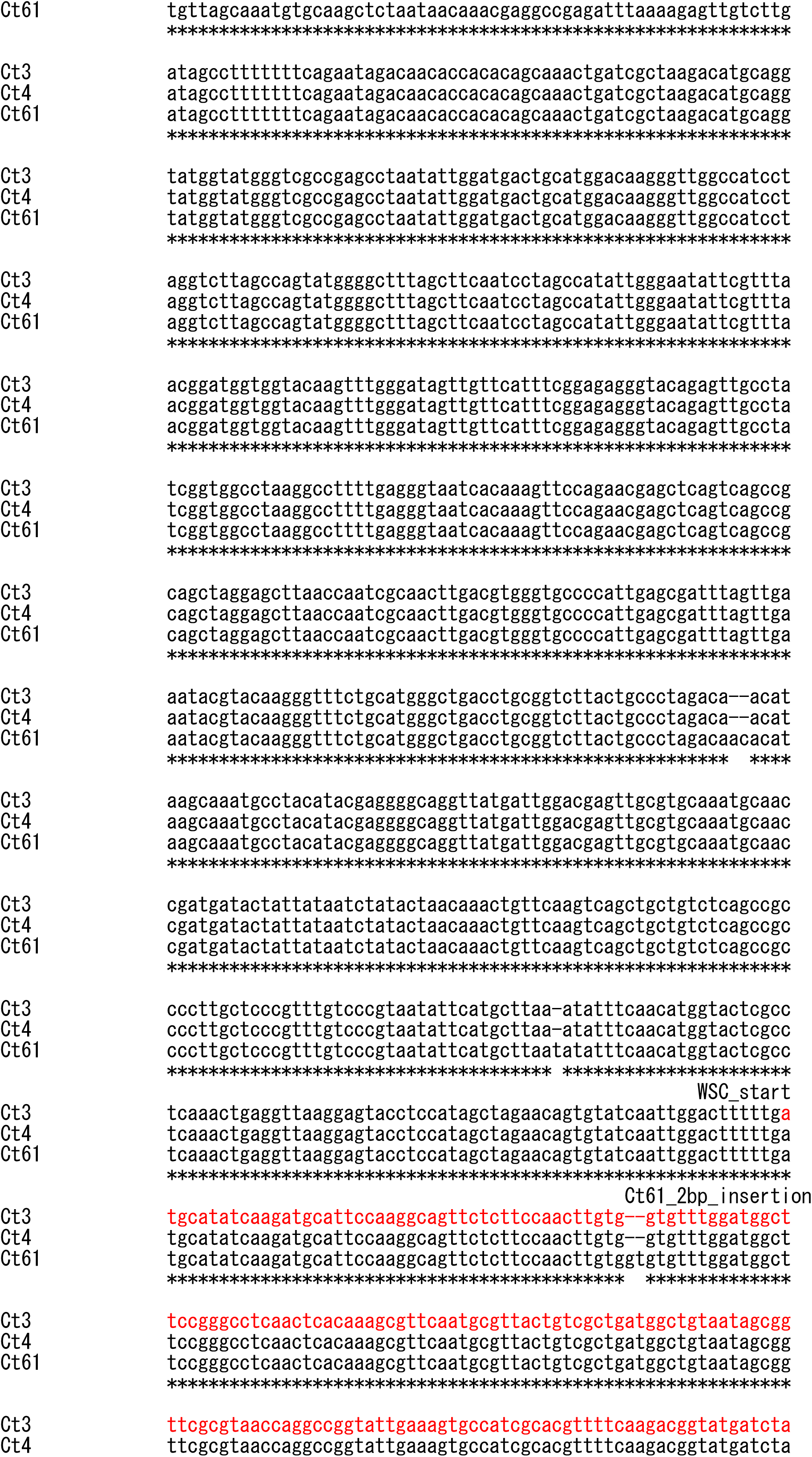
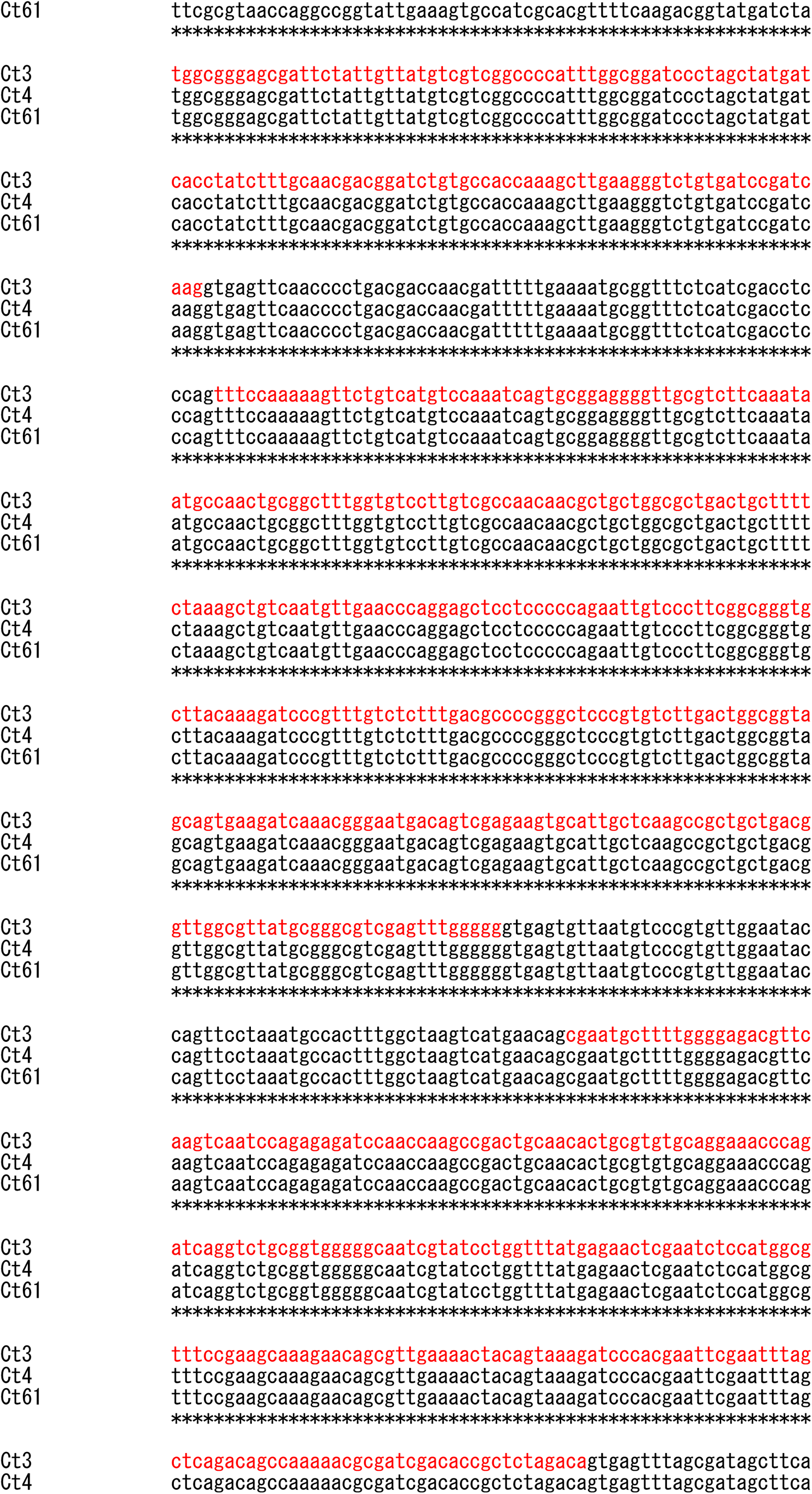
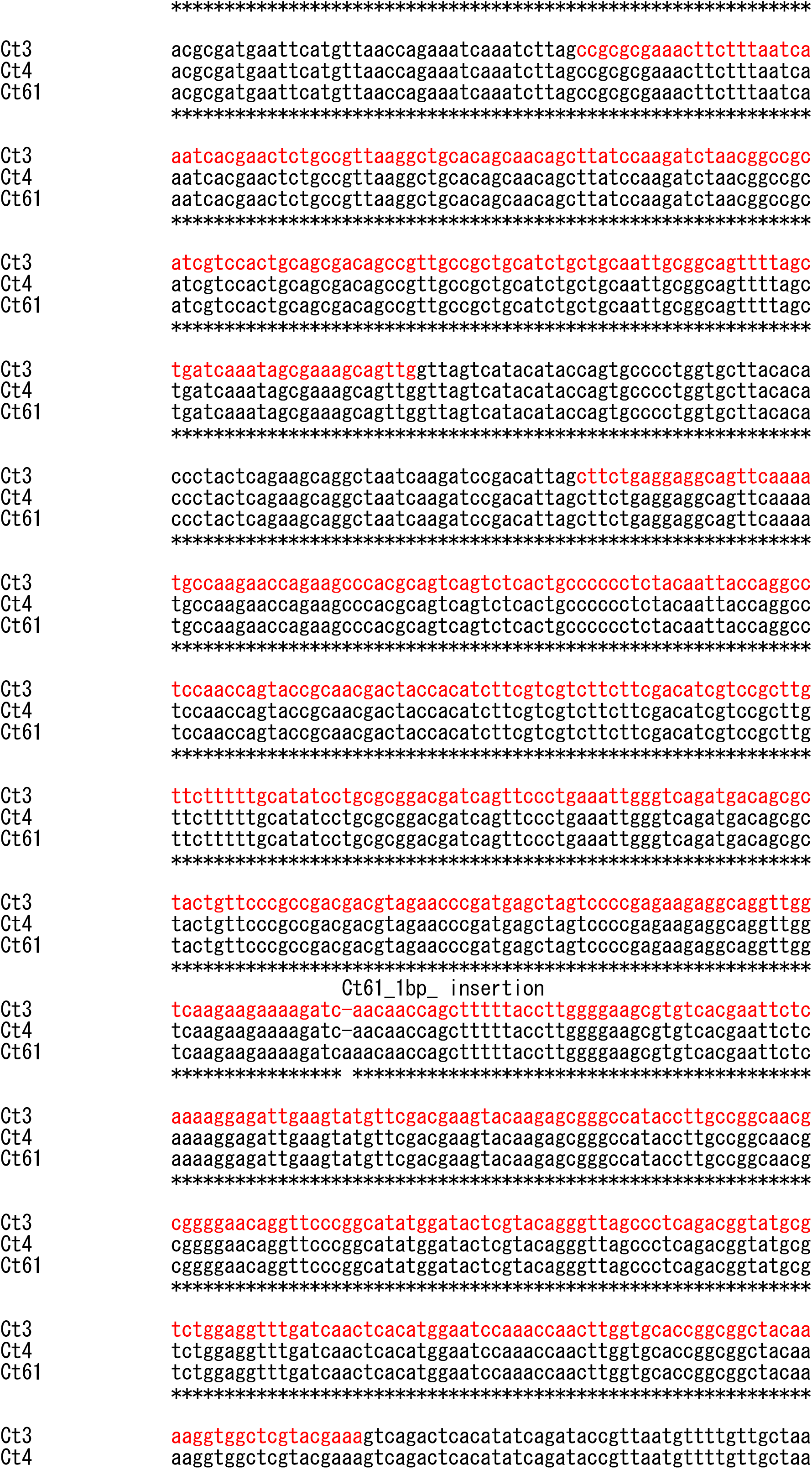
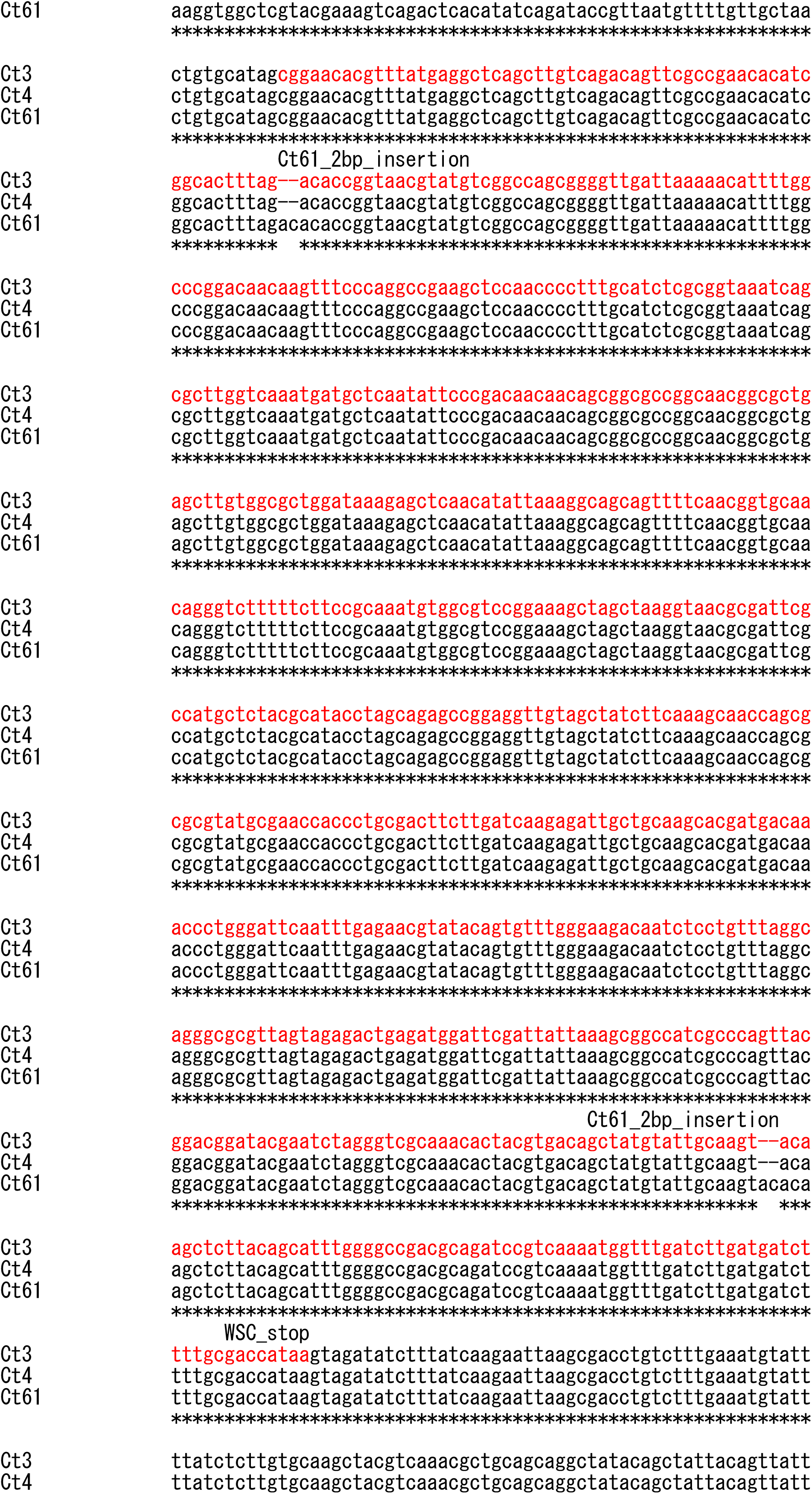
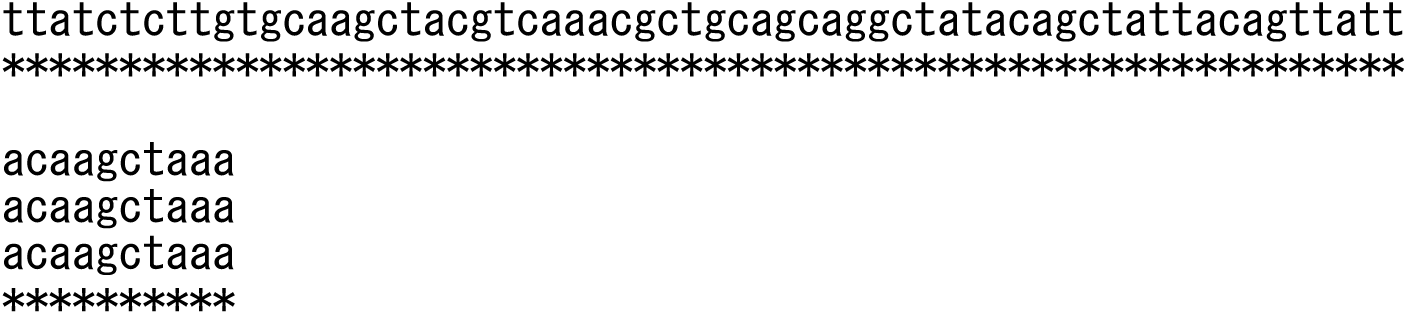
S3N. Alignment of 38k bp region including ABA and BOT clusters of *C. tofieldiae* strains. Sequences were aligned by MAFFT v7.471. Red letters indicate open reading frames of the genes.

## Notes

### Competing Interest Statement

The authors have declared no competing interest.

